# Electro-Fermentation of Grape Must via *Candida tropicalis* SY005: Accelerating Kinetics, Modulating Bio-chemical Pathway and Improving Bio-active Content

**DOI:** 10.64898/2026.07.25.740694

**Authors:** Suryansh Sharma, Sunakshi Gautam, Madhu Gaidher

## Abstract

This study investigates electro-fermentation *candida Tropicalis* SY005 to address fermentation kinetics limitation during grape must fermentation. In comparison with non-stimulated control sample, EF substantially enhanced sugar depletion, TSS drop by day 3 and generated a strongly reduced state (ORP -100 to -143mV). The oxidation-reduction shift enhanced cellular NAD^+^ regeneration, reducing total fermentation duration from 264 h to 72 h. GC-MS analysis showed pronounced major characteristic volatile compound confirming substantial metabolic pathway shifts in flavor of glycolytic flux. Moreover moderate electric field promoted cellular membrane electropermeabilization substantially promoting bioactive extraction.

## 1.0 Introduction

Electro-fermentation system (EFS) is a type of bioelectrochemical system (BES) that uses electrochemistry to control fermentation. In EFS, a carbohydrate or alcohol is fermented, and electrodes are used to provide either an additional source or sink for electrons (Moscoviz *et al*., 2016). The electric current plays a crucial role in facilitating the fermentation process under non-equilibrium conditions. The electric current influences both extracellular and intracellular redox potentials (ORP) at even low current densities, redox potential impacts the balance of NADH and NAD^+^ during anaerobic fermentation process (Choi *et al*., 2012; Speers *et al*., 2014; Xafenias *et al*., 2015). To modify the environmental redox potential, various chemicals with standard redox potentials higher or lower than those of common metabolic components are added to the fermentation broth. Some frequently employed reductants and oxidants for controlling extracellular redox potential include FeCl_3_, Na_2_S, potassium ferricyanide, dithiothreitol, cysteine, methyl viologen, neutral red, H_2_O_2_, and even directly added NADH and NAD+ (Snoep *et al*., 1991).

The balance between NAD+ and NADH, indicating oxidizing tendencies (NAD+ is an oxidizing agent that get reduced to NADH when it accepts electrons from reducing substrates during catabolic reactions like glycolysis which contribute significantly to the turnover of NAD+ and NADH. During glycolysis, glyceraldehyde 3-phosphate is converted to 1,3- bisphosphoglycerate by glyceraldehyde phosphate dehydrogenase, generating NADH. This NADH can be recycled back to NAD+ in two ways: through fermentation, where alcohol dehydrogenase converts acetaldehyde to ethanol using NADH via mitochondrial external NADH dehydrogenase (Murray *et al*., 2011).

There is ample of research reported by many researchers, such as Choi et al. (2014), they shows that altering the NAD+ /NADH in the yeast help to boost ethanol production, this alteration was achieved by using a modified yeast strain in which the glycerol was suppressed, they additionally overexpressed UTR1, a gene encoding ATP-NADH kinase by consuming ATP. This research confirmed that improving the NAD+ /NADH ratio and introducing the foreign GapN pathway could effectively replace glycerol’s role as an anaerobic energy sink, reducing glycerol production and enhancing ethanol yield. So logic behind to increase the ethanol production is To increasing glycolytic flux, as seen in anaerobic yeast fermentation process glycolysis converts NAD^+^→ NADH and glycolysis to continue, NADH must be oxidized back to NAD^+^, So anaerobic fermentation regenerates NAD^+^:-

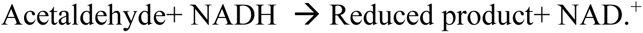

However existing methods to manipulate intracellular NADH/NAD^+^ ratio performing expensive and toxic redox mediator and deploying complex bio-electrochemical setups-faces practical limitation. Although electro-fermentation offers a sustainable stategy using solid state electrode as electron donor and Fe_3_O_4_ particles were act as redox mediator due to their mixed-valence iron state (fe^2+^/fe^3+^ redox couple) and high electron conductivity via electron hopping which allows reversible electron transfer between electrode and microbial metabolic pathway without cell toxicity (Skomurski *et al*., 2010). Its implementation for yeast derived grape must fermentation to concurrently speed up fermentation kinetics alter redox potential and enhance polyphenolic extraction during anaerobic fermentation process is predominantly underexplored. So our hypothesis is, if we providing extracellular electron via electrodes at low current densities will increases the availability of NAD^+^ = Higher glycolytic flux and we get more reduced metabolites . This experiment will be conducted using a wine fermentation model to test this hypothesis, allowing to analyze the ORP and Brix (°B) and GC-MS analysis to track the progress of the yeast metabolic shift towards reduced metabolites like ethanol, 2,3- butanediol and glycerin.

## 2.0 Material and Methods

The freeze dried yeast *Candida tropicalis* SY005 was purchased from the MTCC (Mohali). YPD medium (peptone 20 g/L, yeast extract 10 g/L and glucose 20 g/L) and Fe_3_O_4_ particles were obtained from the HI-MEDIA, LOBA chemicals and sigma aldrich. Grapes were procured from local market of Solan. DAHP, filtering agent and glass ware for storage of wine were purchased from International Scientific and Surgicals.

### 2.1. Wine preparation

Fresh local grape varieties were selected and thoroughly cleaned to remove debris. Crushing was performed as a primary unit operation for must extraction from the pulp. The must was then supplemented with 0.1% of diammonium hydrogen phosphate (DAHP) and 20g of glucose. Finally, the mixture was inoculated with starter culture of *Candida tropicalis* SY005, and fermentation was initiated at a controlled temperature of 25-30°C.

### 2.2. Fabrication of electro-fermentation cell

The fermentation cell consisted of an electrode assembly and inoculated wort contained in a 100 ml of graduated 3 neck round bottom borosilicate glass flask. The electrode assembly comprised a 25 cm² of two conductive carbon fiber cloth both the electrode were connected with specialized designed DC voltage power supply, to monitor the oxidation reduction potential and temperature parameters we placed a 2 in 1 orp + temperature meter were obtained from Ionix India limited.

**Figure 1.**
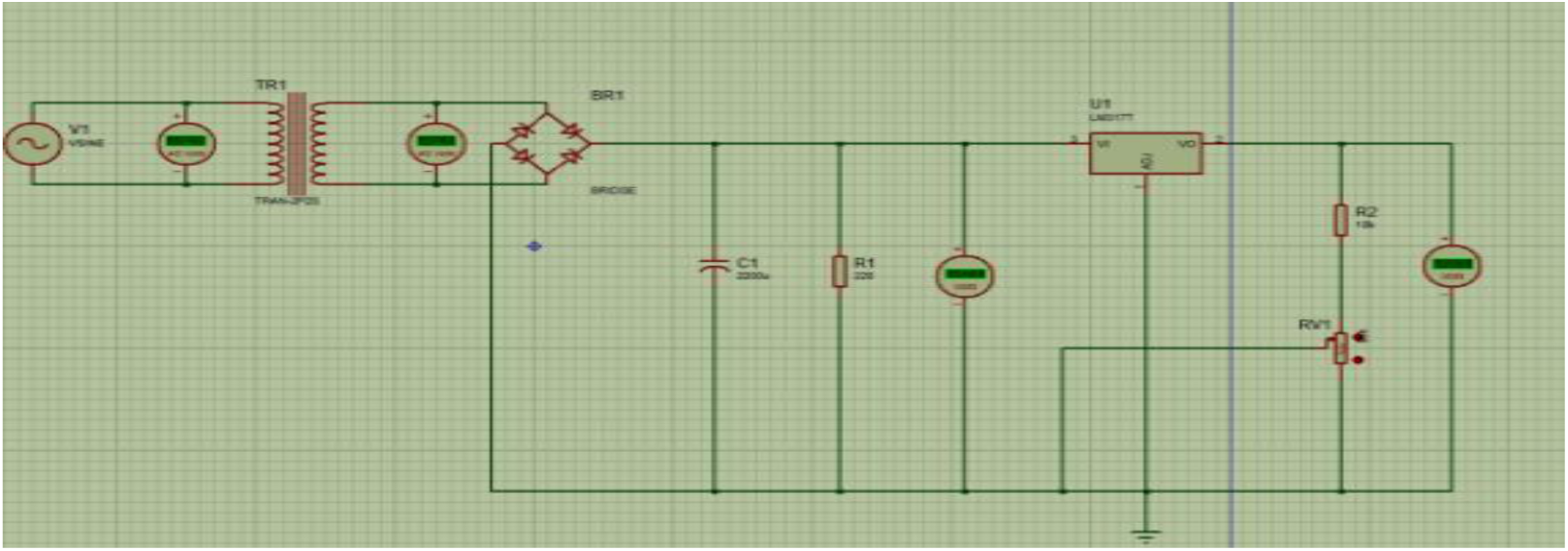
Illustrates the circuit design outlined in Proteus 8 CAD software, which used to regulates voltage and current vairations.

**Figure 2.**
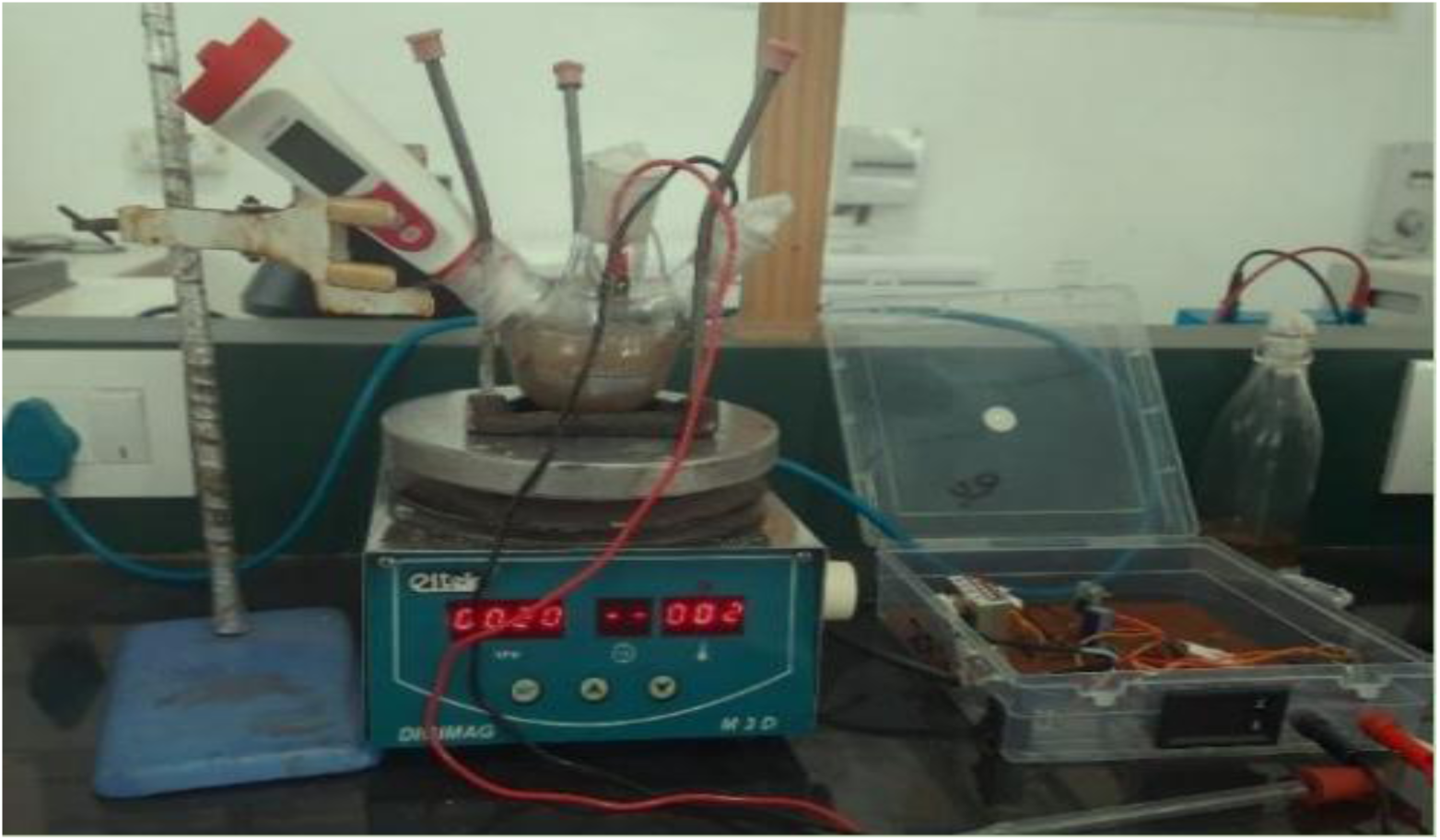
show dual Banana connector pins links the internal circuitry to two carbon fiber cloth electrode via crocodile clips. These electrode are positioned withi a 100 mL, three-neck borosilicate glass flask containing 84 mL of ferementation broth (dextrose+yeast + grape juice). To maintain anaerobic condition, the flask is sealed with cotton plug and it wrapped with paraffin.M.

### 2.3. Total soluble solids

Total Soluble Solids in grape juice and wine was determined by using hand held refractometer model ERMA RHB-32 ATC (Rangnna, 2007). The TSS range measured was 0-32%, expressed in degree units presented as °B.

### 2.4. Estimation of alcohol content

The alcohol content of treated wines is estimated using dichromate oxidation method (Zoeckleim *et al*., 1990), This procedure requires a preparation of specific reagent add potassium dichromate solution (0.0656M) in sulfuric acid (16.25mL) and for indicator solution add 0.050M ferrous sulphate and sulfate/o-phenanthroline (0.150M) in 25 mL of distilled water. For the actual titration a stock solution is first made by diluting the wine sample with distilled water in 1:10 ratio and add 12.5 mL of potassium dichromate solution in 5mL of wine that taken from the stock solution. After the reaction place the flask in water bath at 60-65°C for 30 minutes.

For Back titration Using ferrous ammonium sulphate solution in the Burette. A blank titration must be performed first until the solution turns green at which put 5 drops of Indicator solution are added, titration continue until colour change to brown. Finally the black titre and sample titre are recorded for calculation using the formula given below:-

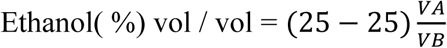

### 2.5. Total phenolic content

Phenolic content was measured using the following method: Prepare a gallic acid standard solution for a calibration curve. Mix the wine sample/standard with distilled water and folin-ciocalteu reagent. The mixture was then vortex rest for 5 minutes. Then add 1.5mL of 20% sodium carbonate solution. Mix well and rest for 2 hours in dark, finally measured the abosrbance at 750nm and calculate phenolic content using standard curve equation (Siddiqui *et al*. 2017).

### 2.6. Total flavonoid content

Flavonoid was measured using the following method: Prepare a rutin standard solution for a calibration curve. The aqueous extract was mixed with 0.3 mL of 5% of sodium nitrite solution after 5 minutes 0.3 ml of 10% AlCl_3_, after 6 minutes 2 mL of 1M NaOH was added and finally measured the abosrbance at 510nm and calculate flavonoid content using standard curve equation (Meda *et al*., 2005).

### 2.7. Anthocyanin Content

Anthocyanin content was measured using the following method: mix 95% ethanol solution and 1.5N HCL solution in the 85:15 ratio this will create the ethanolic HCL, after that 10 ml of wine sample mix with ethanolic HCL and mixture was sealed for overnight at 4°C after incubation period is over, mixture was transferred to dark location for 2 hours and absorbance was taken at 535 nm using spectrophotometry (Rangnna, 2007) and calculate using given formula:

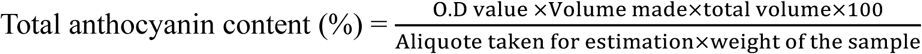

### 2.8. FTIR Spectroscopy analysis

FTIR spectroscopy was used to determine the functional groups changes induced by electro-fermentation. Spectra were recorded using a Thermo Fisher Scientific Nicolet iS5 Fourier Transform InfraRed Spectrometer, Thermo Fisher Scientific Inc., Waltham, Massachusetts, USA. In the spectral range 4000–400 cm⁻¹.

### 2.9. FE-SEM analysis

FE-SEM was used to investigates yeast cellular morphology prior to electro-fermentation. Samples were fixed desiccated using a graded ethanol gradient and gold-sputter coated prior to imaging. Micrographs were captured utilizing a FE-SEM model zeiss sigma 300, JEOL JSM- 7610F, Hitachi SU8010 under an accelerating voltage of (5-15 kv). Elemental characterization was carried out via Energy Dispersive Spectroscopy (EDS) integrated to the same instrument.

### 2.10. GC-MS analysis

Sample volatile were collected using headspace solid-phase Microextraction (HS-SPME). In summary 5ml of fermented sample, 1.5 g NaCl and 10μl internal standard (2-octanol, 50mg/L) were enclosed in a 20 mL headspace vial. After equilibration at 40°C for 10 min (250 rpm), and SPME fiber (50/30μm DVB/CAR/PDMS) was exposed to the head space for 30 min. at 40°C to facilitate volatile sorption to thermal desorption in the GC inlet.

## 3.0 Result and Discussion

### 3.1. Kinetics of sugar consumption oxidation reduction potential modulation, fermentation Kinetics and ethanol yield

Electro fermentation experiment were conducted using *Candida tropicalis* SY005 yeast under standard condition, including a pH of 4.0 and a temperature 30°C. The fermentation system was supplemented with grape must and glucose, while the external voltage was maintained to 1.3-1.7V with current range of 0.12-0.19mA.

**Table 1.**
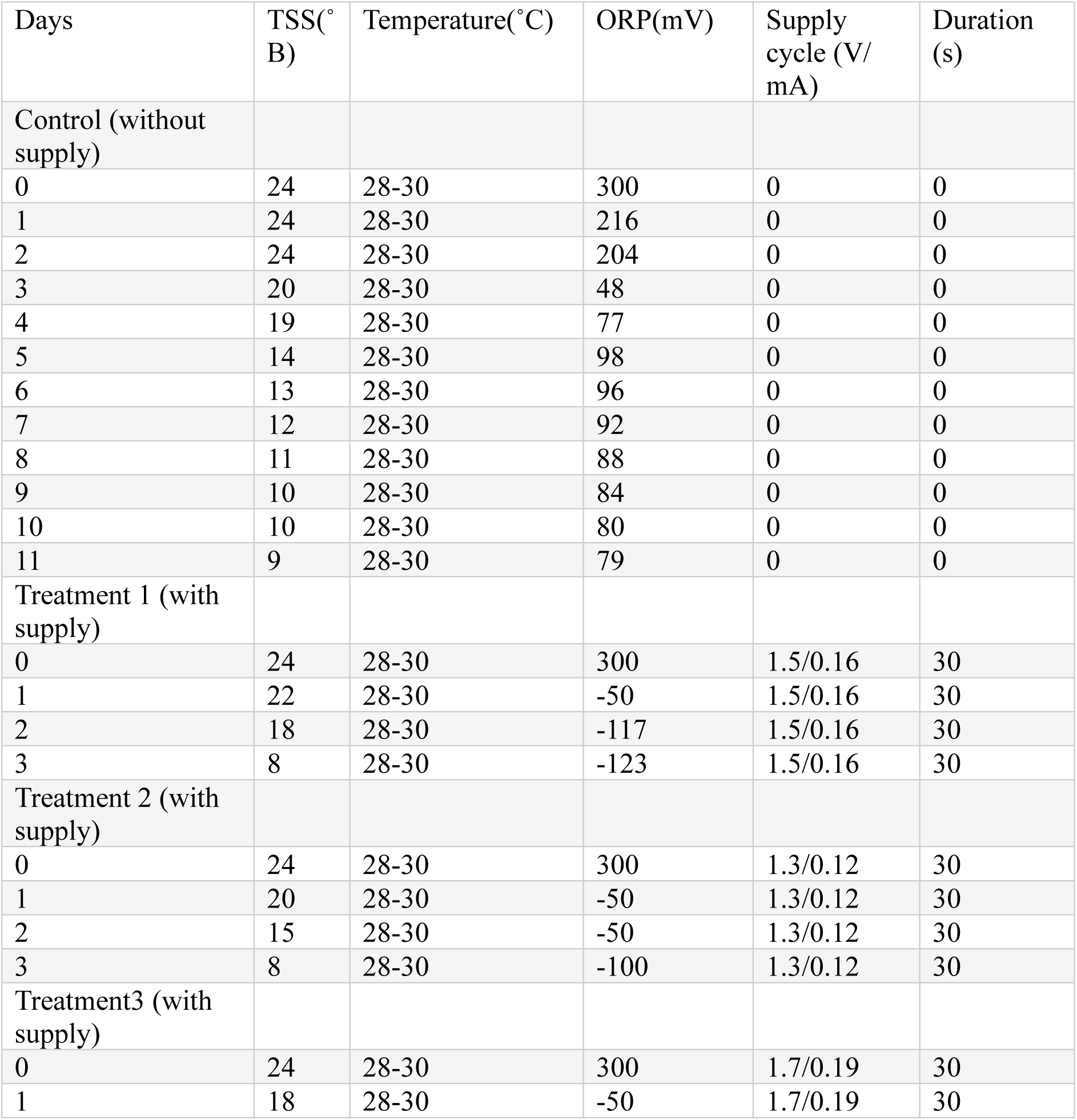
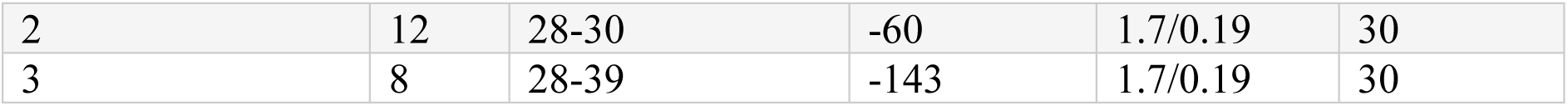
Effect of electro-fermentation on total soluble solids (TSS), temperature and oxidation-reduction potential (ORP) during grape must fermentation.

| Days | TSS(°B) | Temperature(°C) | ORP(mV) | Supply cycle (V/<br>mA) | Duration (s) |
| --- | --- | --- | --- | --- | --- |
| Control (without supply) |  |  |  |  |  |
| 0 | 24 | 28-30 | 300 | 0 | 0 |
| 1 | 24 | 28-30 | 216 | 0 | 0 |
| 2 | 24 | 28-30 | 204 | 0 | 0 |
| 3 | 20 | 28-30 | 48 | 0 | 0 |
| 4 | 19 | 28-30 | 77 | 0 | 0 |
| 5 | 14 | 28-30 | 98 | 0 | 0 |
| 6 | 13 | 28-30 | 96 | 0 | 0 |
| 7 | 12 | 28-30 | 92 | 0 | 0 |
| 8 | 11 | 28-30 | 88 | 0 | 0 |
| 9 | 10 | 28-30 | 84 | 0 | 0 |
| 10 | 10 | 28-30 | 80 | 0 | 0 |
| 11 | 9 | 28-30 | 79 | 0 | 0 |
| Treatment 1 (with supply) |  |  |  |  |  |
| 0 | 24 | 28-30 | 300 | 1.5/0.16 | 30 |
| 1 | 22 | 28-30 | -50 | 1.5/0.16 | 30 |
| 2 | 18 | 28-30 | -117 | 1.5/0.16 | 30 |
| 3 | 8 | 28-30 | -123 | 1.5/0.16 | 30 |
| Treatment 2 (with supply) |  |  |  |  |  |
| 0 | 24 | 28-30 | 300 | 1.3/0.12 | 30 |
| 1 | 20 | 28-30 | -50 | 1.3/0.12 | 30 |
| 2 | 15 | 28-30 | -50 | 1.3/0.12 | 30 |
| 3 | 8 | 28-30 | -100 | 1.3/0.12 | 30 |
| Treatment3 (with supply) |  |  |  |  |  |
| 0 | 24 | 28-30 | 300 | 1.7/0.19 | 30 |
| 1 | 18 | 28-30 | -50 | 1.7/0.19 | 30 |
| 2 | 12 | 28-30 | -60 | 1.7/0.19 | 30 |
| 3 | 8 | 28-39 | -143 | 1.7/0.19 | 30 |

The Table.1 provides the data of various parameters of Electro Fermentation it distinguishes between the control and the group based on the application of the electrical current voltage range. As fermentation control was started with high orp +300 mV at 0.13mA, 9-11 days. It quickly drops around + 80 to 180mV on the 11^th^ day but in the treatment 1,2 and 3 started with high orp +300mV but it reduced drastically at a third day at -100, -117 and -143mV. The ORP result is in the negative that means environment is more reduced. In control TSS reduced to 9°Brix at 11^th^day and in a treatment 1,2 and 3 reduced by 8°Brix at 3^rd^ day. This parameter shows Sugar reduced in electro-fermentation is lot faster in treatment 1,2 and 3. Overall, the rapid decline in TSS accompanied by strongly negative ORP values demonstrated that electro- stimulation creates a highly reducing intracellular environment, significantly accelerating yeast metabolic activity and substrate consumption (Arbter *et al*., 2019).

According to Table 2, ethanol production was significant enhanced, reaching 13.00±0.40% in Treatment 3 (72 h) compared to 7.00±0.26% in the control (264 h). One-way ANOVA, F(3,8)=211.97, P<0.001, tukey’s HSD p<0.05. This shows electro fermentation lowers the redox potential of the medium which creates a reduced environment. This increase the intracellular yeast NADH/NAD^+^ ratio, Which favours ethanol formation as a primary electron sink, resulting in higher ethanol yield. This proved our hypothesis that if we providing the extracellular electron by electrodes and lower the current density will increase the glycolytic flux. The observed ORP from the treatment 1, 2 and 3 -100, -117 and -143mV respectively, and was significantly more positive than standard NADH/NAD^+^ potential -320mV Nelson et al., (2017) indicating redox environment favouring NADH oxidation and enhanced NAD^+^ regeneration (Moscoviz *et al*., 2016). Thus, the lower ORP induced by electro-fermentation not only facilitates cellular redox balance for faster NAD^+^ replenishment but significantly shortens fermentation time simultaneously optimizing ethanol yield.

**Figure 3.**
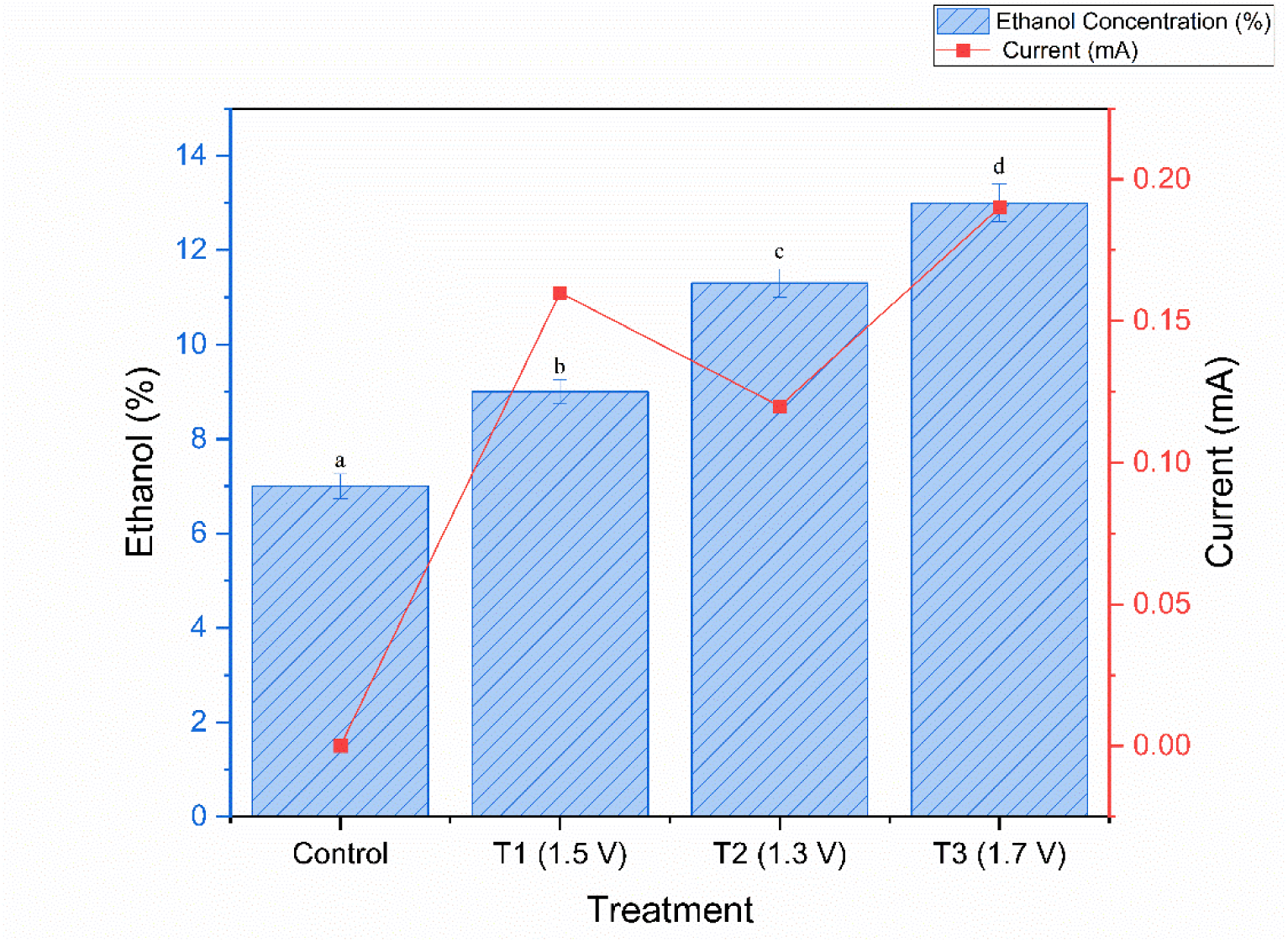
Ethanol concentration (bars, left axis) and current (line, right axis) across treatments. Voltage for each treatment is indicated on the X-axis. Error bars represent SD (n=3). Different superscript letters indicate significant differences (Tukey’s HSD,p<0.05)

**Table 2.** Effect of applied electrical potential and current on ethanol concentration and fermentation time during electro-fermentation.

| Treatments | Supply cycle V/mA | Supply cycle (mili ampere) | Ethanol concentration (%) | Fermentation time (Hours) |
| --- | --- | --- | --- | --- |
| Control | Without electric field | Without electric field | 7.00±0.26<sup>a</sup> | 264 |
| Treatment 1 | 1.5 | 0.16 | 9.00±0.26<sup>b</sup> | 72 |
| Treatment 2 | 1.3 | 0.12 | 11.3±0.30<sup>c</sup> | 72 |
| Treatment 3 | 1.7 | 0.19 | 13±0.40<sup>d</sup> | 72 |

Note: Values are mean±SD (n=3). Means bearing different superscript letters (a,b,c,d) within the same column differ significantly from one another (one-way ANOVA followed by Tukey's HSD post hoc test, p<(0.05)

### 3.2 Effect of Electro-Fermentation on Functional Group Composition (FTIR analysis)

In the higher wavenumber region a broadened absorbance spectral band corresponding to hydroxyl (-OH) stretching arising from water molecules, alcohol group and phenolic group compounds was detected at 3357 cm^-1^ in the control sample (84.81% T). In the electro-treated samples this band shifted towards lower wavenumber with a decreased in % transmittance being observed at 3260 cm^-1^ in WT1 (44.84%) , 3268 cm^-1^ in WT2 (44.81%), 3335 cm^-1^ in WT3(44.10%). The decreased in transmittance within this region indicates increased molecule to molecule hydrogen bonding and higher overall hydroxyl concentration within the electro-fermentation sample. In the aliphatic spectral region (2800-3300 cm^-1^) asymmetric and symmetric -CH_3_ and CH_2_ stretching vibration was observed at 2999 cm^-1^ for the control (95.78% T), WT1(86.08% T) and WT2 (86.38% T), whereas the band shifted to 2992 cm^-1^ in WT3 (83.75% T). The decreased transmittance% within the absorption band indicates an enhanced accumulation of aliphatic ethyl groups derived from ethanol and volatile secondary metabolites. The band located at 1641 cm^-1^ assigned to carbonyl (C=O) stretching of organic acids/esters overlapping with water bending, demonstrated a substantial decreased in transmittance from 90.62% T in the control to 67.13% T, in WT1, (67.21% T) in WT2 and 65.84% T in WT3. Most significantly the dominant characteristics band for ethanol concentration assigned to C-O and C-C stretching vibration was indentifies at 1045 cm^-1^. The control sample exhibited a high % transmittance of 94.62% T at the wavenumber indicating low ethanol production. However electro-fermentation led to clear decline of tranmittance at 1045 cm^-1^ to 82.95% T in WT1 and 83.95%T in WT2 reaching the lowest transmittance of 73.60% in WT3. Additionally primary/secondary alcohol related absorption bands located at 1082 cm^-1^ (C-O) stretching and 806 cm^-1^ (C-H) bending showed a comparable reduction in trasnmittance following the order : Control > WT2= WT1>WT3.

**Figure 4.**
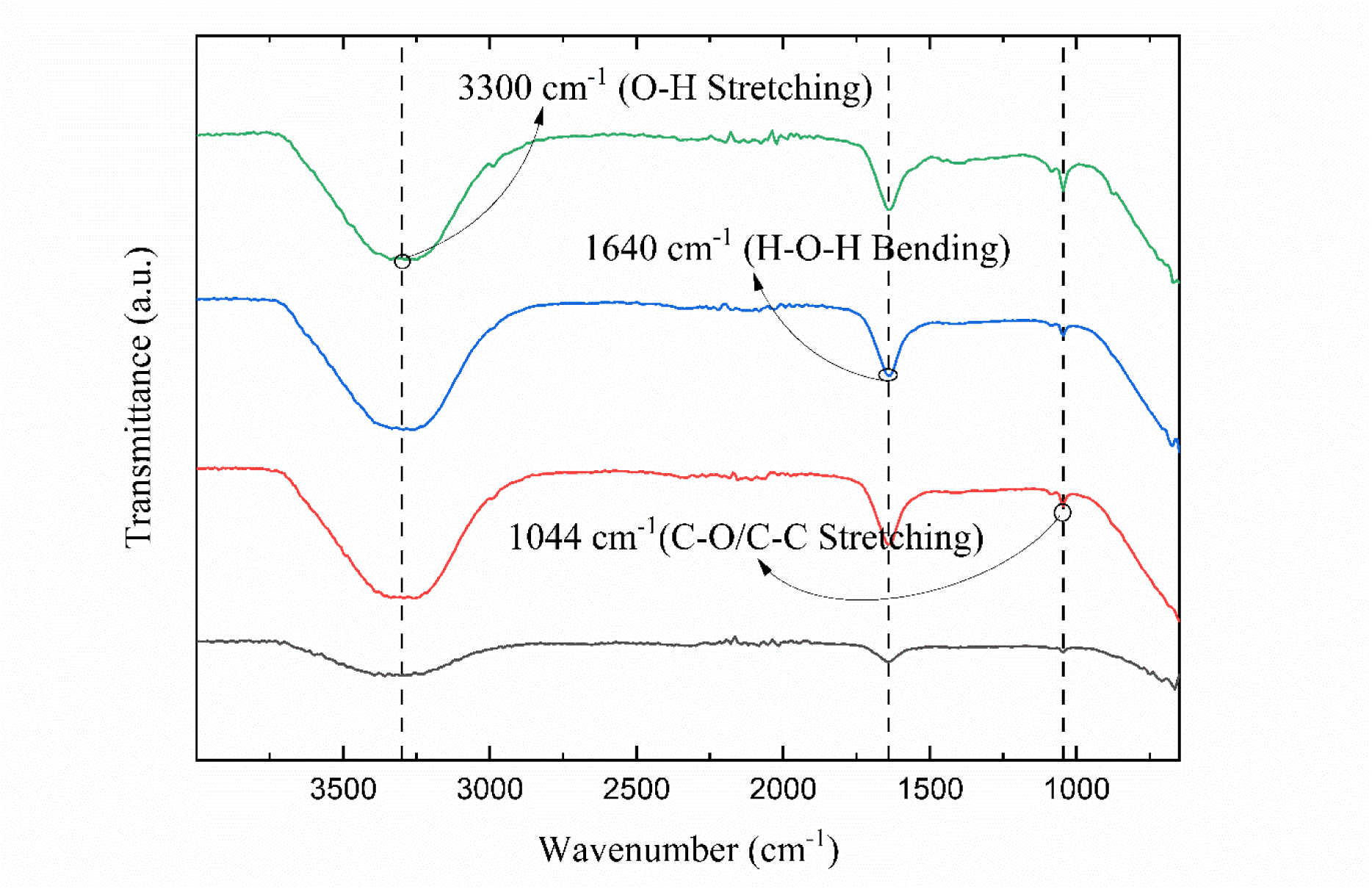
FTIR spectrum plot of the control wine (WC1), and electrofermentation samples (WT1,WT2,WT3) recorded over the 4000-600 cm^-1^ region

**Table 3.** Major FTIR spectral peak assignments and comparative transmittance values of control wine electro-fermentation treatments.

| Wavenum<br>ber ($\text{cm}^{-1}$) | Functional Group | Vibrational<br>Mode | Control<br>(WC1) | WT1 | WT2 | WT3 | Key<br>Indicator |
| --- | --- | --- | --- | --- | --- | --- | --- |
| 3357-3260 | Hydroxyl group (-OH) | O-H stretching | 84.81%<br>(0.072) | 44.84%<br>(0.348) | 44.81%<br>(0.349) | 45.10%<br>(0.346) | Water,<br>alcohols,<br>phenolics |
| 2999-2992 | Aliphatic Carbon (-$\text{CH}_3$, -$\text{CH}_2$-) | C-H asymmetric stretching | 95.78%<br>(0.019) | 86.08%<br>(0.065) | 86.38%<br>(0.064) | 83.75%<br>(0.077) | Higher<br>Alcohols |
| 1641 | Carbonyl/Hydroxyl | $\text{C}=\text{O}$ stretching/H<br>-O-H bending | 90.62%<br>(0.043) | 67.13%<br>(0.173) | 67.21%<br>(0.173) | 65.84%<br>(0.182) | Organic<br>acids,<br>aromatic<br>esters |
| 1045 | Primary alcohol | C-O and C-C-stretching | 94.62%<br>(0.024) | 82.95%<br>(0.081) | 83.64%<br>(0.078) | 73.60%<br>(0.133) | Ethanol<br>(Primary peak) |
| 1082 | polyol | C-O stretching | 96.23 %<br>(0.017) | 87.92%<br>(0.056) | 87.96%<br>(0.056) | 81.70%<br>(0.088) | Ethanol/Glycerol<br>skeletal |
| 806 | Aliphatic C-H | C-H out of plane deformation | 89.85 %<br>(0.046) | 63.42 %<br>(0.198) | 63.31%<br>(0.199) | 61.12%<br>(0.214) | Volatile aromatic compound |

### 3.3. Effect of Electro-Fermentation on Volatile Compound Profile (GC-MS analysis)

GC-MS characterization of the electro-fermentation samples confirmed give major metabolites 2,3-butanediol (R_t_ 4.54 min, 13.05%), glycerin (R_t_ 9.28 min, 14.33%), alpha-D-Glucopyranose derivative (R_t_ 18.34 min, 2.04%), hexadecanopic acid, methyl ester (R_t_ 23.34 min, 4.15%) and heptadecanoic acid (R_t_ 26.06 min, 1.77%) with compound assignments confirmed by comparing with mass spectral database together with published literature retention indices (LRI_lit_) showed in Table5. The generated GC-MS spectrum demonstrated a evident accumulation of strongly reduced metabolic end-product supporting the hypothesis that cathode-derived electron influx promotes intracellular redox homeostasis (2e^-^ + 2H^+^ NAD^+^ →NADH). As shown in Table5, (2,3-butanediol) reflects an active cellular response to an altered intracellular redox state, the enzyme-catalyzed biochemical transformation pyruvate via acetoin reductase is obligatorily NADH dependent pathway. Formation of 2,3-butanediol and glycerin supports that the cathode-derived electrical potential supplied surplus reducing electrons, which were efficiently utilized and regenerate NADH back to NAD^+^ under Cathodic polarization (Medina *et al*., 2010;Tefft *et al*., 2019). This metabolic homeostasis altering bio- chemical flux towards reduced products. The detection of long-chain fatty acid and their corresponding methyl ester (Hexadecanoic acid and Heptadecanoic acid) align with well documented lipid producing and metabolic active lipid metabolic machinery of the *candida tropicalis* which can channel substantial metabolic flux towards cellular lipid deposition (Chattopadhyay *et al*., 2020)

**Figure 5.**
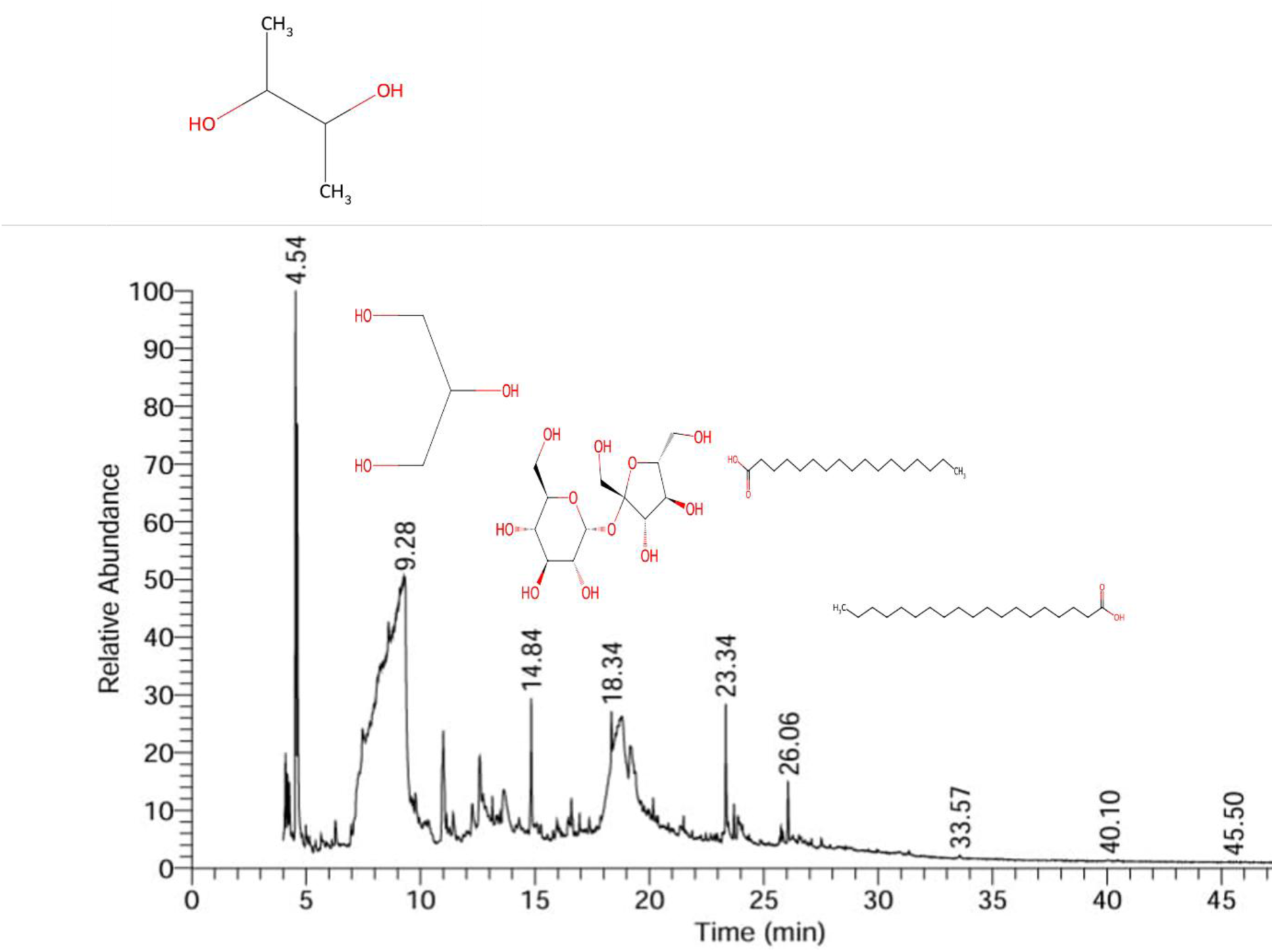
Gas Chromatography-Mass Spectrometry (GC-MS) total ion chromatogram (TIC) of the electro-fermentation sample with annotated chemical structure above the enhanced peaks with respect to retention time.

**Table 4.** GC-MS Identification Literature Retention Index, Relative Peak Areas Percentage and Functional Roles of metabolites in the electro fermentation sample.

| Retention Time (min.) | Compound Name | Molecular Formula | Peak Area(%) | $LRI_{lit}$ (NIST) | Probability | Functional Role |
| --- | --- | --- | --- | --- | --- | --- |
| 4.54 | 2,3-Butanediol | $C_4H_{10}O_2$ | 13.05% | 1494(Carbowax 20M) | 56.19 | Redox sink/Indicator of high NADH/$NAD^+$ |
| 9.28 | Glycerin | $C_3H_8O_3$ | 14.33% | 2322 (DB-Wax) | 91.17 | Polyol/Sugar Alcohol |
| 18.34 | α-D<br>Glucopyranose | C<sub>12</sub>H<sub>22</sub>O<sub>11</sub> | 2.04% | N/A* | 14.38 | Substrate<br>Intermedi<br>ate<br>/Sugar |
| 23.34 | Hexadecanoic<br>acid, methyl ester | C<sub>17</sub>H<sub>34</sub>O<sub>2</sub> | 4.15 | 1904.1 (OV-101) | 64.25 | Fatty<br>Acid<br>Ester |
| 26.06 | Heptadecanoic<br>acid | C<sub>19</sub>H<sub>38</sub>O<sub>2</sub> | 1.77 | 2074 (SE-30) | 34.13 | LongCha<br>in Fatty<br>Acid<br>Ester |

### 3.4. Effect of electro-fermentation on Surface Morphology (FE-SEM analysis)

FE-SEM coupled with Energy Dispersive Spectroscopy (EDS) was performed to characterize the morphological interaction between the yeast cell wall and Fe_3_O_4_ particles. Prior to electro-fermentation the yeast cell wall displayed a smooth uniform surface morphology (Figure 6A). Following electro-fermentation in the presence of Fe_3_O_4_ clear evident micro-crystalline aggregates were detected incorporated into the yeast cell biomass matrix (Figure 6B). Chemical element point quantification by EDS (Table 5) verified that the crystalline particle were mainly composed of iron (Fe) and oxygen (O) confirming the deposition of Fe_3_O_4_ particles. Moreover surface characterization of the surrounding cell wall revealed a substantial iron content of 17.86 wt% while the organic biomass matrix showed major elemental signal for biologically derived carbon, oxygen, phosphorus and potassium. The direct physical association between the Fe_3_O_4_ particle and the yeast cell wall indicate the establishment of a bio-inorganic contact zone. This electron-conducting Fe_3_O_4_ interface potentially act as extracellular electron mediator established across the cell wall to maintain the cellular redox state NADH/NAD^+^ (Luo *et al*., 2022), and promote metabolic flux into reduced products like ethanol, 2,3butanediol and glycerin.

**Figure 6.**
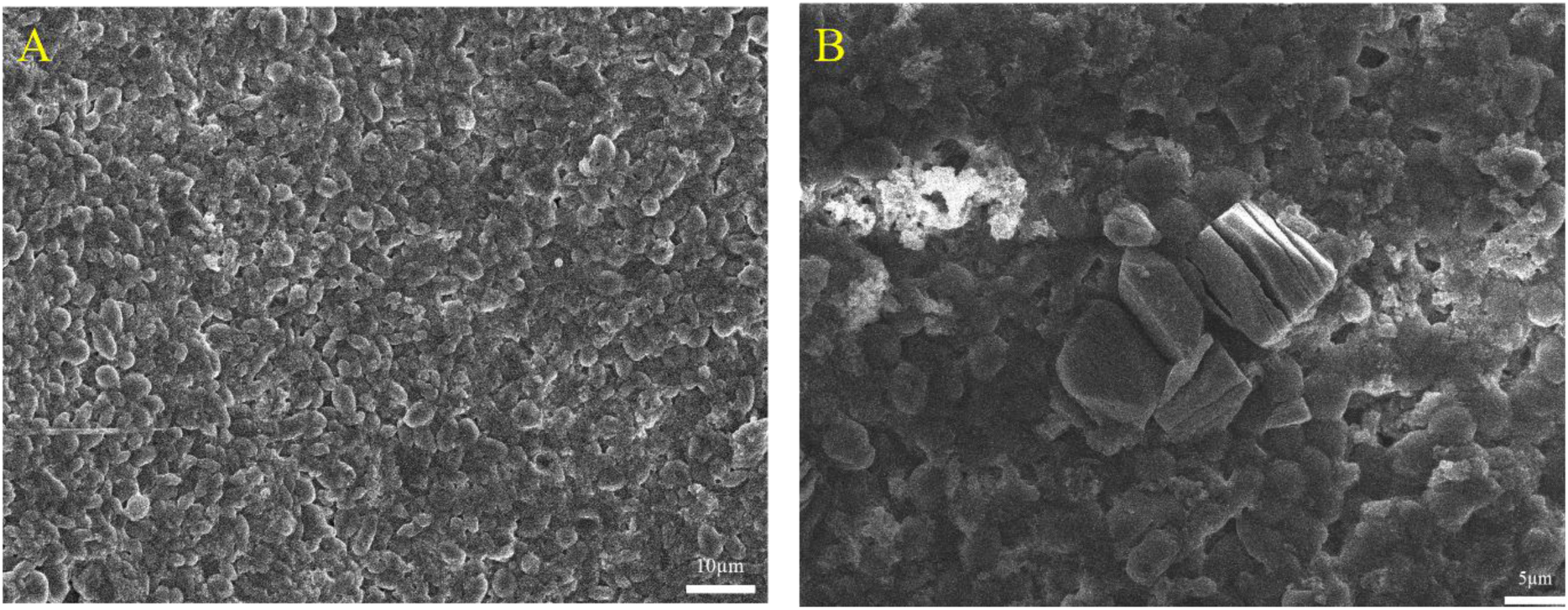
FE-SEM micrographs illustrating yeast cellular surface morphology (A) before electro-fermentation showing smooth uniform biomass distribution and (B) after electrofermentation in the presence of Fe_3_O_4_ particles showing evident microcrystalline deposits incorporated into the yeast cell wall.

**Figure 7.**
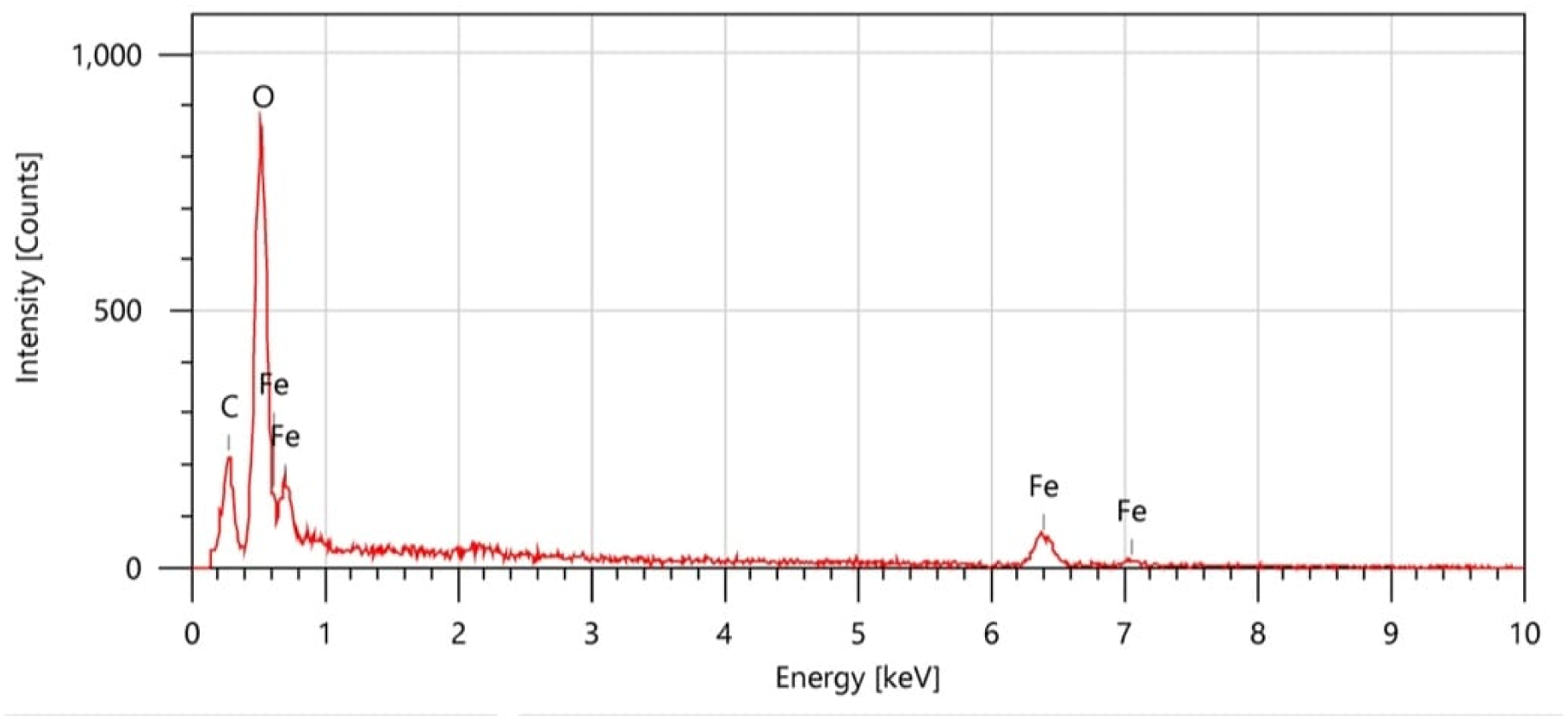
EDS spectrum obtained from the microcrystalline deposit located on the yeast cell wall after electro-fermentation illustrating elemental peak corresponding to iron (Fe,6.4 KeV and 7.0 KeV), Oxygen (O, 0.5KeV) and Carbon (C, 0.27 KeV).

**Table 5.** Elemental composition of yeast cell biomass and crystalline precipitates after electro-fermentation.

| Spectrum ID | Element | Mass (%) | Atom (%) |
| --- | --- | --- | --- |
| Crystalline precipitate | Carbon (C) | 12.46±0.18 | 23.92±0.34 |
|  | Oxygen (O) | 38.79±0.44 | 55.94±0.64 |
|  | Iron (Fe) | 48.75±1.88 | 58.99±0.44 |
| Biomass background matrix | Carbon (C) | 50.04±0.37 | 56.99±0.44 |
|  | Oxygen (O) | 43.39±0.78 | 38.40±0.69 |
|  | Phosphorus (P) | 2.42±0.21 | 1.11±0.12 |
|  | Potassium (K) | 4.15±0.34 | 1.50±0.12 |
| Cell wall surface deposit | Carbon (C) | 23.80±0.24 | 34.33±0.35 |
|  | Oxygen (O) | 53.64±0.64 | 58.10±0.70 |
|  | Calcium (Ca) | 4.70±0.36 | 2.03±0.15 |
|  | Iron (Fe) | 17.86±1.35 | 5.54±0.42 |

### 3.5. Effect of Electro-fermentation on Bio-active Compound Content

To understand the effect of electro-fermentation on bioactive compound release, Total phenolic content (TPC), Total flavonoid content (TFC) and Total anthocyanin content (TAC) were quantified across all treatments groups. One-way ANOVA reveals that all three physicochemical parameters increased significantly across all treatments compared to raw and control samples, TPC:-F(4,10)=97.66,p<(0.001), TFC:-F(4,10)=25.42,p<(0.001), TAC:-F(4,10)=205.12,p<(0.001), TPC increased progressively from 28.50 ±1.15 mg GAE/100mL, In the raw sample in the raw sample to 30.03±1.08 mg GAE/100mL, in the control and further to 42.00±1.66, 43.63±1.70 and 49.64±2.10 mgGAE/100mL in treatment 1,2,and 3 respectively. A similar trend was observed for TFC, 30.03±1.19mg RUE/100mL (raw) and 32.27±1.14 mg RUE/100mL (control) to 34.47±1.23, 35.17±1.07 and 39.18±1.07 mgRUE/100mL across treatment 1-3. TAC showed the most significant increase from, 20.03±1.06mg/100mL (raw) and 32.63±1.42mg/100mL (control) to 43.07±1.63,47.97±1.70 and 53.87±2.10 mg/100mL, in treatments 1-3 respectively. Tukey’s HSD post hoc test (p<0.05) confirmed that Treatment 3 differed significantly in all three parameters, while TAC demonstrated complete distinction across all five treatment groups, reflecting robust dose-dependent response to electro-fermentation. This pattern indicates the impact of electro-fermentation treatment on cell wall structure and polyphenol release. It may be possible that small induced electroporation of grape skin and pulp cell membrane structure altering cellular compartmentation and improving the release of bound phenolic, flavonoid and anthocyanin compounds, that would otherwise remain in bound form to cell wall polysaccharides matrix during traditional fermentation process (Donsi *et al*., 2010).

**Table 6.** Effects of electro-fermentation on bio-active content in wine sample.

| Treatment (V/mA) | Total Phenolic Content (mg GAE/100mL) | Total Flavonoid Content (mg RUE/100mL) | Total Anthocyanin Content (mg/100mL) |
| --- | --- | --- | --- |
| Raw (Red grape juice) | 28.5±1.15<sup>a</sup> | 30.03±1.19<sup>a</sup> | 20.04±1.06<sup>a</sup> |
| Control (Traditional fermentation method) | 30.02±1.08<sup>a</sup> | 32.27±1.14<sup>ab</sup> | 32.64±1.42<sup>b</sup> |
| Treatment 1 (1.5/0.16) | 42.01±1.66<sup>b</sup> | 34.45±1.23<sup>b</sup> | 43.08±1.63<sup>c</sup> |
| Treatment 2 (1.3/0.12) | 43.29±2.10<sup>b</sup> | 35.59±1.47<sup>b</sup> | 47.89±2.10<sup>d</sup> |
| Treatment 3 (1.7/0.19) | 49.64±1.70<sup>c</sup> | 39.18±1.07<sup>c</sup> | 53.97±1.70<sup>e</sup> |

Note: Values Values are mean±SD (n=3). Means bearing different superscript letters (a,b,c,d,e) within the same column differ significantly from one another (one-way ANOVA followed by Tukey's HSD post hoc test,p<0.05)

## 4.0 Conclusion

This study demonstrated that electro-fermentation (EF) using *Candida tropicalis* SY005 mediated by carbon fiber cloth electrode and Fe_3_O_4_ redox active particles, significantly increase the fermentation kinetics while enhancing ethanol production as well bioactive compound in grape must fermentation. Use of low electric field (1.3-1.7 V, 0.12-0.19 mA) formed strongly reduced inracellular environmet (ORP: -100 to -143mV), which significantly reduced total fermentation time from 264h in the control to just 72 h across all treatments a almost 3/7 times acceleration. Such reducing redox redox condition changed the intracellular NADH/NAD^+^ ratio stimulating glycolytic flux while enhancing NAD^+^ regeneration and consequently increased ethanol production from 7.00±0.26% to 13.00±0.40% (Treatment 3) F(3,8)=211.97, P<0.001.

FTIR evaluation confirmed this metabolic shift revealing a considerable decline in transmittance at the O-H (33357-3260 cm^-1^), C-H (2999-2992 cm^-1^), and C=O/C-O (1641 and 1045 cm-1) infrared absorption spectral bands within electro-fermentation sample relative to control group in agreement with higher ethanol and polyol production. GC-MS analysis further confirmed the observed redox mediated alteration identifying 2,3butanediol (13.05%) and glycerin (14.33%) as predominant highly reduced metabolites both established markers of increased NADH dependent flux during perturbed cellular oxidation-reduction state. FE-SEM together with EDS analysis revealed that Fe_3_O_4_ particle created a direct physical association with the yeast cell wall, forming a biological interaction zone with an iron content of 17.86 wt %. This interfacial region probably act as a redox conduit for intracellular electron transfer consistent with the suggested process of electrode-to-cell electron movement without the toxicity. Apart from fermentation kinetics electro-fermentation significantly improved the release of bio-active compounds, with total phenolic (49.64±2.10 mgGAE/100mL), flavonoid (39.18±1.07 mgRUE/100mL) and anthocyanin (53.87±2.10 mg/100mL) all reaching their highest value in treatment 3 (one-way ANOVA TPC: F(4,10)=97.66, TFC: F(4,10)=25.42, F(4,10)=205.12, all p<0.001). this attributes to electric-field-induced electroporation of grape skin and pulp cell membrane structure altering cellular compartmentation and improving the release of bound phenolic, flavonoid and anthocyanin compounds, that would otherwise remain in bound form to cell wall polysaccharides matrix during traditional fermentation process. Collectively these outcomes establish electro-fermentation as a sustainable approach for accelerating fermentation kinetics while improving the bioactive profile of fermented beverages. Future studies should focus on validating the proposed mechanism under pilot-scale condition evaluating prolonged electrode stability and Fe_3_O_4_ particle recovery, reuse as well as characterizing organoleptic and volatile flavor composition changes linked with electro-fermentation.

