## supplemental for "Electro-Fermentation of Grape Must via *Candida tropicalis* SY005: Accelerating Kinetics, Modulating Bio-chemical Pathway and Improving Bio-active Content": GC-MS.pdf

### Library Search Report

| | | | |
|---|---|---|---|
| Data File: | 28052024_T3 | Original Data Path: | C:\FTL\Data |
| Current Data Path: | C:\FTL\Data | Sample Type: | Unknown |
| Sample ID: | 5 | Sample Name: | |
| Imran Ansari | TSQ81802503 | Acquisition Date: | 05/28/24 05:21:36 PM |
| Run Time(min): | 45.69 | Comments: | |
| Vial: | 8 | Scans: | 13548 |
| Low Mass(m/z): | 45.00000 | High Mass(m/z): | 450.00702 |
| Sample Weight: | 0.00 | ISTD Amount: | 0.000 |
| Calibration Level: | | Dilution Factor: | 1.00 |
| Instrument Method: | C:\FTL\Instrument Method\PrinceMusExtract.meth | | |
| Original Processing Method: | C:\FTL\Processing Method\XYZ_EXTRACT | | |
| Current Processing Method: | N/A | | |

RT: 0.00 - 49.69 SM: 7B

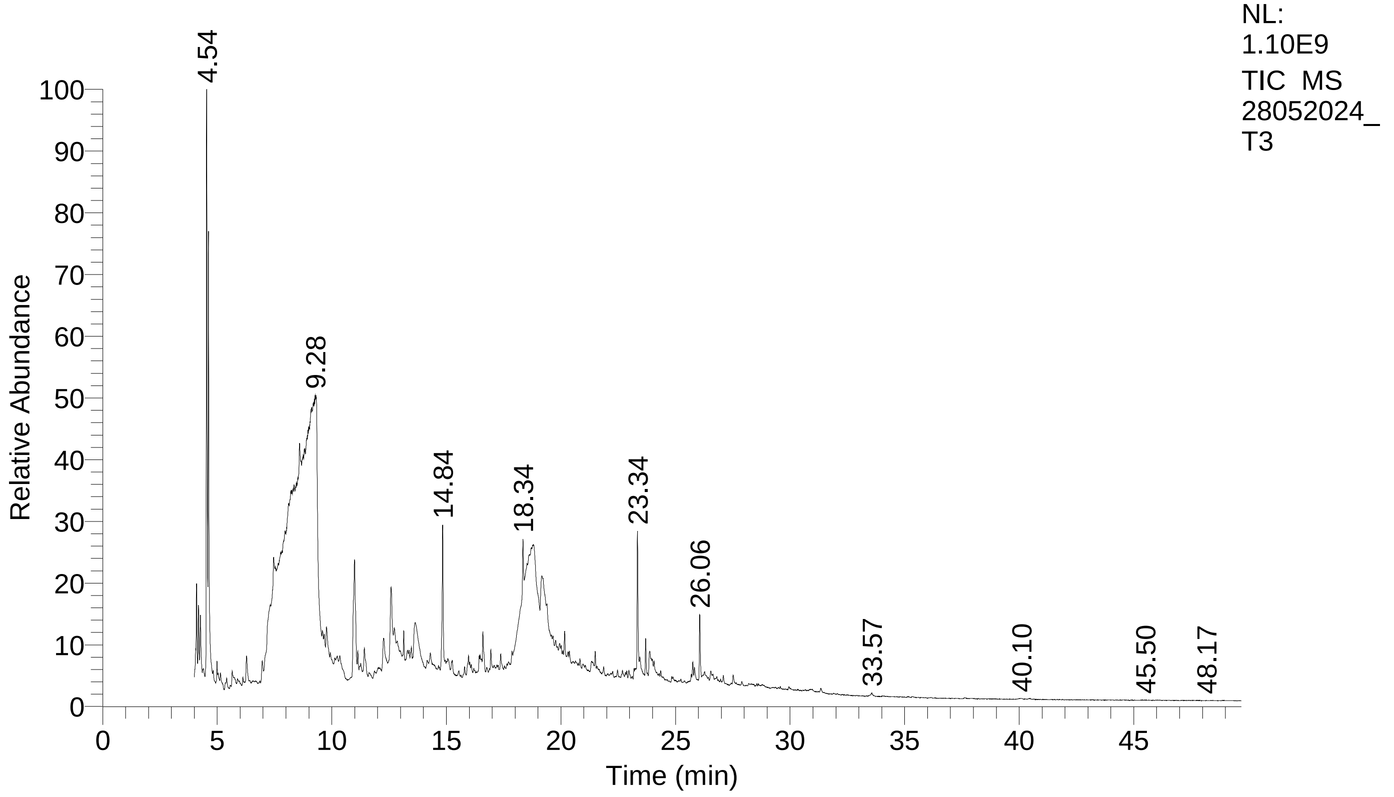

### Library Search Report

#### Lib. Search Graphics Table

| Compound Structure | Hit Spectrum |
|---|---|

Acetoin
Formula C4H8O2, MW 88, CAS# 513-86-0, Entry# 16903
2-Butanone, 3-hydroxy-

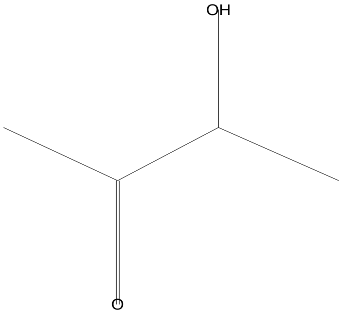


SI 808, RSI 869, mainlib, Entry# 16903, CAS# 513-86-0, Acetoin

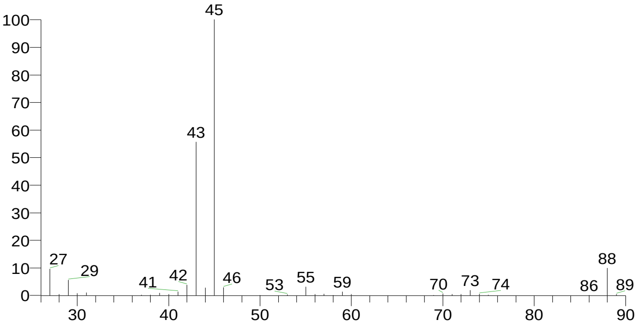


Hydrazine, (2-methylpropyl)-
Formula C4H12N2, MW 88, CAS# 42504-87-0, Entry# 17360
Isobutylhydrazine

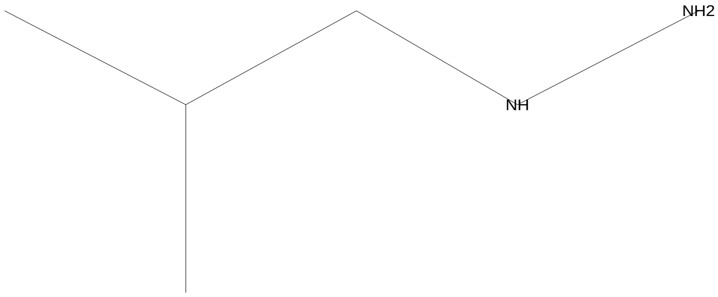


SI 765, RSI 845, mainlib, Entry# 17360, CAS# 42504-87-0, Hydrazine, (2-methylpropyl)-

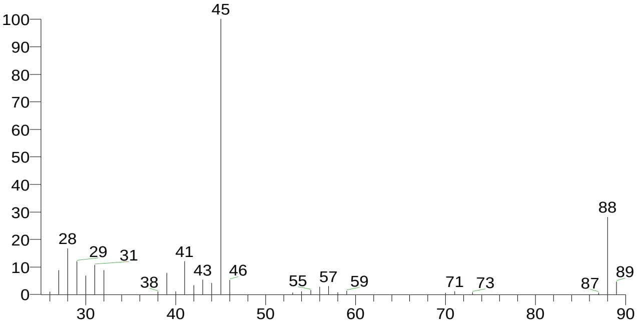


(3-Methyl-oxiran-2-yl)-methanol
Formula C4H8O2, MW 88, CAS# NA, Entry# 6815
(3-Methyl-2-oxiranyl)methanol #

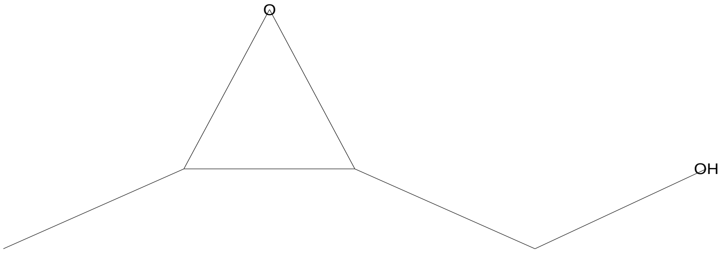


SI 749, RSI 782, mainlib, Entry# 6815, CAS# NA, (3-Methyl-oxiran-2-yl)-methanol

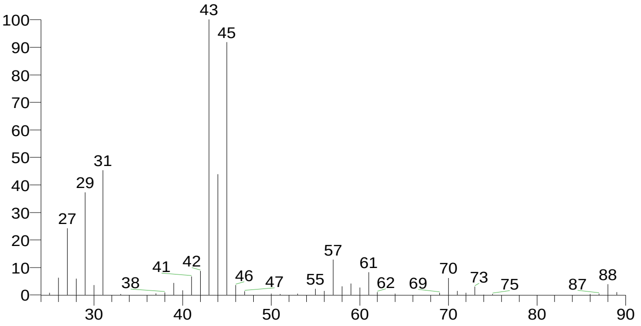


Acetic acid, methoxy-, ethyl ester
Formula C5H10O3, MW 118, CAS# 3938-96-3, Entry# 3941
Ethyl methoxyacetate

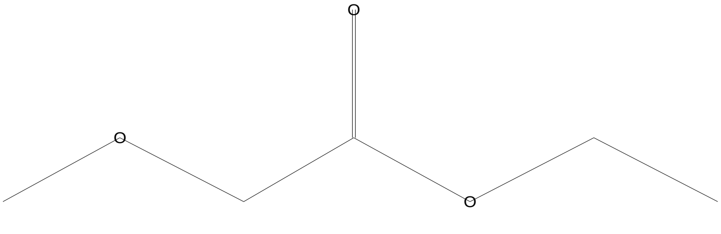


SI 743, RSI 881, replib, Entry# 3941, CAS# 3938-96-3, Acetic acid, methoxy-, ethyl ester

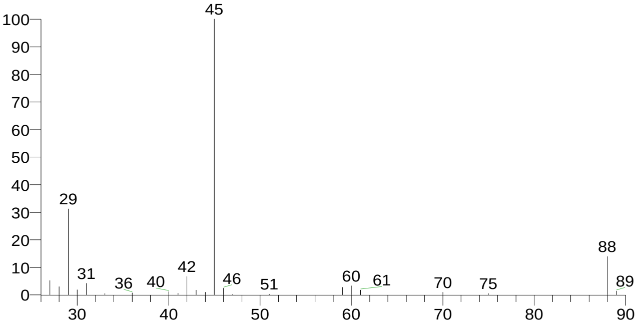

### Library Search Report

Compound Structure

Propane, 1-methoxy-2-methyl-
Formula C5H12O, MW 88, CAS# 625-44-5, Entry# 16742
Ether, isobutyl methyl

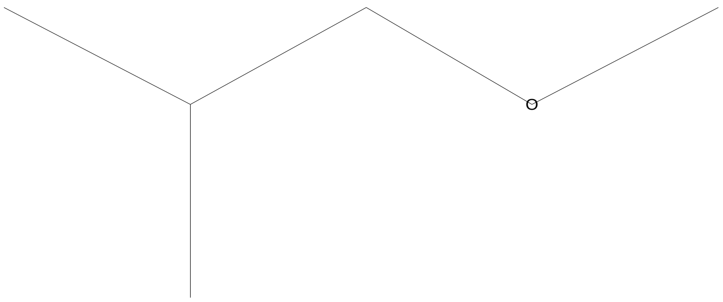


Hit Spectrum

SI 729, RSI 770, mainlib, Entry# 16742, CAS# 625-44-5, Propane, 1-methoxy-2-methyl-

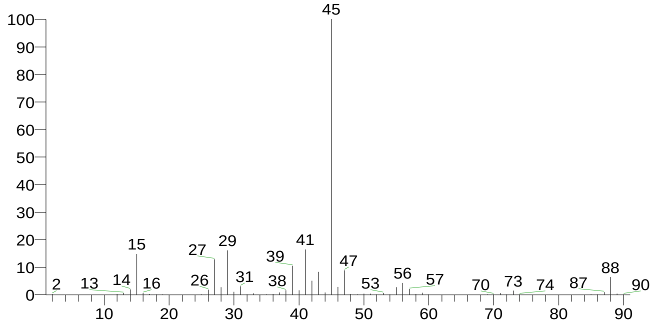


| RT | Compound Name | Area % | Molecular Weight | Molecular Formula | Probability |
|---|---|---|---|---|---|
| 4.09 | Acetoin | 2.69 | 88 | C4H8O2 | 62.51 |
| 4.09 | Hydrazine, (2-methylpropyl)- | 2.69 | 88 | C4H12N2 | 13.78 |
| 4.09 | (3-Methyl-oxiran-2-yl)-methanol | 2.69 | 88 | C4H8O2 | 7.94 |
| 4.09 | Acetic acid, methoxy-, ethyl ester | 2.69 | 118 | C5H10O3 | 6.24 |
| 4.09 | Propane, 1-methoxy-2-methyl- | 2.69 | 88 | C5H12O | 3.90 |

There is no signature data to report. Anamika Singh

### Library Search Report

#### Lib. Search Graphics Table

| Compound Structure | Hit Spectrum |
|---|---|

2,3-Butanediol
Formula C4H10O2, MW 90, CAS# 513-85-9, Entry# 3932
Butane-2,3-diol

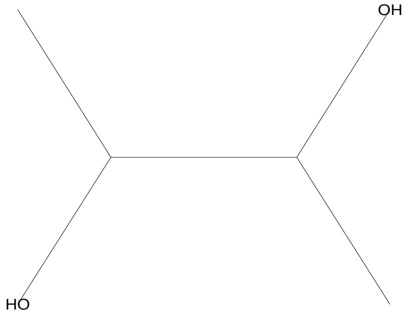


SI 755, RSI 936, replib, Entry# 3932, CAS# 513-85-9, 2,3-Butanediol

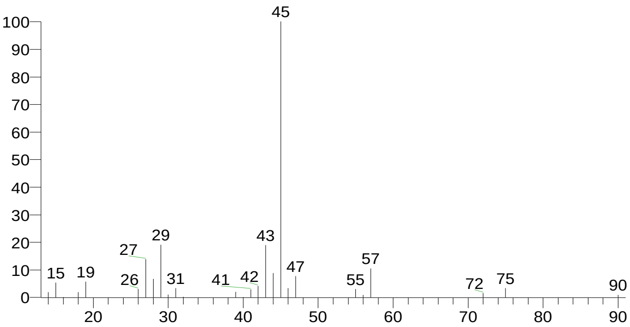


2,3-Butanediol
Formula C4H10O2, MW 90, CAS# 513-85-9, Entry# 3977
Butane-2,3-diol

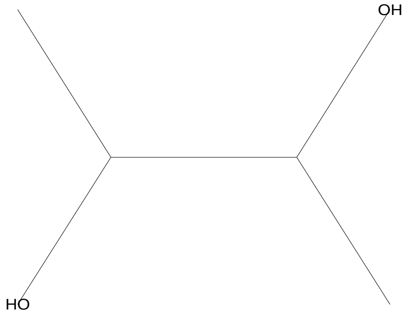


SI 743, RSI 834, replib, Entry# 3977, CAS# 513-85-9, 2,3-Butanediol

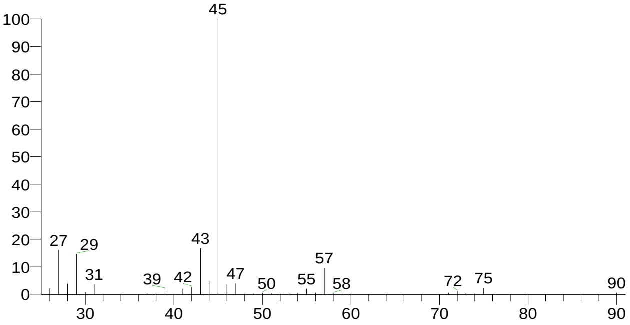


2,3-Butanediol, [R-(R*,R*)]-
Formula C4H10O2, MW 90, CAS# 24347-58-8, Entry# 16854
2,3-Butanediol #

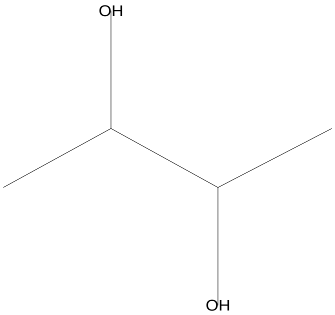


SI 740, RSI 830, mainlib, Entry# 16854, CAS# 24347-58-8, 2,3-Butanediol, [R-(R*,R*)]-

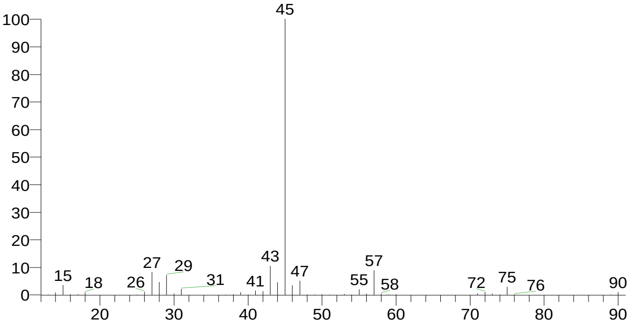


Acetic acid, ethoxyhydroxy-, ethyl ester
Formula C6H12O4, MW 148, CAS# 49653-17-0, Entry# 438
Ethyl ethoxy(hydroxy)acetate #

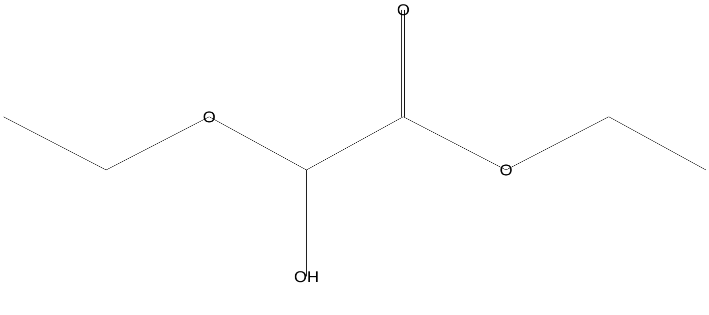


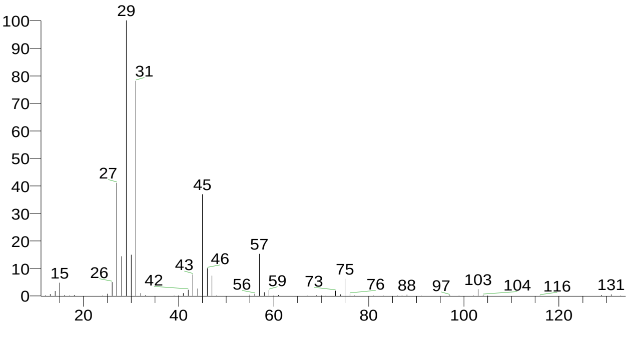


There is no signature data to report. Anamika Singh

### Library Search Report

| Compound Structure | Hit Spectrum |
|---|---|
| 2,3-Butanediol<br>Formula C4H10O2, MW 90, CAS# 513-85-9, Entry# 16849<br>Butane-2,3-diol | SI 733, RSI 827, mainlib, Entry# 16849, CAS# 513-85-9, 2,3-Butanediol |

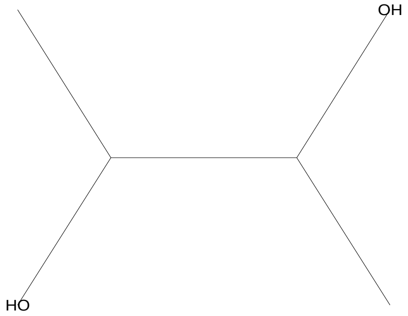


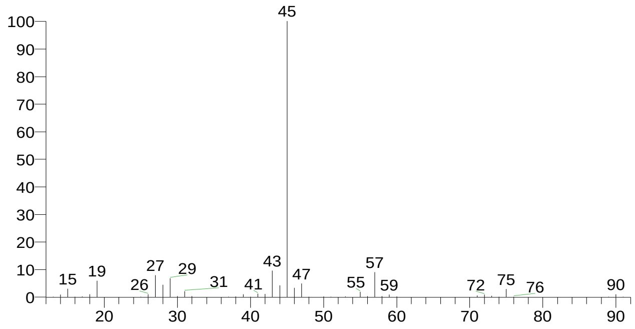


| RT | Compound Name | Area % | Molecular Weight | Molecular Formula | Probability |
|---|---|---|---|---|---|
| 4.18 | 2,3-Butanediol | 1.74 | 90 | C4H10O2 | 28.37 |
| 4.18 | 2,3-Butanediol | 1.74 | 90 | C4H10O2 | 28.37 |
| 4.18 | 2,3-Butanediol | 1.74 | 90 | C4H10O2 | 28.37 |
| 4.18 | 2,3-Butanediol, [R-(R*,R*)]- | 1.74 | 90 | C4H10O2 | 17.19 |
| 4.18 | Acetic acid, ethoxyhydroxy-, ethyl ester | 1.74 | 148 | C6H12O4 | 14.52 |

### Library Search Report

#### Lib. Search Graphics Table

| Compound Structure | Hit Spectrum |
|---|---|

1,2,3-Butanetriol
Formula C4H10O3, MW 106, CAS# 4435-50-1, Entry# 15460
$:28YAXKTBLXMTYWDQ-UHFFFAOYSA-N

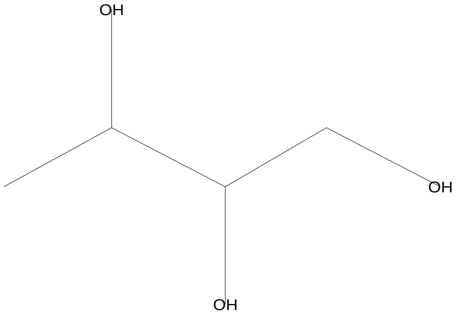


SI 700, RSI 761, mainlib, Entry# 15460, CAS# 4435-50-1, 1,2,3-Butanetriol

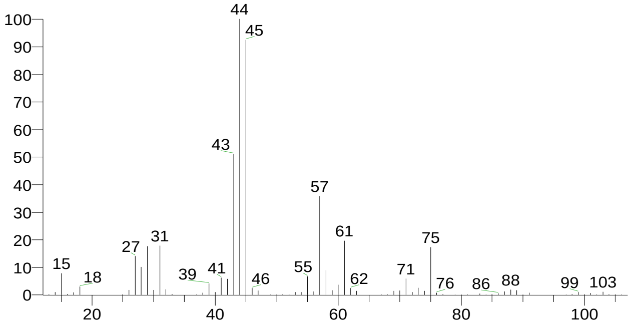


(3-Methyl-oxiran-2-yl)-methanol
Formula C4H8O2, MW 88, CAS# NA, Entry# 6815
(3-Methyl-2-oxiranyl)methanol #

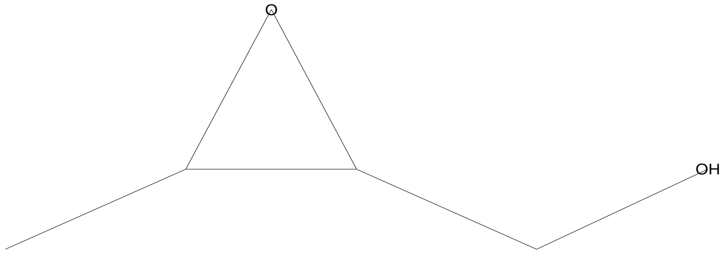


SI 697, RSI 805, mainlib, Entry# 6815, CAS# NA, (3-Methyl-oxiran-2-yl)-methanol

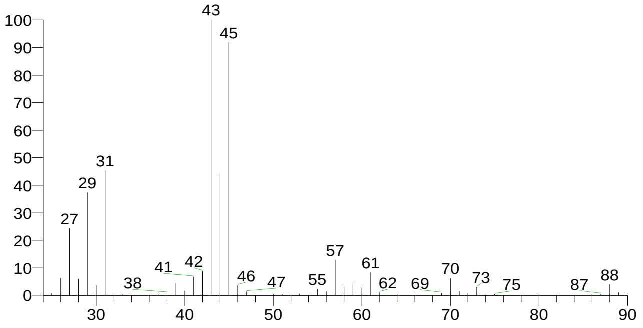


2,3-Butanediol
Formula C4H10O2, MW 90, CAS# 513-85-9, Entry# 3932
Butane-2,3-diol

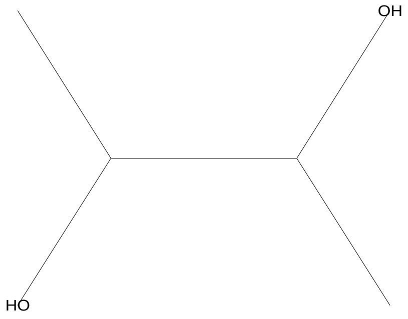


SI 690, RSI 948, replib, Entry# 3932, CAS# 513-85-9, 2,3-Butanediol

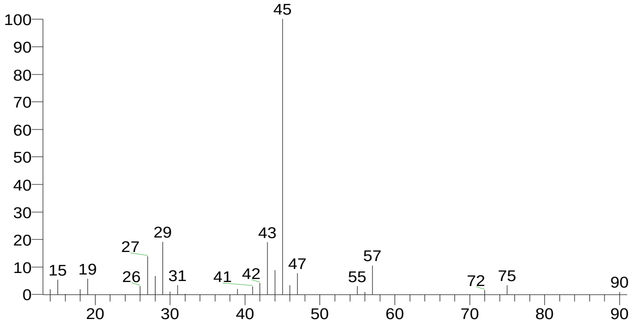


2,3-Butanediol
Formula C4H10O2, MW 90, CAS# 513-85-9, Entry# 3977
Butane-2,3-diol

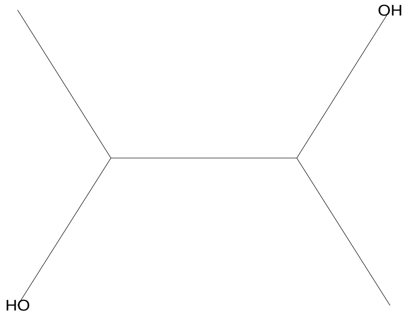


SI 690, RSI 838, replib, Entry# 3977, CAS# 513-85-9, 2,3-Butanediol

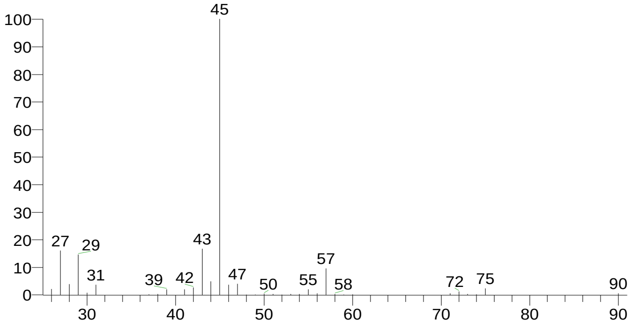

### Library Search Report

| Compound Structure | Hit Spectrum |
|---|---|
| Lactic acid<br>Formula C3H6O3, MW 90, CAS# 50-21-5, Entry# 3913<br>Propanoic acid, 2-hydroxy- | SI 665, RSI 731, replib, Entry# 3913, CAS# 50-21-5, Lactic acid |

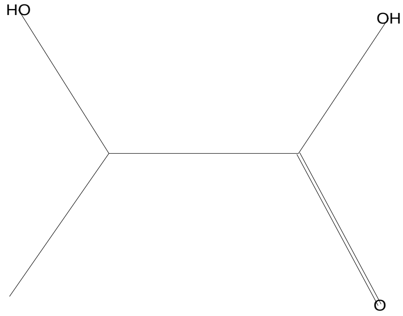


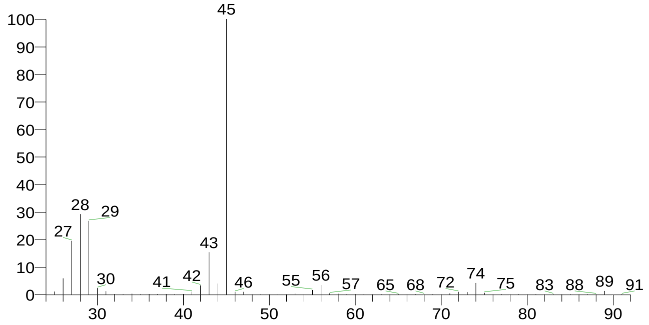


| RT | Compound Name | Area % | Molecular Weight | Molecular Formula | Probability |
|---|---|---|---|---|---|
| 4.26 | 1,2,3-Butanetriol | 1.84 | 106 | C4H10O3 | 21.27 |
| 4.26 | (3-Methyl-oxiran-2-yl)-methanol | 1.84 | 88 | C4H8O2 | 18.79 |
| 4.26 | 2,3-Butanediol | 1.84 | 90 | C4H10O2 | 14.39 |
| 4.26 | 2,3-Butanediol | 1.84 | 90 | C4H10O2 | 14.39 |
| 4.26 | Lactic acid | 1.84 | 90 | C3H6O3 | 4.39 |

There is no signature data to report. Anamika Singh

### Library Search Report

#### Lib. Search Graphics Table

| Compound Structure | Hit Spectrum |
|---|---|

2,3-Butanediol, [R-(R*,R*)]-
Formula C4H10O2, MW 90, CAS# 24347-58-8, Entry# 16854
2,3-Butanediol #

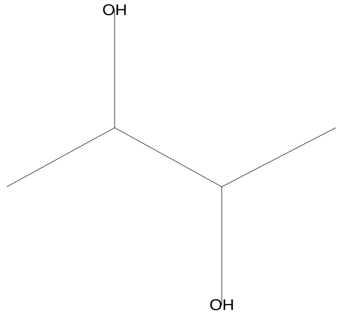


SI 863, RSI 865, mainlib, Entry# 16854, CAS# 24347-58-8, 2,3-Butanediol, [R-(R*,R*)]-

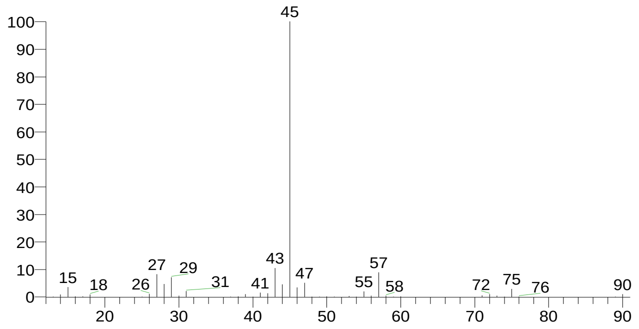


2,3-Butanediol
Formula C4H10O2, MW 90, CAS# 513-85-9, Entry# 16849
Butane-2,3-diol

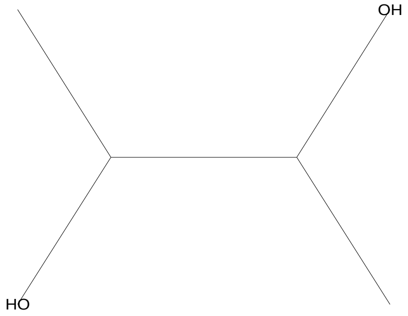


SI 841, RSI 843, mainlib, Entry# 16849, CAS# 513-85-9, 2,3-Butanediol

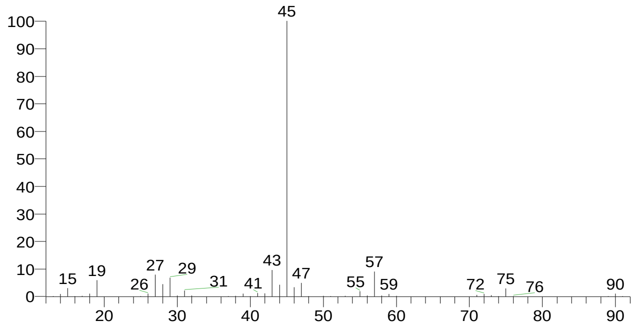


2,3-Butanediol, [S-(R*,R*)]-
Formula C4H10O2, MW 90, CAS# 19132-06-0, Entry# 16803
(2S,3S)-(+)-2,3-Butanediol

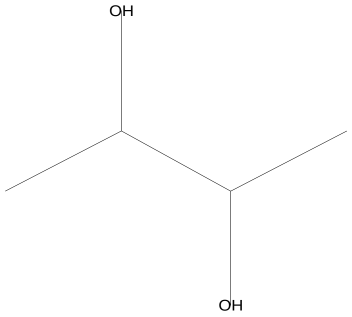


SI 832, RSI 856, mainlib, Entry# 16803, CAS# 19132-06-0, 2,3-Butanediol, [S-(R*,R*)]-

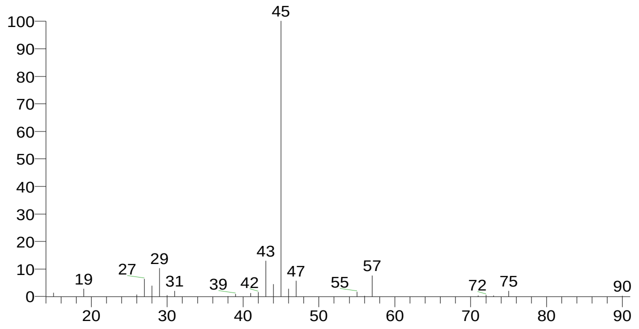


2,3-Butanediol
Formula C4H10O2, MW 90, CAS# 513-85-9, Entry# 3977
Butane-2,3-diol

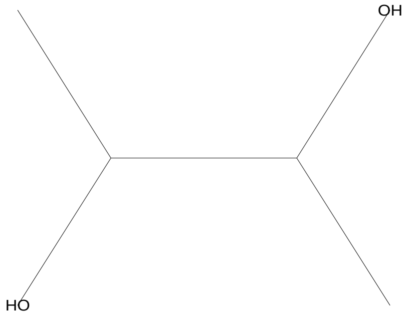


SI 800, RSI 806, replib, Entry# 3977, CAS# 513-85-9, 2,3-Butanediol

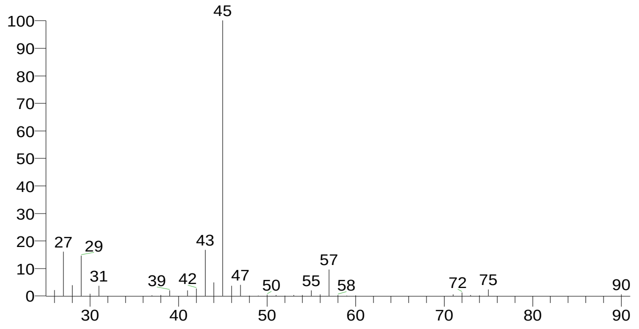

### Library Search Report

Compound Structure

2,3-Butanediol
Formula C4H10O2, MW 90, CAS# 513-85-9, Entry# 3932
Butane-2,3-diol

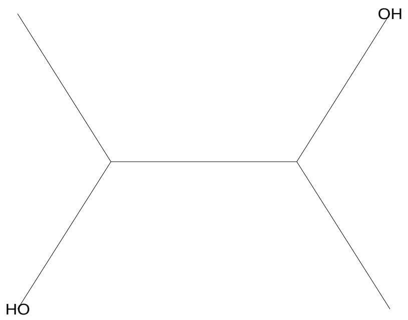


Hit Spectrum

SI 789, RSI 842, replib, Entry# 3932, CAS# 513-85-9, 2,3-Butanediol

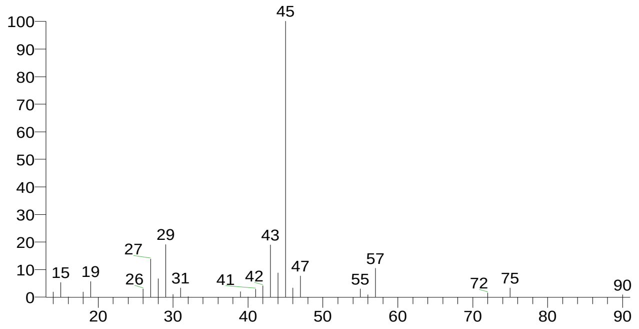


| RT | Compound Name | Area % | Molecular Weight | Molecular Formula | Probability |
|---|---|---|---|---|---|
| 4.54 | 2,3-Butanediol, [R-(R*,R*)]- | 13.05 | 90 | C4H10O2 | 56.19 |
| 4.54 | 2,3-Butanediol | 13.05 | 90 | C4H10O2 | 22.21 |
| 4.54 | 2,3-Butanediol | 13.05 | 90 | C4H10O2 | 22.21 |
| 4.54 | 2,3-Butanediol | 13.05 | 90 | C4H10O2 | 22.21 |
| 4.54 | 2,3-Butanediol, [S-(R*,R*)]- | 13.05 | 90 | C4H10O2 | 16.12 |

There is no signature data to report. Anamika Singh

### Library Search Report

#### Lib. Search Graphics Table

| Compound Structure | Hit Spectrum |
|---|---|

2,3-Butanediol, [R-(R*,R*)]-
Formula C4H10O2, MW 90, CAS# 24347-58-8, Entry# 16854
2,3-Butanediol #

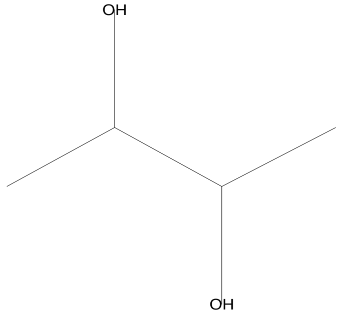


SI 884, RSI 896, mainlib, Entry# 16854, CAS# 24347-58-8, 2,3-Butanediol, [R-(R*,R*)]-

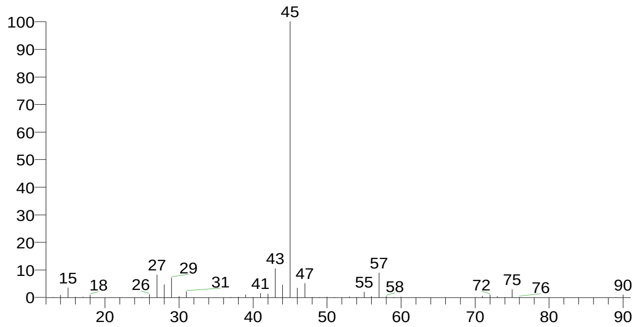


2,3-Butanediol, [S-(R*,R*)]-
Formula C4H10O2, MW 90, CAS# 19132-06-0, Entry# 16803
(2S,3S)-(+)-2,3-Butanediol

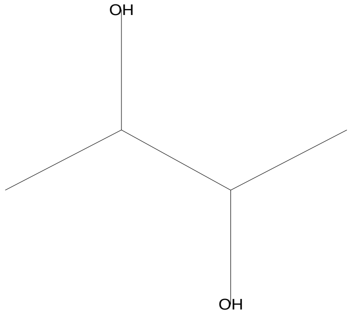


SI 873, RSI 908, mainlib, Entry# 16803, CAS# 19132-06-0, 2,3-Butanediol, [S-(R*,R*)]-

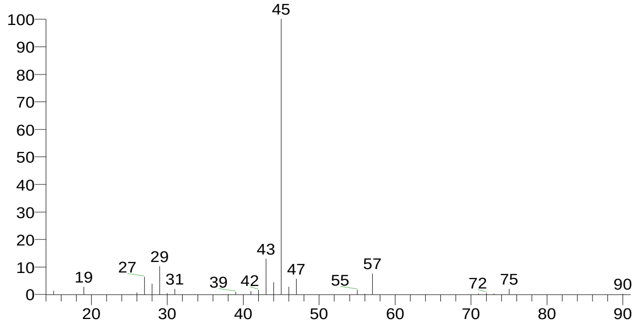


2,3-Butanediol
Formula C4H10O2, MW 90, CAS# 513-85-9, Entry# 16849
Butane-2,3-diol

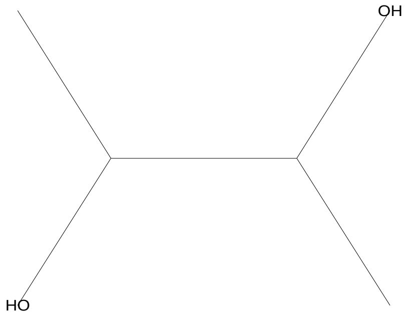


SI 862, RSI 874, mainlib, Entry# 16849, CAS# 513-85-9, 2,3-Butanediol

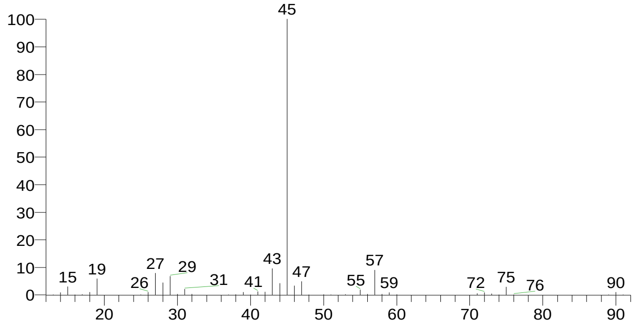


2,3-Butanediol
Formula C4H10O2, MW 90, CAS# 513-85-9, Entry# 3932
Butane-2,3-diol

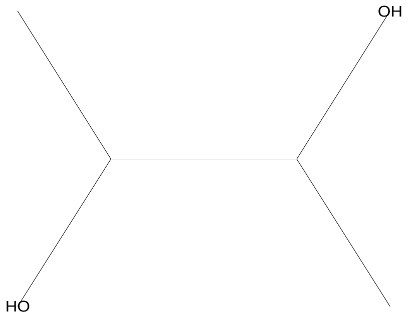


SI 826, RSI 887, replib, Entry# 3932, CAS# 513-85-9, 2,3-Butanediol

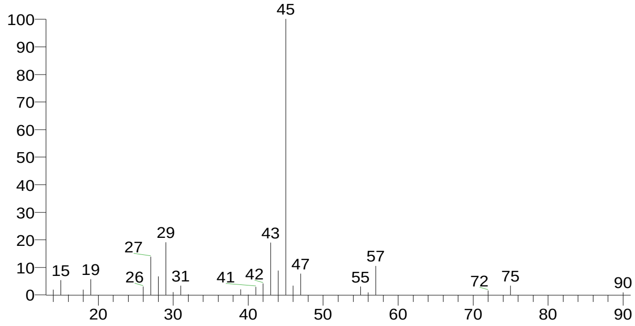

### Library Search Report

Compound Structure

2,3-Butanediol
Formula C4H10O2, MW 90, CAS# 513-85-9, Entry# 3977
Butane-2,3-diol

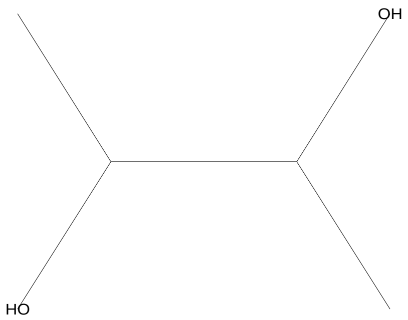


Hit Spectrum

SI 818, RSI 833, replib, Entry# 3977, CAS# 513-85-9, 2,3-Butanediol

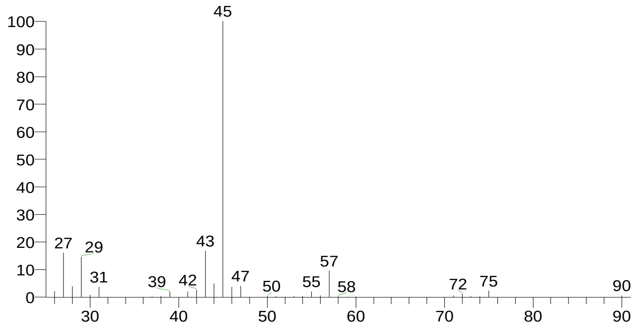


| RT | Compound Name | Area % | Molecular Weight | Molecular Formula | Probability |
|---|---|---|---|---|---|
| 4.61 | 2,3-Butanediol, [R-(R*,R*)]- | 9.54 | 90 | C4H10O2 | 44.00 |
| 4.61 | 2,3-Butanediol, [S-(R*,R*)]- | 9.54 | 90 | C4H10O2 | 30.19 |
| 4.61 | 2,3-Butanediol | 9.54 | 90 | C4H10O2 | 20.71 |
| 4.61 | 2,3-Butanediol | 9.54 | 90 | C4H10O2 | 20.71 |
| 4.61 | 2,3-Butanediol | 9.54 | 90 | C4H10O2 | 20.71 |

There is no signature data to report. Anamika Singh

### Library Search Report

#### Lib. Search Graphics Table

| Compound Structure | Hit Spectrum |
|---|---|

6-Oxa-bicyclo[3.1.0]hexan-3-one
Formula C5H6O2, MW 98, CAS# 74017-10-0, Entry# 4491
$:28VZMRGSWFCBXBPW-UHFFFAOYSA-N

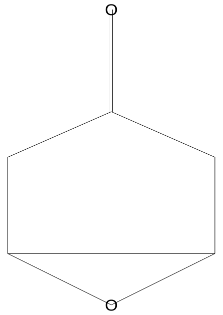


SI 714, RSI 861, mainlib, Entry# 4491, CAS# 74017-10-0, 6-Oxa-bicyclo[3.1.0]hexan-3-one

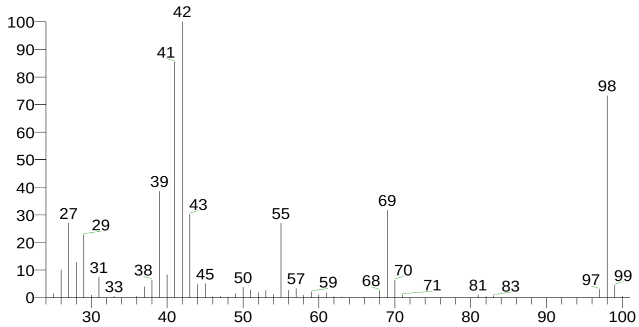


1-(3,3,3-Trifluoro-2-hydroxypropyl)piperidine
Formula C8H14F3NO, MW 197, CAS# NA, Entry# 71001
1,1,1-Trifluoro-3-(1-piperidinyl)-2-propanol #

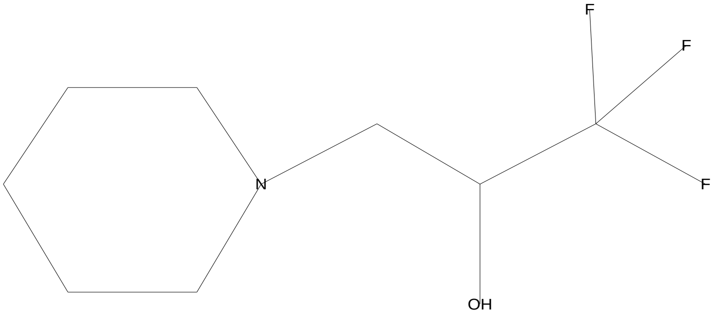


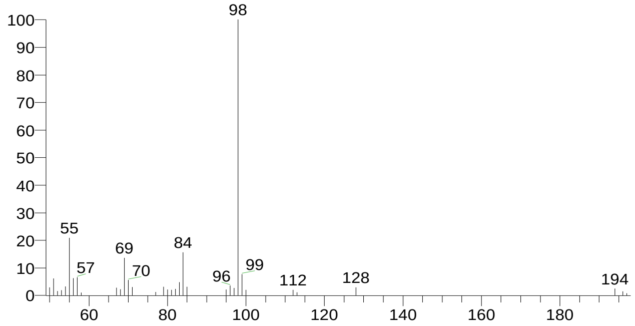


1-Pyridineacetic acid, hexahydro-
Formula C7H13NO2, MW 143, CAS# NA, Entry# 70724
$:28VRDBIJCCXDEZJN-UHFFFAOYSA-N

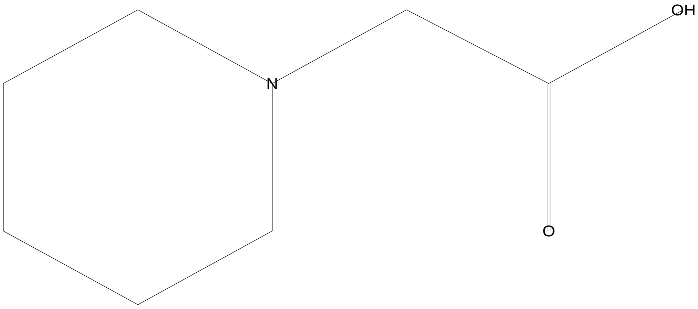


SI 656, RSI 803, mainlib, Entry# 70724, CAS# NA, 1-Pyridineacetic acid, hexahydro-

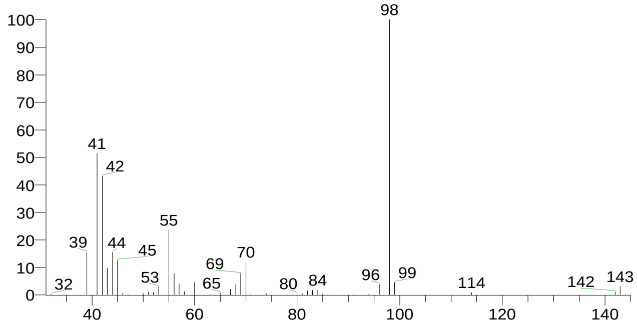


Carbonic acid, (ethyl)(1,2,4-triazol-1-ylmethyl) diester
Formula C6H9N3O3, MW 171, CAS# NA, Entry# 51601
Ethyl 1H-1,2,4-triazol-1-ylmethyl carbonate #

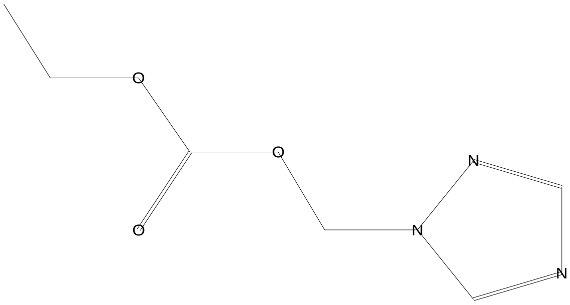


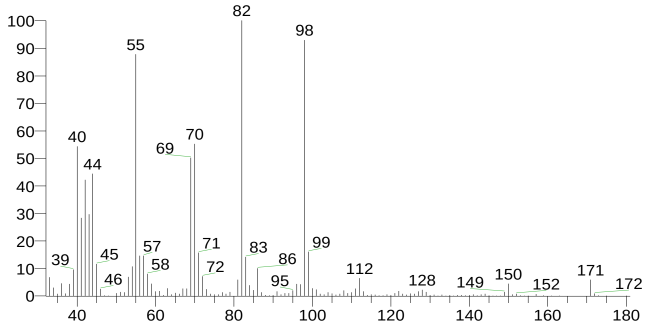


There is no signature data to report. Anamika Singh

### Library Search Report

Compound Structure

6-Oxabicyclo[3.1.0]hexan-2-one
Formula C5H6O2, MW 98, CAS# 6705-52-8, Entry# 4437
Cyclopentanone, 2,3-epoxy-

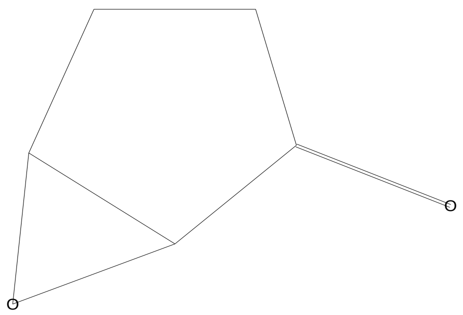


Hit Spectrum

SI 654, RSI 859, mainlib, Entry# 4437, CAS# 6705-52-8, 6-Oxabicyclo[3.1.0]hexan-2-one

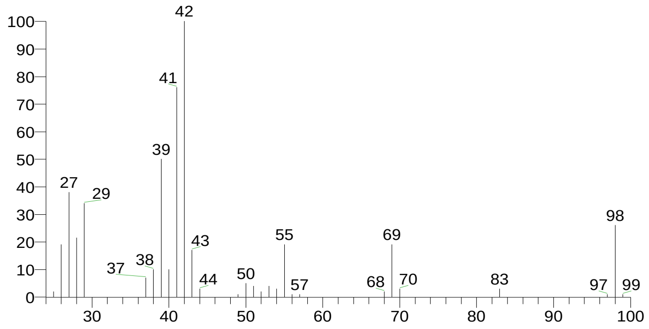


| RT | Compound Name | Area % | Molecular Weight | Molecular Formula | Probability |
|---|---|---|---|---|---|
| 6.28 | 6-Oxa-bicyclo[3.1.0]hexan-3-one | 1.02 | 98 | C5H6O2 | 27.82 |
| 6.28 | 1-(3,3,3-Trifluoro-2-hydroxypropyl)piperidine | 1.02 | 197 | C8H14F3NO | 5.76 |
| 6.28 | Carbonic acid, (ethyl)(1,2,4-triazol-1-ylmethyl) diester | 1.02 | 171 | C6H9N3O3 | 4.07 |
| 6.28 | 1-Pyridineacetic acid, hexahydro- | 1.02 | 143 | C7H13NO2 | 4.07 |
| 6.28 | 6-Oxabicyclo[3.1.0]hexan-2-one | 1.02 | 98 | C5H6O2 | 3.75 |

### Library Search Report

#### Lib. Search Graphics Table

| Compound Structure | Hit Spectrum |
|---|---|

Desulphosinigrin
Formula C10H17NO6S, MW 279, CAS# 5115-81-1, Entry# 30961
1-S-[(1E)-N-Hydroxy-3-butenimidoyl]-1-thiohexopyranose #

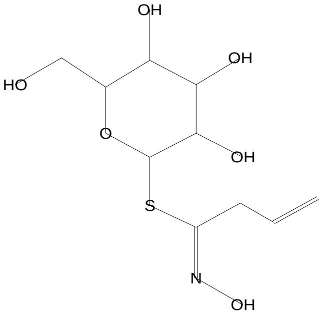


SI 673, RSI 686, mainlib, Entry# 30961, CAS# 5115-81-1, Desulphosinigrin

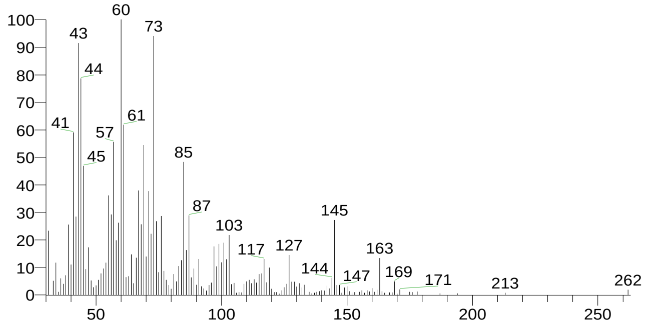


d-Mannose
Formula C6H12O6, MW 180, CAS# 3458-28-4, Entry# 41111
Mannose, d-

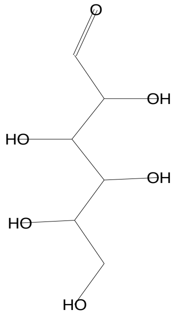


SI 648, RSI 659, mainlib, Entry# 41111, CAS# 3458-28-4, d-Mannose

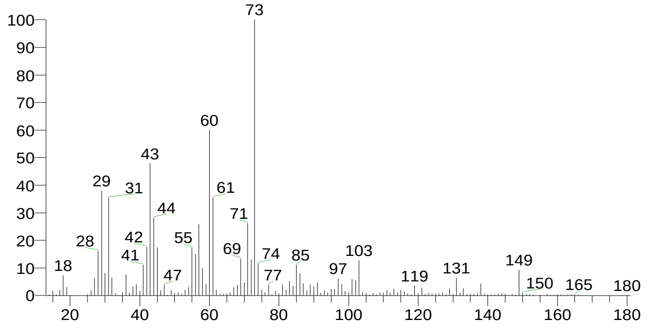


Melezitose
Formula C18H32O16, MW 504, CAS# 597-12-6, Entry# 41139
à-D-Glucopyranoside, O-à-D-glucopyranosyl-(1.fwdarw.3)-á-D-fructofuranosyl

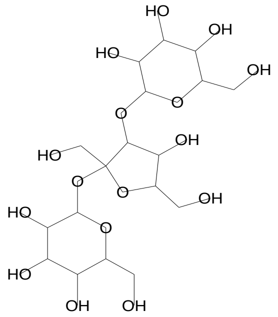


SI 648, RSI 655, mainlib, Entry# 41139, CAS# 597-12-6, Melezitose

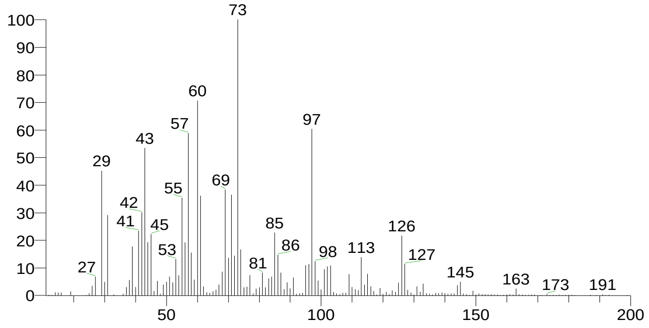


d-Glycero-d-ido-heptose
Formula C7H14O7, MW 210, CAS# NA, Entry# 9206
Heptose #

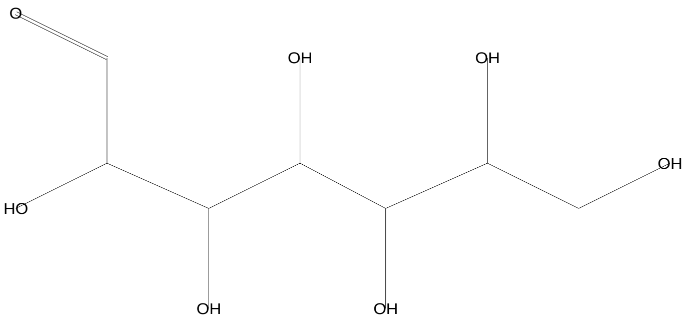


SI 646, RSI 658, mainlib, Entry# 9206, CAS# NA, d-Glycero-d-ido-heptose

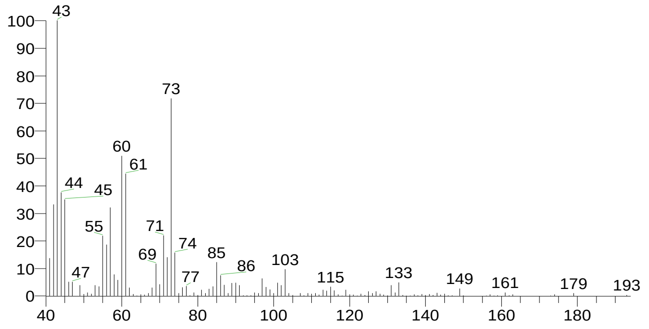

### Library Search Report

| Compound Structure | Hit Spectrum |
|---|---|

Diglycerol
Formula C6H14O5, MW 166, CAS# 627-82-7, Entry# 31152
1,2-Propanediol, 3,3'-oxybis-

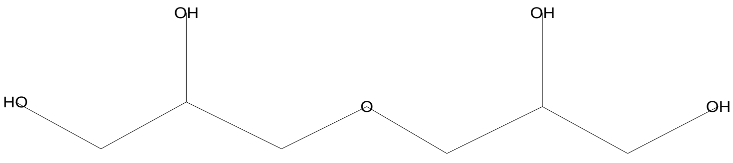


SI 642, RSI 708, mainlib, Entry# 31152, CAS# 627-82-7, Diglycerol

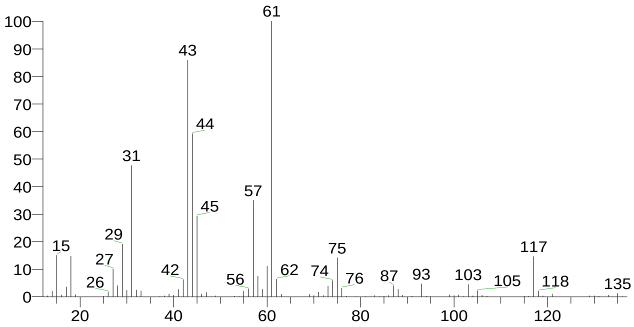


| RT | Compound Name | Area % | Molecular Weight | Molecular Formula | Probability |
|---|---|---|---|---|---|
| 7.20 | Desulphosinigrin | 1.19 | 279 | C10H17NO6S | 18.90 |
| 7.20 | Melezitose | 1.19 | 504 | C18H32O16 | 5.76 |
| 7.20 | d-Mannose | 1.19 | 180 | C6H12O6 | 5.76 |
| 7.20 | d-Glycero-d-ido-heptose | 1.19 | 210 | C7H14O7 | 5.31 |
| 7.20 | Diglycerol | 1.19 | 166 | C6H14O5 | 4.49 |

### Library Search Report

#### Lib. Search Graphics Table

| Compound Structure | Hit Spectrum |
|---|---|

Diglycerol
Formula C6H14O5, MW 166, CAS# 627-82-7, Entry# 31152
1,2-Propanediol, 3,3'-oxybis-

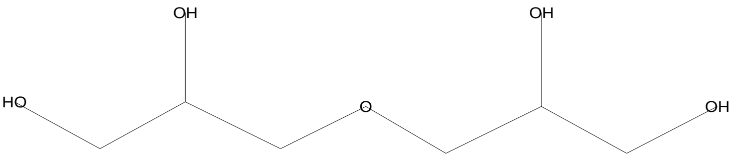


SI 732, RSI 753, mainlib, Entry# 31152, CAS# 627-82-7, Diglycerol

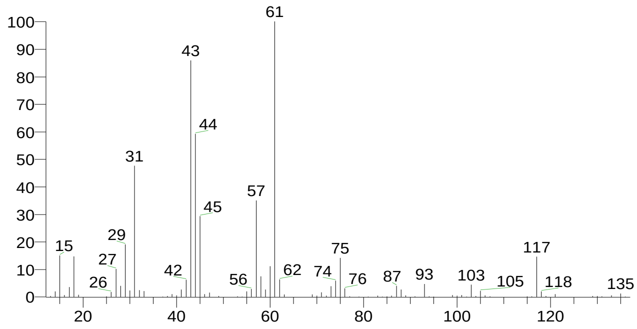


Glycerin
Formula C3H8O3, MW 92, CAS# 56-81-5, Entry# 7604
1,2,3-Propanetriol

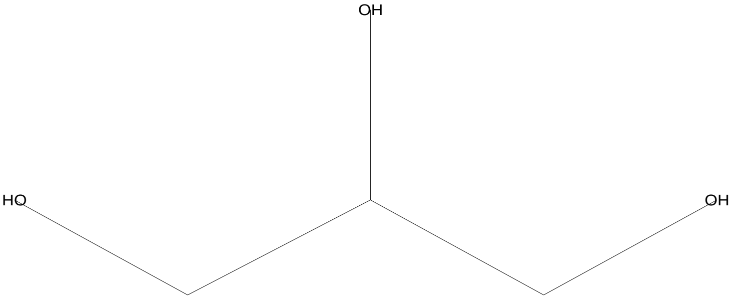


SI 696, RSI 774, replib, Entry# 7604, CAS# 56-81-5, Glycerin

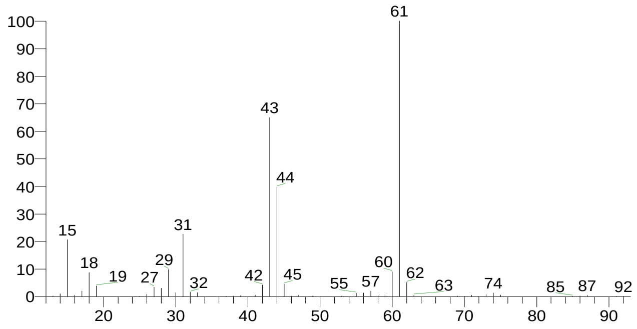


Glycerin
Formula C3H8O3, MW 92, CAS# 56-81-5, Entry# 7605
1,2,3-Propanetriol

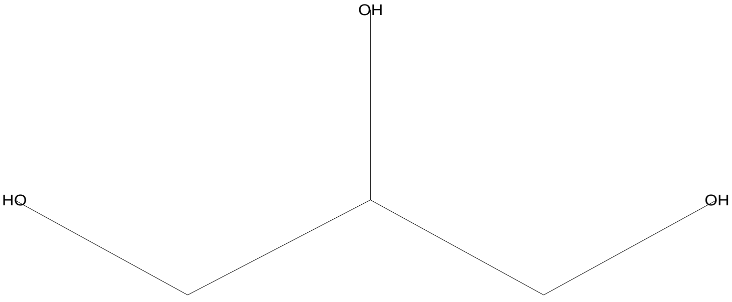


SI 682, RSI 781, replib, Entry# 7605, CAS# 56-81-5, Glycerin

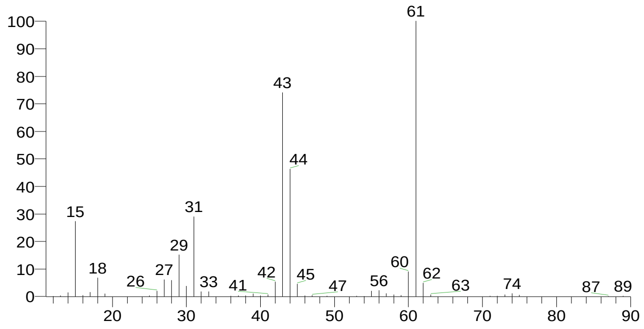


Erythritol
Formula C4H10O4, MW 122, CAS# 149-32-6, Entry# 31179
2(R),3(S)-1,2,3,4-Butanetetrol

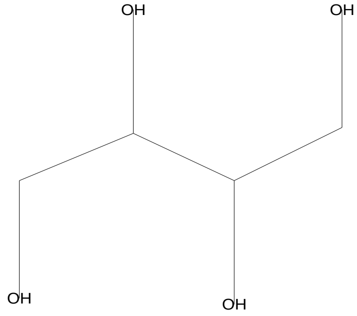


SI 674, RSI 768, mainlib, Entry# 31179, CAS# 149-32-6, Erythritol

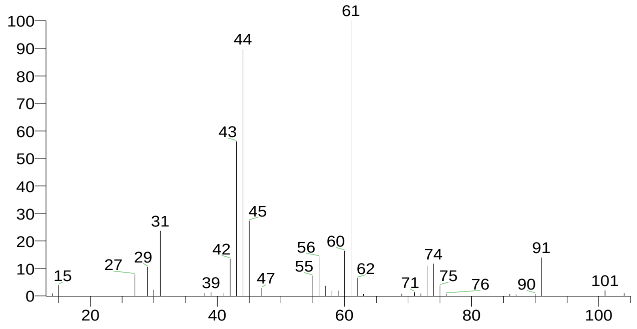

### Library Search Report

| Compound Structure | Hit Spectrum |
|---|---|

Propanal, 2,3-dihydroxy-, (S)-
Formula C3H6O3, MW 90, CAS# 497-09-6, Entry# 8305
2,3-Dihydroxypropanal #

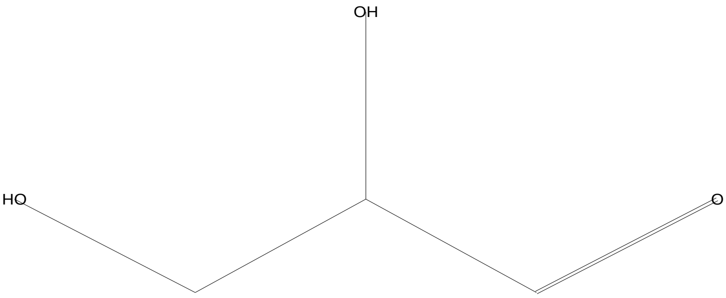


SI 670, RSI 745, mainlib, Entry# 8305, CAS# 497-09-6, Propanal, 2,3-dihydroxy-, (S)-

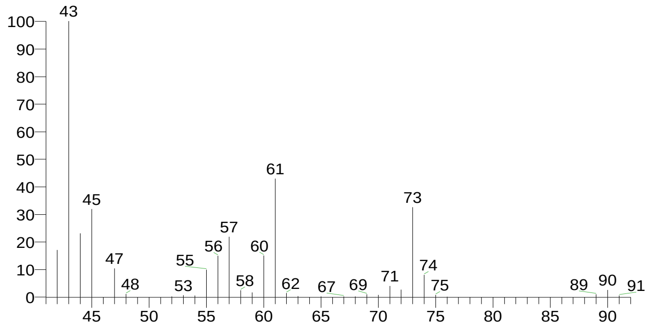


| RT | Compound Name | Area % | Molecular Weight | Molecular Formula | Probability |
|---|---|---|---|---|---|
| 7.46 | Diglycerol | 1.34 | 166 | C6H14O5 | 39.21 |
| 7.46 | Glycerin | 1.34 | 92 | C3H8O3 | 9.69 |
| 7.46 | Glycerin | 1.34 | 92 | C3H8O3 | 9.69 |
| 7.46 | Erythritol | 1.34 | 122 | C4H10O4 | 3.83 |
| 7.46 | Propanal, 2,3-dihydroxy-, (S)- | 1.34 | 90 | C3H6O3 | 3.23 |

There is no signature data to report. Anamika Singh

### Library Search Report

#### Lib. Search Graphics Table

| Compound Structure | Hit Spectrum |
|---|---|

Glycerin
Formula C3H8O3, MW 92, CAS# 56-81-5, Entry# 7605
1,2,3-Propanetriol

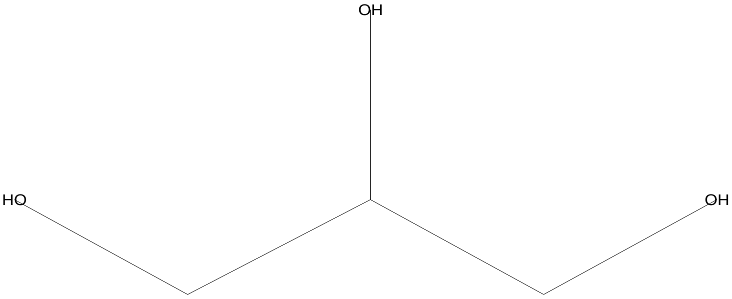


SI 811, RSI 876, replib, Entry# 7605, CAS# 56-81-5, Glycerin

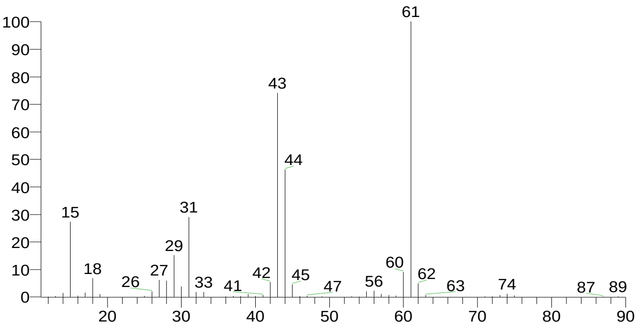


Glycerin
Formula C3H8O3, MW 92, CAS# 56-81-5, Entry# 7604
1,2,3-Propanetriol

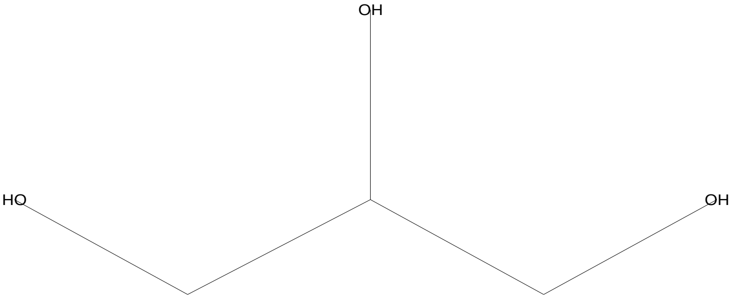


SI 803, RSI 854, replib, Entry# 7604, CAS# 56-81-5, Glycerin

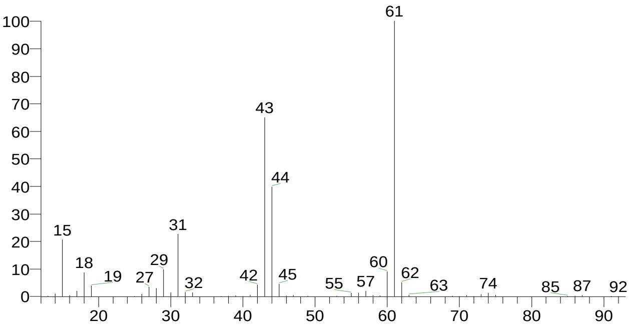


Glycerin
Formula C3H8O3, MW 92, CAS# 56-81-5, Entry# 31153
1,2,3-Propanetriol

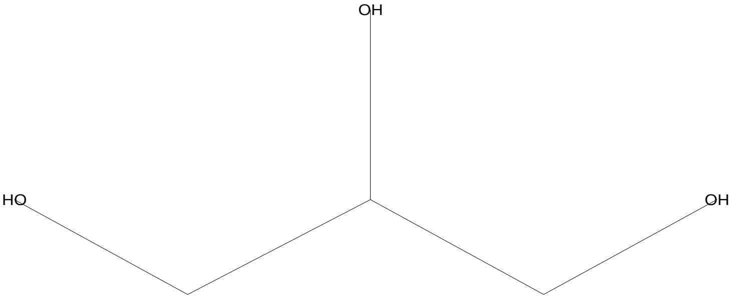


SI 791, RSI 889, mainlib, Entry# 31153, CAS# 56-81-5, Glycerin

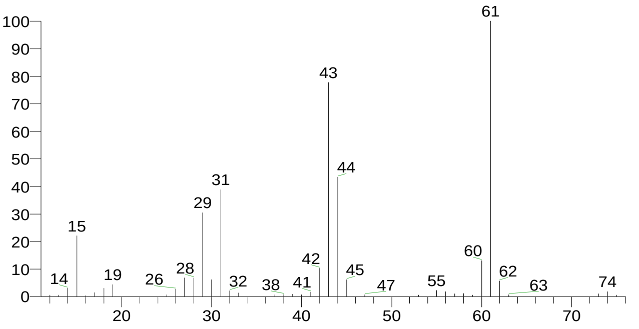


Erythritol
Formula C4H10O4, MW 122, CAS# 149-32-6, Entry# 7617
2(R),3(S)-1,2,3,4-Butanetetrol

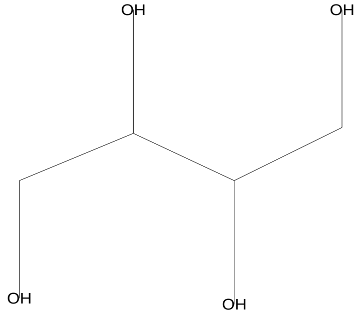


SI 706, RSI 745, replib, Entry# 7617, CAS# 149-32-6, Erythritol

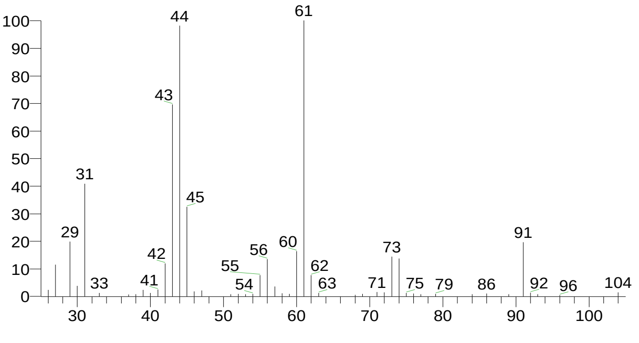


| There is no signature data to report. | Anamika Singh |
|---|---|

### Library Search Report

Compound Structure | Hit Spectrum

N,N-Dimethyl-O-(1-methyl-butyl)-hydroxylamine
Formula C7H17NO, MW 131, CAS# NA, Entry# 31245
2-[(Dimethylamino)oxy]pentane #

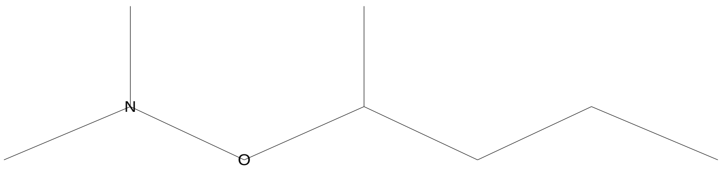


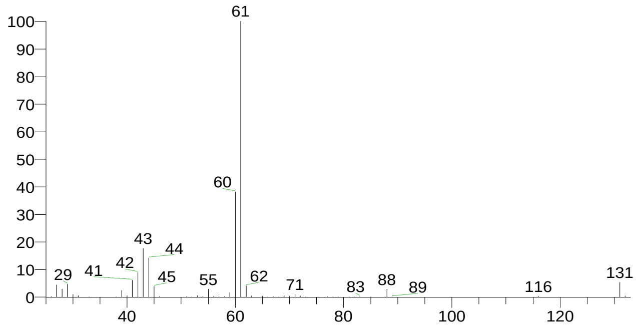


| RT | Compound Name | Area % | Molecular Weight | Molecular Formula | Probability |
|---|---|---|---|---|---|
| 8.21 | Glycerin | 1.41 | 92 | C3H8O3 | 80.22 |
| 8.21 | Glycerin | 1.41 | 92 | C3H8O3 | 80.22 |
| 8.21 | Glycerin | 1.41 | 92 | C3H8O3 | 80.22 |
| 8.21 | Erythritol | 1.41 | 122 | C4H10O4 | 5.45 |
| 8.21 | N,N-Dimethyl-O-(1-methyl-butyl )-hydroxylamine | 1.41 | 131 | C7H17NO | 4.82 |

### Library Search Report

#### Lib. Search Graphics Table

Compound Structure | Hit Spectrum

Glycerin
Formula C3H8O3, MW 92, CAS# 56-81-5, Entry# 7605
1,2,3-Propanetriol

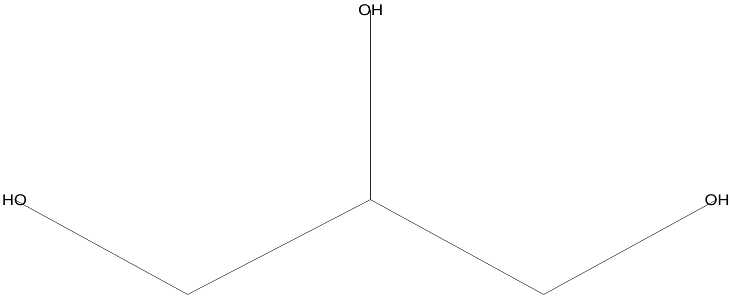


SI 852, RSI 928, replib, Entry# 7605, CAS# 56-81-5, Glycerin

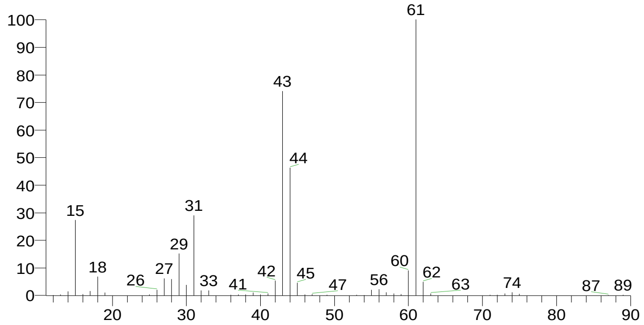


Glycerin
Formula C3H8O3, MW 92, CAS# 56-81-5, Entry# 7604
1,2,3-Propanetriol

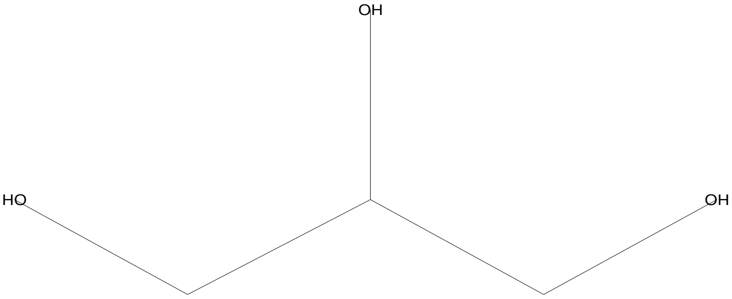


SI 832, RSI 879, replib, Entry# 7604, CAS# 56-81-5, Glycerin

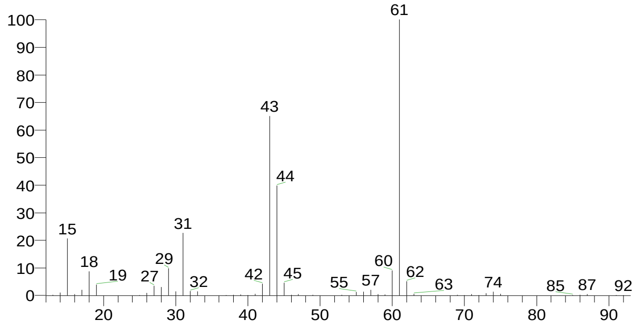


Glycerin
Formula C3H8O3, MW 92, CAS# 56-81-5, Entry# 31153
1,2,3-Propanetriol

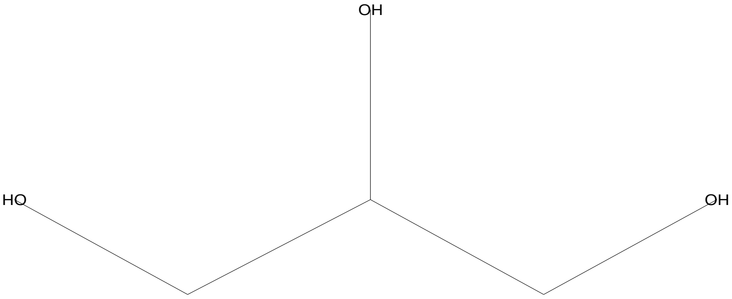


SI 822, RSI 914, mainlib, Entry# 31153, CAS# 56-81-5, Glycerin

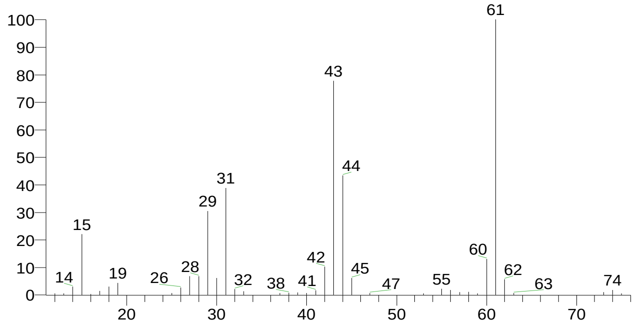


Erythritol
Formula C4H10O4, MW 122, CAS# 149-32-6, Entry# 31179
2(R),3(S)-1,2,3,4-Butanetetrol

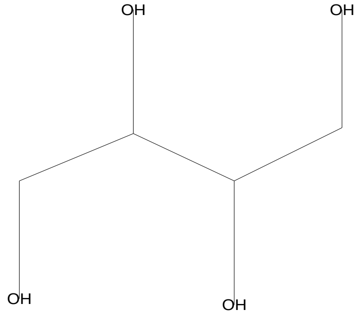


SI 714, RSI 771, mainlib, Entry# 31179, CAS# 149-32-6, Erythritol

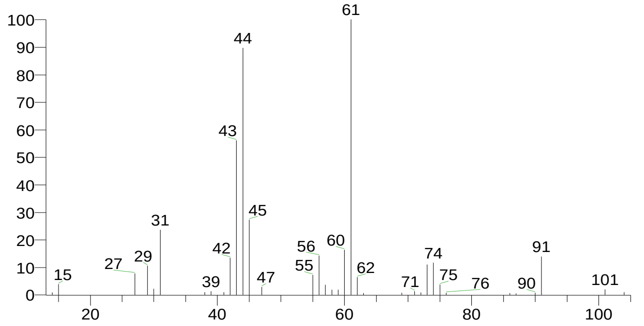

### Library Search Report

Compound Structure

N,N-Dimethyl-O-(1-methyl-butyl)-hydroxylamine
Formula C7H17NO, MW 131, CAS# NA, Entry# 31245
2-[(Dimethylamino)oxy]pentane #

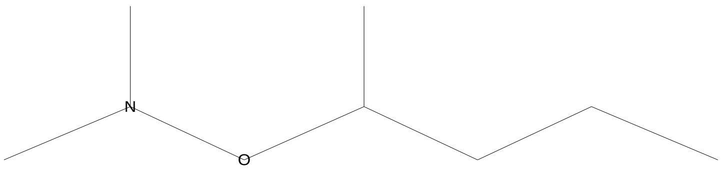


Hit Spectrum

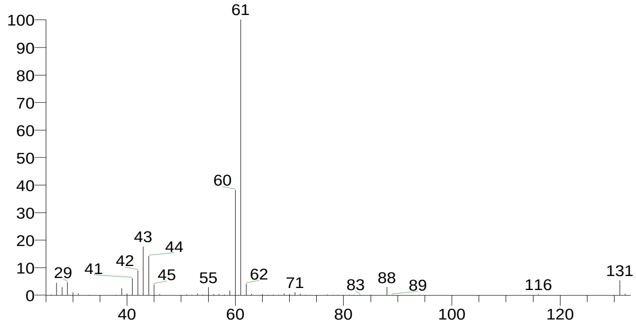


| RT | Compound Name | Area % | Molecular Weight | Molecular Formula | Probability |
|---|---|---|---|---|---|
| 8.59 | Glycerin | 1.13 | 92 | C3H8O3 | 92.54 |
| 8.59 | Glycerin | 1.13 | 92 | C3H8O3 | 92.54 |
| 8.59 | Glycerin | 1.13 | 92 | C3H8O3 | 92.54 |
| 8.59 | Erythritol | 1.13 | 122 | C4H10O4 | 3.29 |
| 8.59 | N,N-Dimethyl-O-(1-methyl-butyl)-hydroxylamine | 1.13 | 131 | C7H17NO | 1.69 |

There is no signature data to report. Anamika Singh

### Library Search Report

#### Lib. Search Graphics Table

| Compound Structure | Hit Spectrum |
|---|---|
| Glycerin<br>Formula C3H8O3, MW 92, CAS# 56-81-5, Entry# 31153<br>1,2,3-Propanetriol | SI 909, RSI 939, mainlib, Entry# 31153, CAS# 56-81-5, Glycerin |

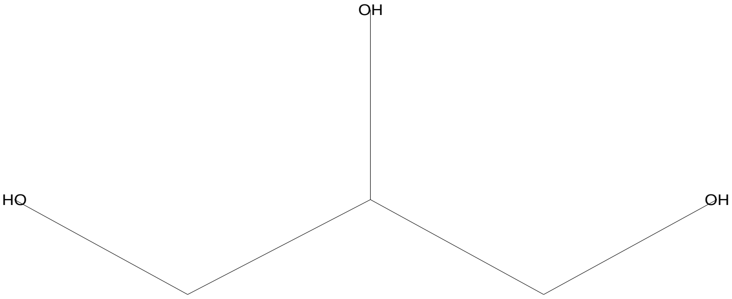


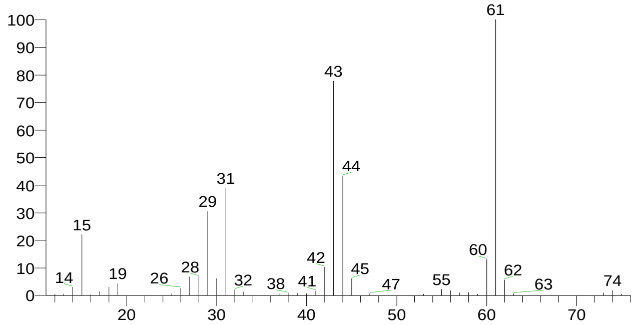


| Compound Structure | Hit Spectrum |
|---|---|
| Glycerin<br>Formula C3H8O3, MW 92, CAS# 56-81-5, Entry# 7604<br>1,2,3-Propanetriol | SI 882, RSI 902, replib, Entry# 7604, CAS# 56-81-5, Glycerin |

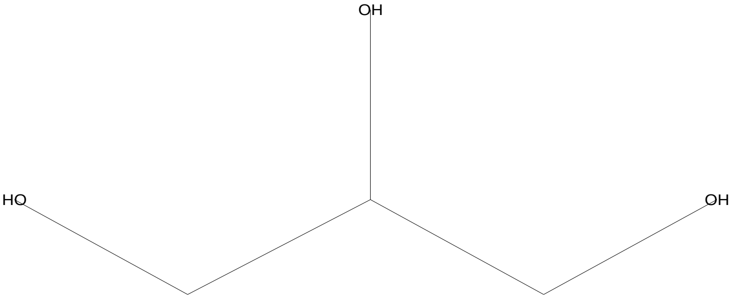


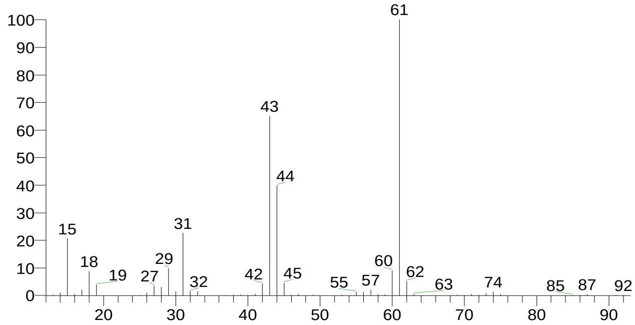


| Compound Structure | Hit Spectrum |
|---|---|
| Glycerin<br>Formula C3H8O3, MW 92, CAS# 56-81-5, Entry# 7605<br>1,2,3-Propanetriol | SI 877, RSI 901, replib, Entry# 7605, CAS# 56-81-5, Glycerin |

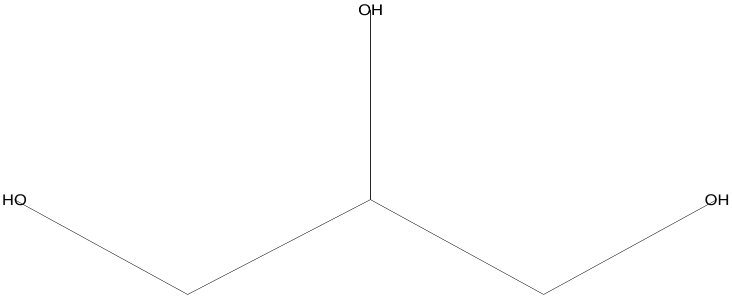


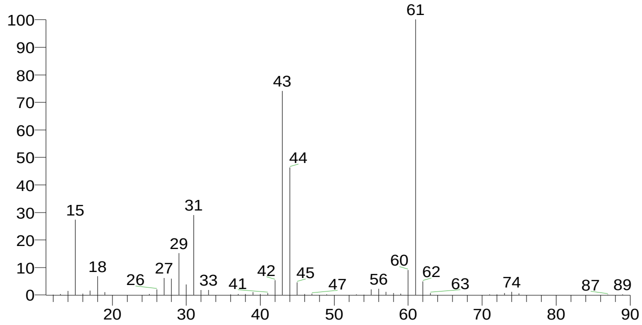


| Compound Structure | Hit Spectrum |
|---|---|
| Erythritol<br>Formula C4H10O4, MW 122, CAS# 149-32-6, Entry# 31179<br>2(R),3(S)-1,2,3,4-Butanetetrol | SI 776, RSI 792, mainlib, Entry# 31179, CAS# 149-32-6, Erythritol |

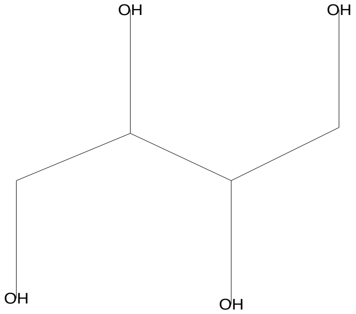


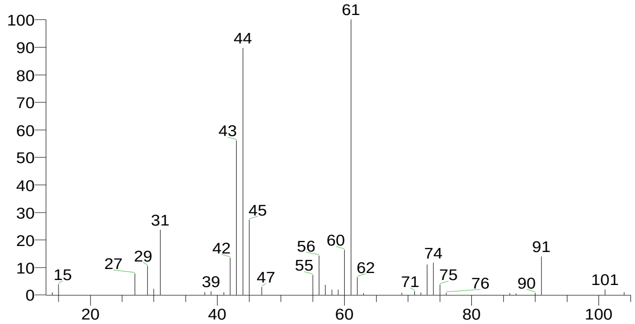

### Library Search Report

Compound Structure

Propane, 2-fluoro-2-methyl-
Formula C4H9F, MW 76, CAS# 353-61-7, Entry# 31132
tert-Butyl Fluoride

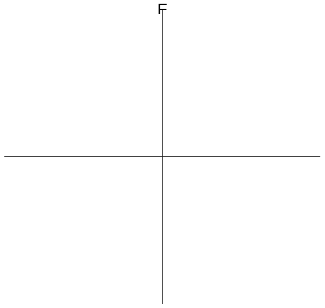


Hit Spectrum

SI 741, RSI 783, mainlib, Entry# 31132, CAS# 353-61-7, Propane, 2-fluoro-2-methyl-

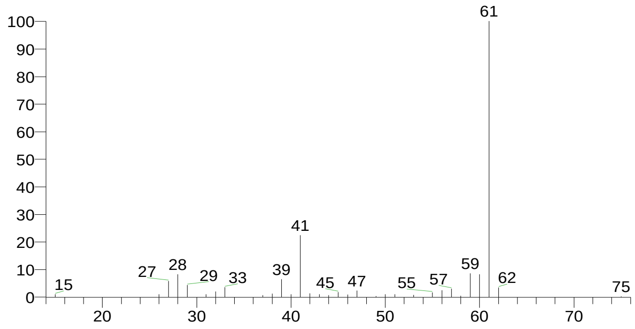


| RT | Compound Name | Area % | Molecular Weight | Molecular Formula | Probability |
|---|---|---|---|---|---|
| 9.33 | Glycerin | 16.94 | 92 | C3H8O3 | 94.03 |
| 9.33 | Glycerin | 16.94 | 92 | C3H8O3 | 94.03 |
| 9.33 | Glycerin | 16.94 | 92 | C3H8O3 | 94.03 |
| 9.33 | Erythritol | 16.94 | 122 | C4H10O4 | 3.70 |
| 9.33 | Propane, 2-fluoro-2-methyl- | 16.94 | 76 | C4H9F | 0.92 |

### Library Search Report

#### Lib. Search Graphics Table

Compound Structure | Hit Spectrum

Desulphosinigrin
Formula C10H17NO6S, MW 279, CAS# 5115-81-1, Entry# 30961
1-S-[(1E)-N-Hydroxy-3-butenimidoyl]-1-thiohexopyranose #

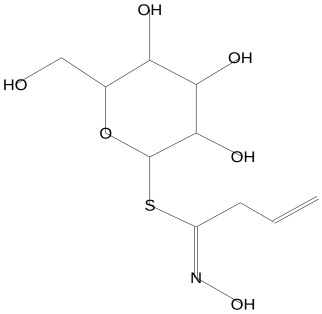


SI 645, RSI 666, mainlib, Entry# 30961, CAS# 5115-81-1, Desulphosinigrin

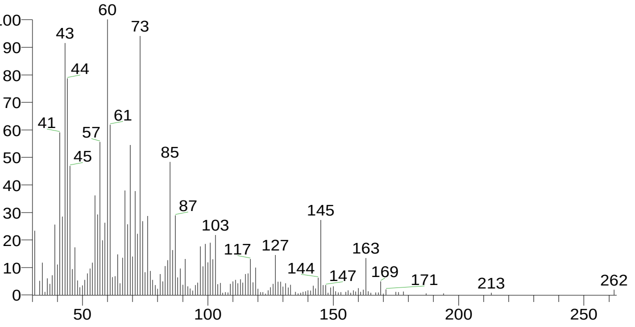


D-Glucose, cyclic 1,2-ethanediyl mercaptal, pentaacetate
Formula C18H26O10S2, MW 466, CAS# 17429-98-0, Entry# 78224
D-Glucose, cyclic ethylene mercaptal, pentaacetate

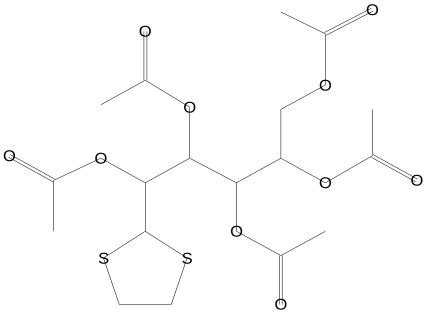


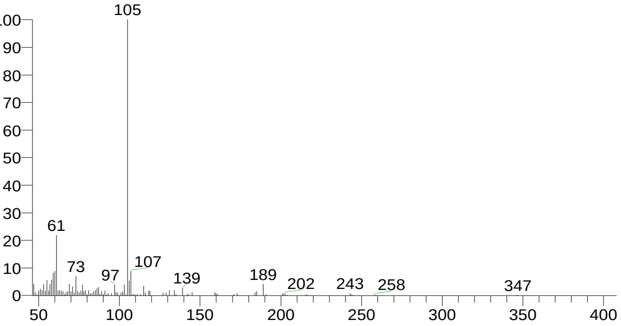


á-D-Glucopyranose, 4-O-á-D-galactopyranosyl-
Formula C12H22O11, MW 342, CAS# 5965-66-2, Entry# 9475
Lactose, á-

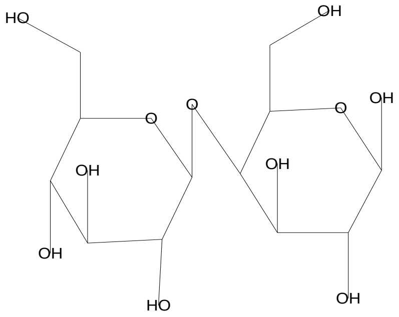


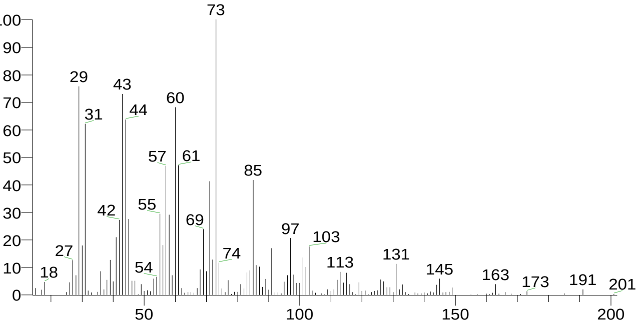


3-tert-Butyl-5-chloro-2-hydroxybenzophenone
Formula C17H17ClO2, MW 288, CAS# 52196-47-1, Entry# 77947
(3-tert-Butyl-5-chloro-2-hydroxyphenyl)(phenyl)methanone #

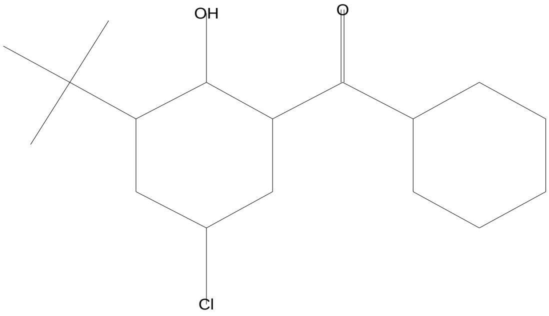


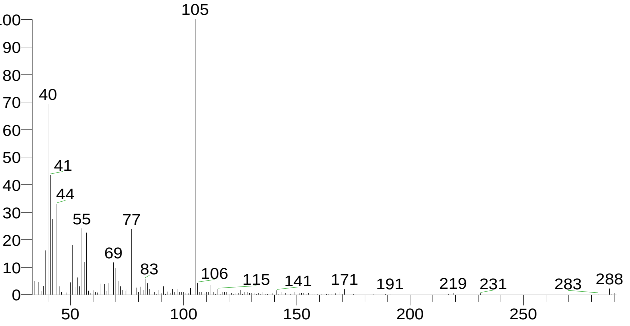

### Library Search Report

Compound Structure | Hit Spectrum

á-D-Glucopyranose, 4-O-á-D-galactopyranosyl-
Formula C12H22O11, MW 342, CAS# 5965-66-2, Entry# 40630
Lactose, á-

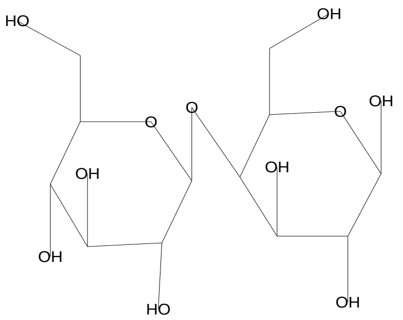


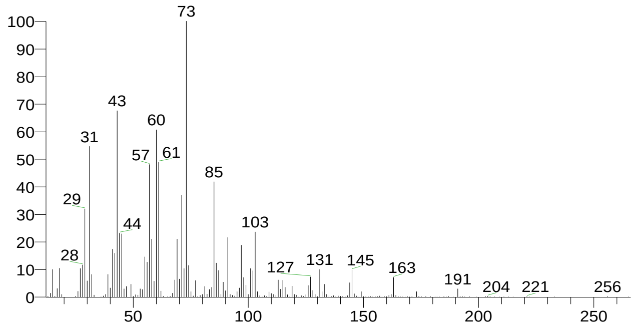


| RT | Compound Name | Area % | Molecular Weight | Molecular Formula | Probability |
|---|---|---|---|---|---|
| 10.35 | Desulphosinigrin | 1.36 | 279 | C10H17NO6S | 29.74 |
| 10.35 | D-Glucose, cyclic 1,2-ethanediyl mercaptal, pentaacetate | 1.36 | 466 | C18H26O10S2 | 19.21 |
| 10.35 | á-D-Glucopyranose, 4-O-á-D-galactopyranosyl- | 1.36 | 342 | C12H22O11 | 15.10 |
| 10.35 | á-D-Glucopyranose, 4-O-á-D-galactopyranosyl- | 1.36 | 342 | C12H22O11 | 15.10 |
| 10.35 | 3-tert-Butyl-5-chloro-2-hydroxyb enzophenone | 1.36 | 288 | C17H17ClO2 | 4.27 |

There is no signature data to report. Anamika Singh

### Library Search Report

#### Lib. Search Graphics Table

Compound Structure | Hit Spectrum

Dodecane
Formula C12H26, MW 170, CAS# 112-40-3, Entry# 24230
n-Dodecane

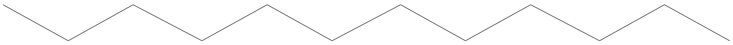

SI 857, RSI 911, mainlib, Entry# 24230, CAS# 112-40-3, Dodecane

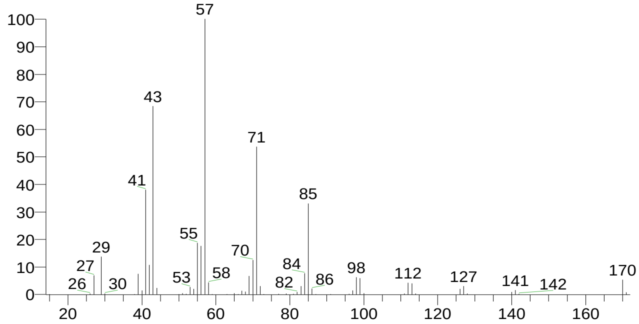


Dodecane
Formula C12H26, MW 170, CAS# 112-40-3, Entry# 6083
n-Dodecane

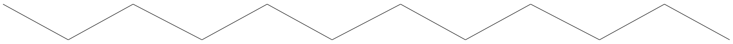

SI 856, RSI 914, replib, Entry# 6083, CAS# 112-40-3, Dodecane

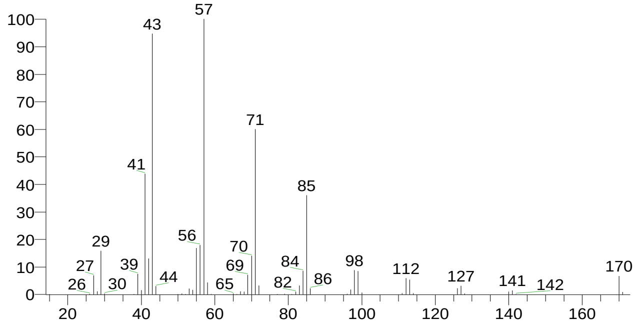


Dodecane
Formula C12H26, MW 170, CAS# 112-40-3, Entry# 2294
n-Dodecane

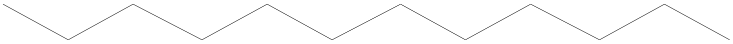

SI 850, RSI 933, replib, Entry# 2294, CAS# 112-40-3, Dodecane

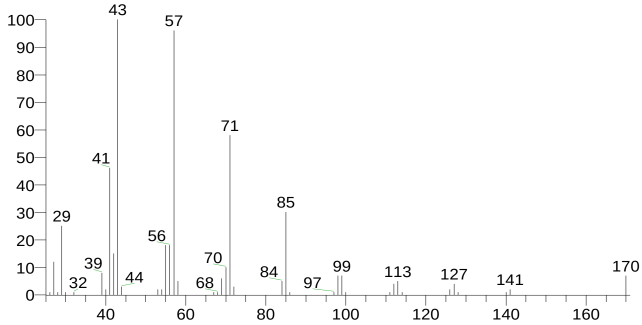


Tridecane
Formula C13H28, MW 184, CAS# 629-50-5, Entry# 5965
n-Tridecane

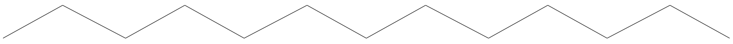

SI 842, RSI 915, replib, Entry# 5965, CAS# 629-50-5, Tridecane

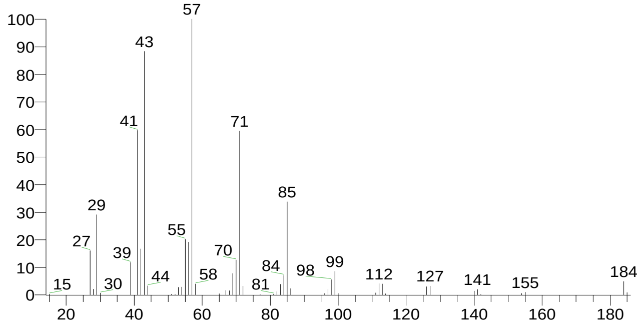

### Library Search Report

| Compound Structure | Hit Spectrum |
|---|---|
| Dodecane<br>Formula C12H26, MW 170, CAS# 112-40-3, Entry# 6084<br>n-Dodecane | SI 840, RSI 901, replib, Entry# 6084, CAS# 112-40-3, Dodecane |

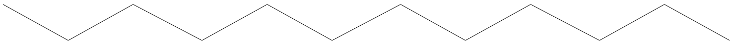

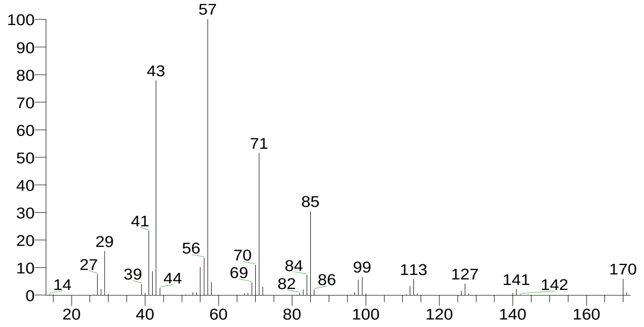


| RT | Compound Name | Area % | Molecular Weight | Molecular Formula | Probability |
|---|---|---|---|---|---|
| 10.99 | Dodecane | 6.33 | 170 | C12H26 | 20.04 |
| 10.99 | Dodecane | 6.33 | 170 | C12H26 | 20.04 |
| 10.99 | Dodecane | 6.33 | 170 | C12H26 | 20.04 |
| 10.99 | Dodecane | 6.33 | 170 | C12H26 | 20.04 |
| 10.99 | Tridecane | 6.33 | 184 | C13H28 | 12.14 |

There is no signature data to report. Anamika Singh

### Library Search Report

#### Lib. Search Graphics Table

| Compound Structure | Hit Spectrum |
|---|---|

2-Acetonyl-9-[3-deoxy-á-d-ribouranosyl]hypoxanthine
Formula C13H16N4O5, MW 308, CAS# 132121-67-6, Entry# 7684
$:28MBQOESUAWPYLQO-UHFFFAOYSA-N

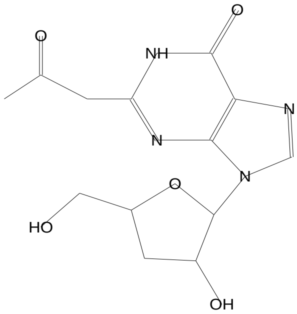


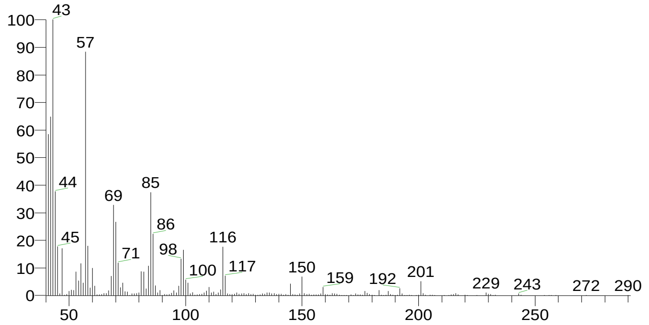


2-Vinyl-9-[3-deoxy-á-d-ribofuranosyl]hypoxanthine
Formula C12H14N4O4, MW 278, CAS# 132121-65-4, Entry# 23904
$:28LYWVBVVDTLPXMF-UHFFFAOYSA-N

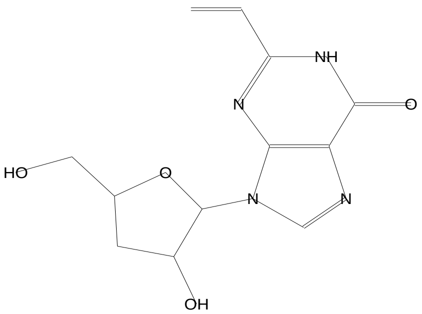


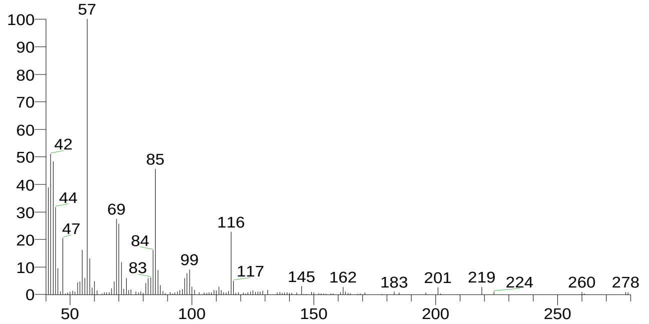


1-Dodecanol, 3,7,11-trimethyl-
Formula C15H32O, MW 228, CAS# 6750-34-1, Entry# 6352
Hexa-hydro-farnesol

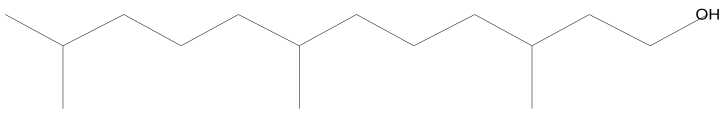


SI 654, RSI 678, replib, Entry# 6352, CAS# 6750-34-1, 1-Dodecanol, 3,7,11-trimethyl-

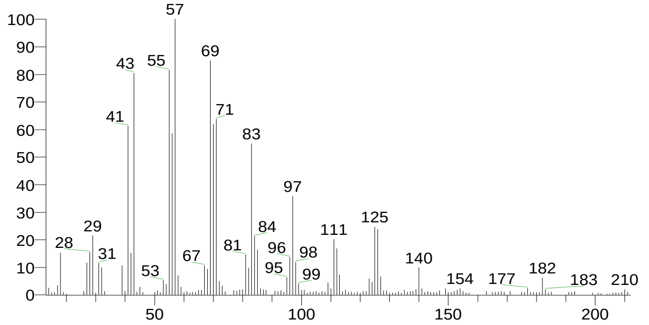


tert-Hexadecanethiol
Formula C16H34S, MW 258, CAS# 25360-09-2, Entry# 25117
1,1-Dimethyltetradecyl hydrosulfide #

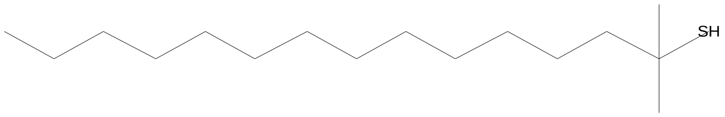


SI 654, RSI 671, mainlib, Entry# 25117, CAS# 25360-09-2, tert-Hexadecanethiol

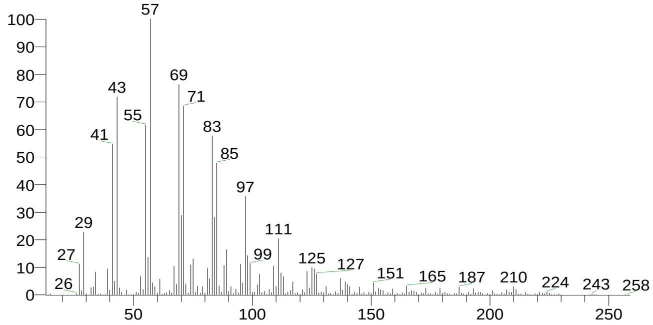


There is no signature data to report. Anamika Singh

### Library Search Report

Compound Structure

1-Hexadecanol, 2-methyl-
Formula C17H36O, MW 256, CAS# 2490-48-4, Entry# 24113
2-Methylhexadecan-1-ol

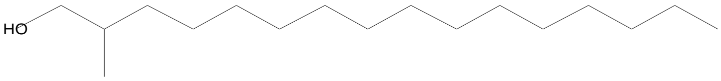


Hit Spectrum

SI 649, RSI 664, mainlib, Entry# 24113, CAS# 2490-48-4, 1-Hexadecanol, 2-methyl-

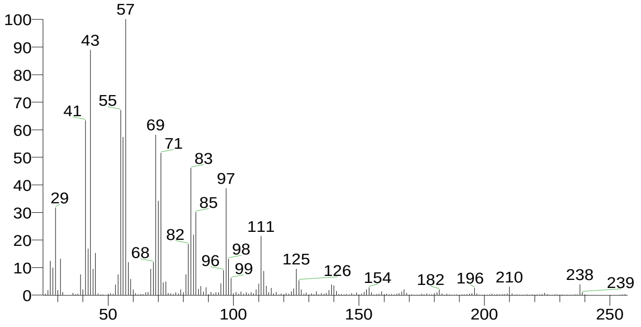


| RT | Compound Name | Area % | Molecular Weight | Molecular Formula | Probability |
|---|---|---|---|---|---|
| 11.42 | 2-Acetonyl-9-[3-deoxy-á-d-ribouranosyl]hypoxanthine | 1.13 | 308 | C13H16N4O5 | 10.93 |
| 11.42 | 2-Vinyl-9-[3-deoxy-á-d-ribofuranosyl]hypoxanthine | 1.13 | 278 | C12H14N4O4 | 5.96 |
| 11.42 | tert-Hexadecanethiol | 1.13 | 258 | C16H34S | 4.80 |
| 11.42 | 1-Dodecanol, 3,7,11-trimethyl- | 1.13 | 228 | C15H32O | 4.80 |
| 11.42 | 1-Hexadecanol, 2-methyl- | 1.13 | 256 | C17H36O | 3.87 |

There is no signature data to report. Anamika Singh

### Library Search Report

#### Lib. Search Graphics Table

| Compound Structure | Hit Spectrum |
| --- | --- |

á-D-Glucopyranose, 4-O-á-D-galactopyranosyl-
Formula C12H22O11, MW 342, CAS# 5965-66-2, Entry# 9475
Lactose, á-

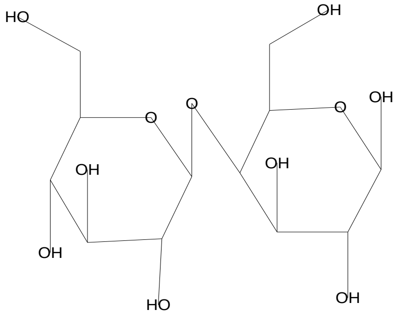


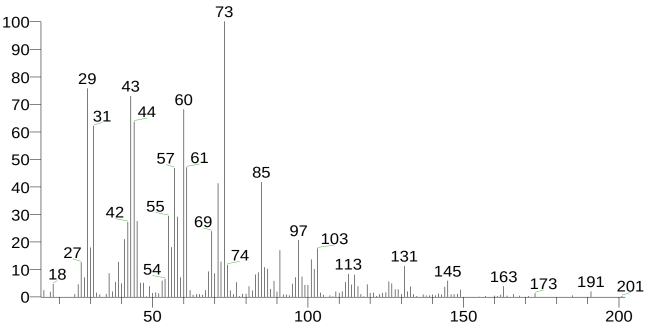


2-Myristynoyl pantetheine
Formula C25H44N2O5S, MW 484, CAS# NA, Entry# 4545

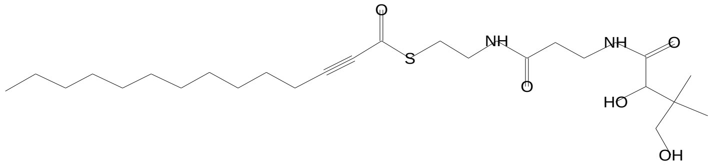


SI 677, RSI 682, mainlib, Entry# 4545, CAS# NA, 2-Myristynoyl pantetheine

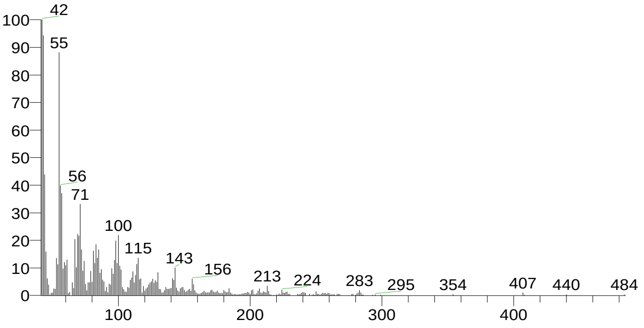


d-Glycero-d-galacto-heptose
Formula C7H14O7, MW 210, CAS# 77770-51-5, Entry# 9209
Heptose #

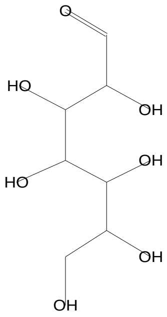


SI 673, RSI 704, mainlib, Entry# 9209, CAS# 77770-51-5, d-Glycero-d-galacto-heptose

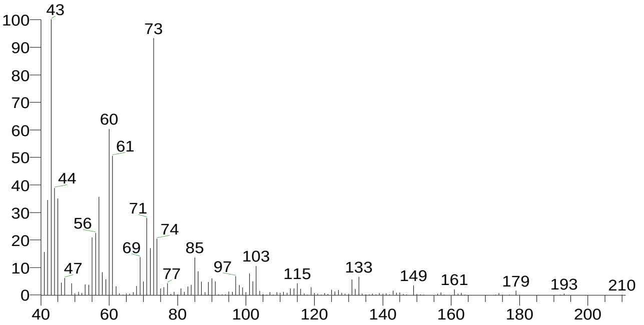


d-Glycero-d-ido-heptose
Formula C7H14O7, MW 210, CAS# NA, Entry# 9206
Heptose #

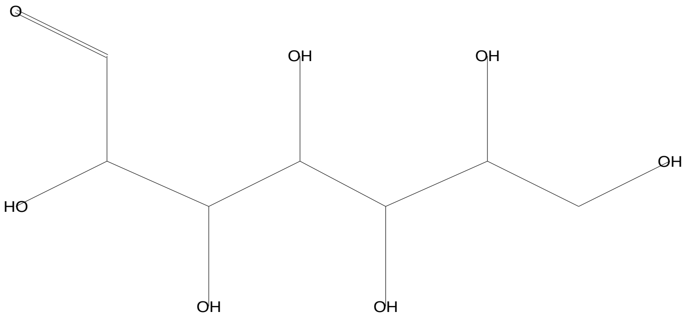


SI 671, RSI 703, mainlib, Entry# 9206, CAS# NA, d-Glycero-d-ido-heptose

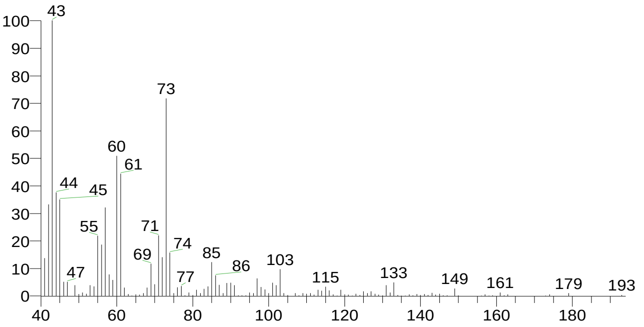

### Library Search Report

Compound Structure | Hit Spectrum

á-D-Glucopyranose, 4-O-á-D-galactopyranosyl-
Formula C12H22O11, MW 342, CAS# 5965-66-2, Entry# 40630
Lactose, á-

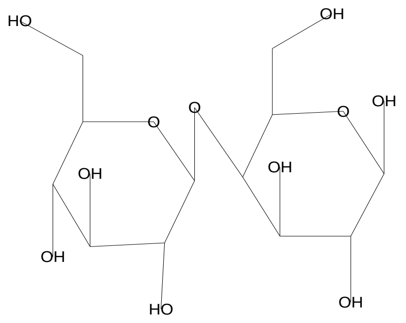


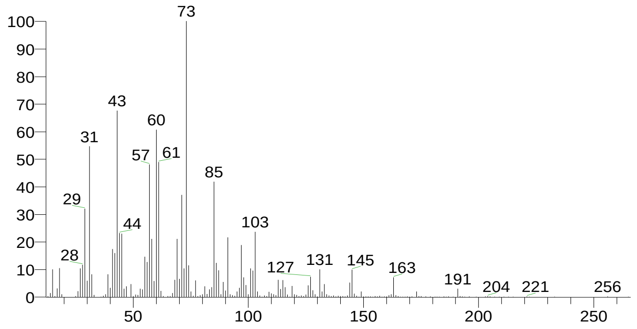


| RT | Compound Name | Area % | Molecular Weight | Molecular Formula | Probability |
|---|---|---|---|---|---|
| 12.26 | á-D-Glucopyranose, 4-O-á-D-galactopyranosyl- | 1.53 | 342 | C12H22O11 | 26.89 |
| 12.26 | á-D-Glucopyranose, 4-O-á-D-galactopyranosyl- | 1.53 | 342 | C12H22O11 | 26.89 |
| 12.26 | 2-Myristynoyl pantetheine | 1.53 | 484 | C25H44N2O5S | 7.33 |
| 12.26 | d-Glycero-d-galacto-heptose | 1.53 | 210 | C7H14O7 | 6.19 |
| 12.26 | d-Glycero-d-ido-heptose | 1.53 | 210 | C7H14O7 | 5.71 |

There is no signature data to report. Anamika Singh

### Library Search Report

#### Lib. Search Graphics Table

| Compound Structure | Hit Spectrum |
|---|---|

á-D-Glucopyranose, 4-O-á-D-galactopyranosyl-
Formula C12H22O11, MW 342, CAS# 5965-66-2, Entry# 9475
Lactose, á-

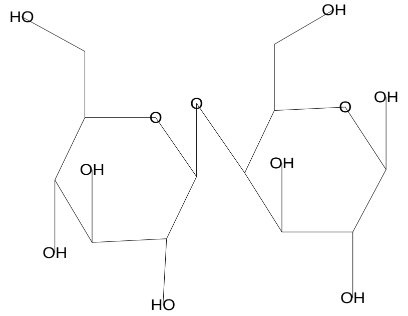


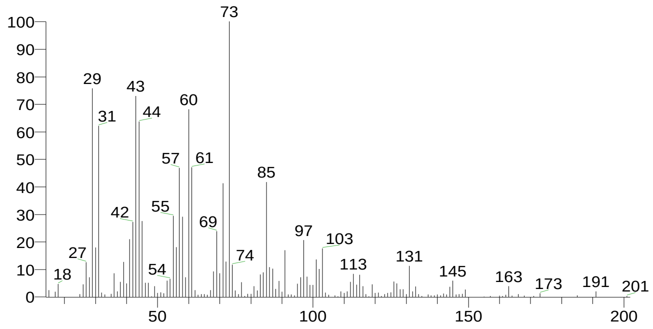


2-Deoxy-D-galactose
Formula C6H12O5, MW 164, CAS# 1949-89-9, Entry# 1474
2-Desoxy-D-galactose

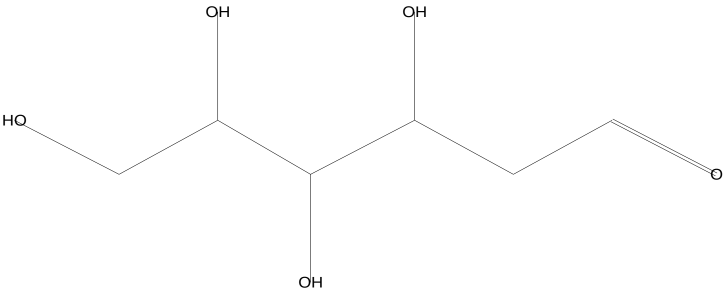


SI 719, RSI 750, mainlib, Entry# 1474, CAS# 1949-89-9, 2-Deoxy-D-galactose

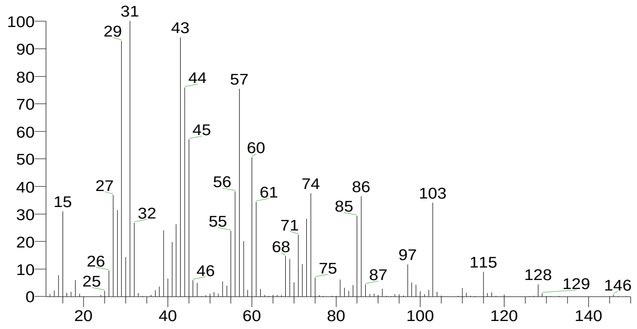


á-D-Glucopyranose, 4-O-á-D-galactopyranosyl-
Formula C12H22O11, MW 342, CAS# 5965-66-2, Entry# 40630
Lactose, á-

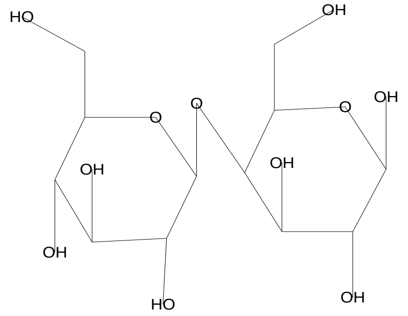


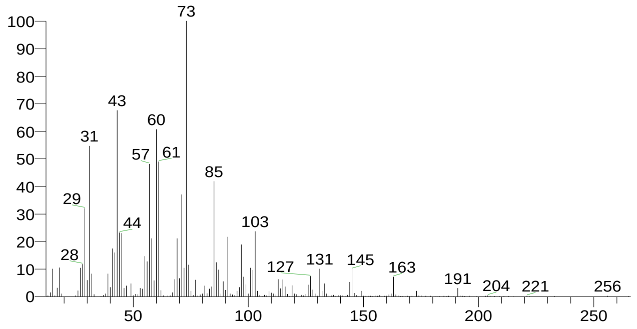


d-Glycero-d-ido-heptose
Formula C7H14O7, MW 210, CAS# NA, Entry# 9206
Heptose #

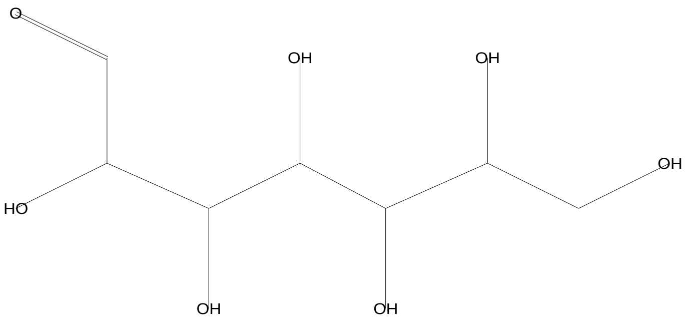


SI 716, RSI 729, mainlib, Entry# 9206, CAS# NA, d-Glycero-d-ido-heptose

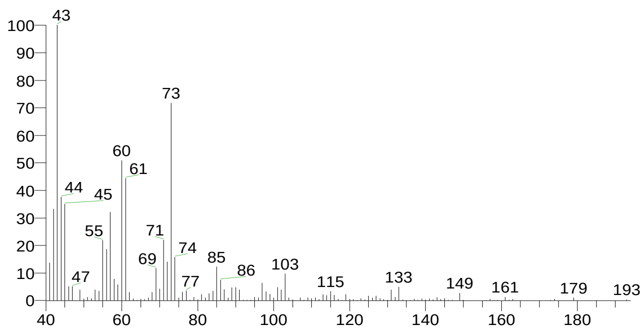


There is no signature data to report. Anamika Singh

### Library Search Report

| Compound Structure | Hit Spectrum |
|---|---|
| d-Glycero-d-galacto-heptose<br>Formula C7H14O7, MW 210, CAS# 77770-51-5, Entry# 9209<br>Heptose # | SI 714, RSI 726, mainlib, Entry# 9209, CAS# 77770-51-5, d-Glycero-d-galacto-heptose |

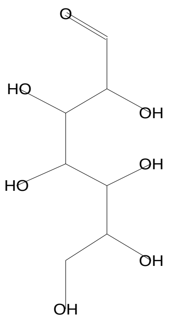


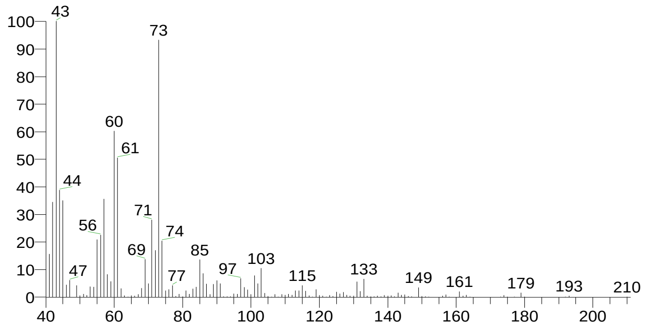


| RT | Compound Name | Area % | Molecular Weight | Molecular Formula | Probability |
|---|---|---|---|---|---|
| 12.58 | á-D-Glucopyranose, 4-O-á-D-galactopyranosyl- | 3.04 | 342 | C12H22O11 | 25.84 |
| 12.58 | á-D-Glucopyranose, 4-O-á-D-galactopyranosyl- | 3.04 | 342 | C12H22O11 | 25.84 |
| 12.58 | 2-Deoxy-D-galactose | 3.04 | 164 | C6H12O5 | 7.04 |
| 12.58 | d-Glycero-d-ido-heptose | 3.04 | 210 | C7H14O7 | 6.22 |
| 12.58 | d-Glycero-d-galacto-heptose | 3.04 | 210 | C7H14O7 | 5.74 |

### Library Search Report

#### Lib. Search Graphics Table

| Compound Structure | Hit Spectrum |
|---|---|

á-D-Glucopyranose, 4-O-á-D-galactopyranosyl-
Formula C12H22O11, MW 342, CAS# 5965-66-2, Entry# 9475
Lactose, á-

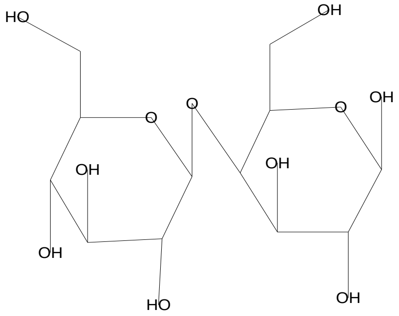


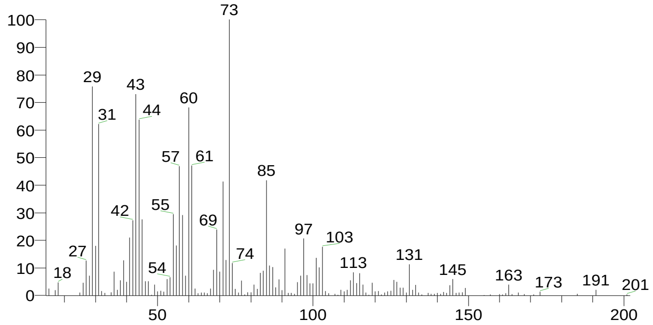


Ribitol
Formula C5H12O5, MW 152, CAS# 488-81-3, Entry# 44657
Adonit

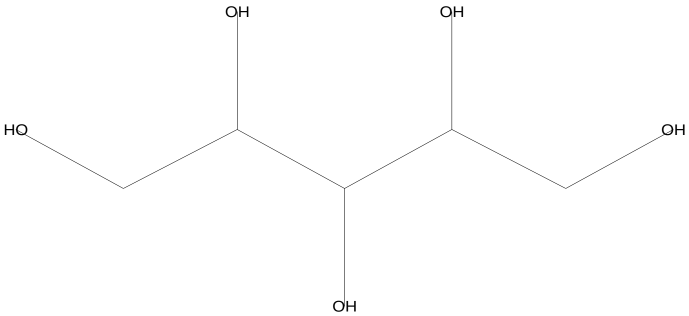


SI 733, RSI 782, mainlib, Entry# 44657, CAS# 488-81-3, Ribitol

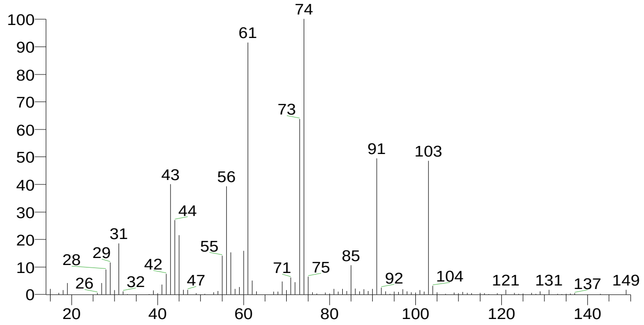


d-Glycero-d-ido-heptose
Formula C7H14O7, MW 210, CAS# NA, Entry# 9206
Heptose #

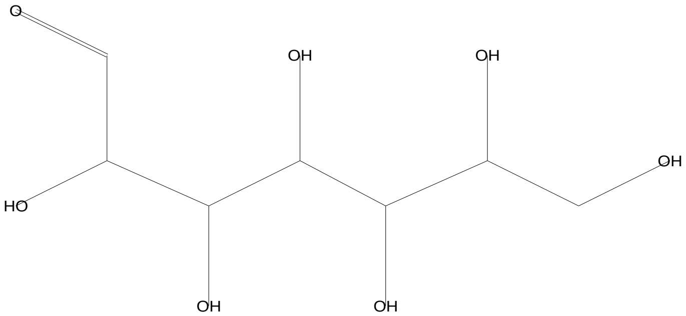


SI 730, RSI 750, mainlib, Entry# 9206, CAS# NA, d-Glycero-d-ido-heptose

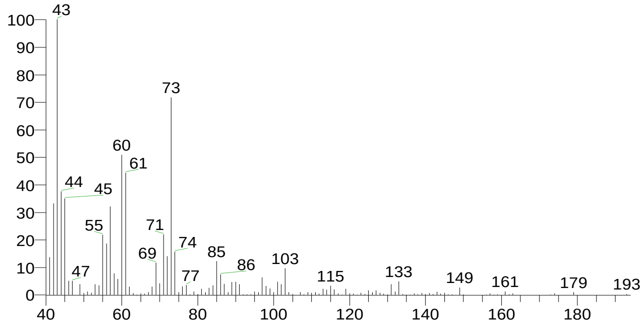


L-Glucose
Formula C6H12O6, MW 180, CAS# 921-60-8, Entry# 40628
Hexopyranose #

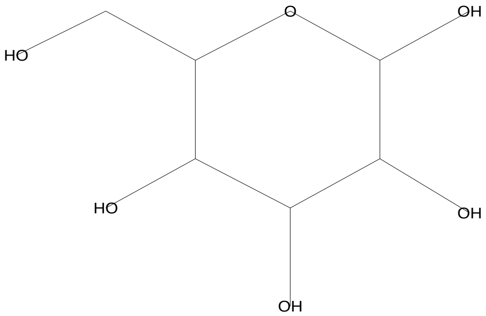


SI 726, RSI 744, mainlib, Entry# 40628, CAS# 921-60-8, L-Glucose

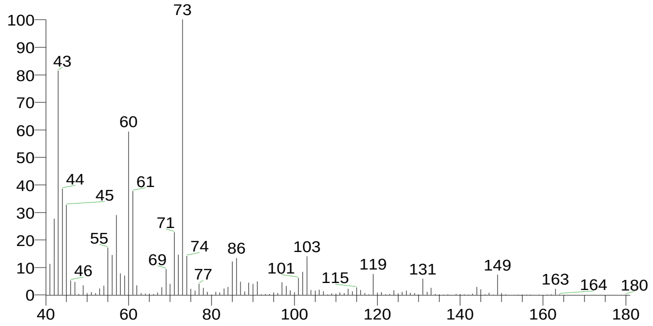

### Library Search Report

| Compound Structure | Hit Spectrum |
|---|---|
| d-Glycero-d-galacto-heptose<br>Formula C7H14O7, MW 210, CAS# 77770-51-5, Entry# 9209<br>Heptose # | SI 725, RSI 743, mainlib, Entry# 9209, CAS# 77770-51-5, d-Glycero-d-galacto-heptose |

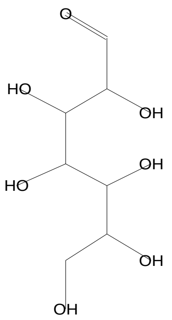


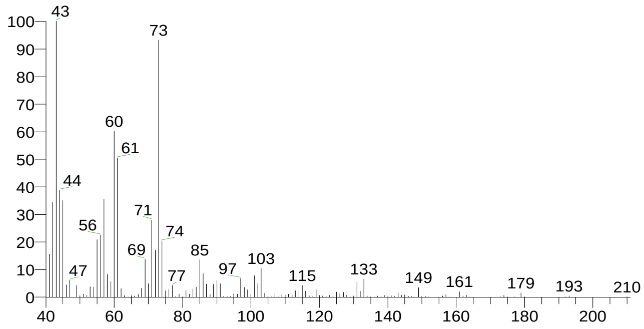


| RT | Compound Name | Area % | Molecular Weight | Molecular Formula | Probability |
|---|---|---|---|---|---|
| 13.64 | á-D-Glucopyranose, 4-O-á-D-galactopyranosyl- | 2.86 | 342 | C12H22O11 | 13.56 |
| 13.64 | Ribitol | 2.86 | 152 | C5H12O5 | 9.58 |
| 13.64 | d-Glycero-d-ido-heptose | 2.86 | 210 | C7H14O7 | 8.46 |
| 13.64 | L-Glucose | 2.86 | 180 | C6H12O6 | 7.15 |
| 13.64 | d-Glycero-d-galacto-heptose | 2.86 | 210 | C7H14O7 | 6.87 |

### Library Search Report

#### Lib. Search Graphics Table

Compound Structure | Hit Spectrum

Tetradecane
Formula C14H30, MW 198, CAS# 629-59-4, Entry# 6116
n-Tetradecane

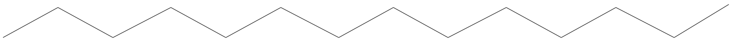

SI 881, RSI 960, replib, Entry# 6116, CAS# 629-59-4, Tetradecane

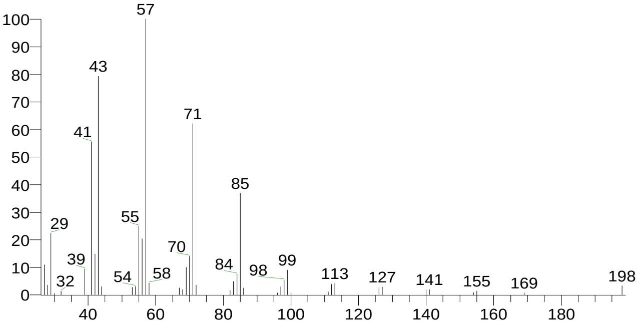


Tetradecane
Formula C14H30, MW 198, CAS# 629-59-4, Entry# 6115
n-Tetradecane

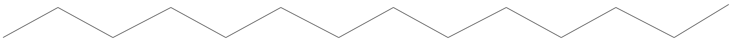

SI 861, RSI 918, replib, Entry# 6115, CAS# 629-59-4, Tetradecane

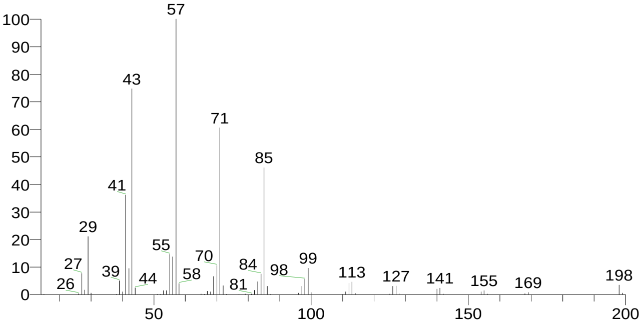


Hexadecane
Formula C16H34, MW 226, CAS# 544-76-3, Entry# 6168
n-Cetane

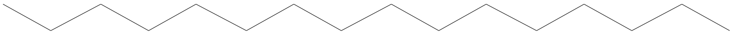

SI 852, RSI 924, replib, Entry# 6168, CAS# 544-76-3, Hexadecane

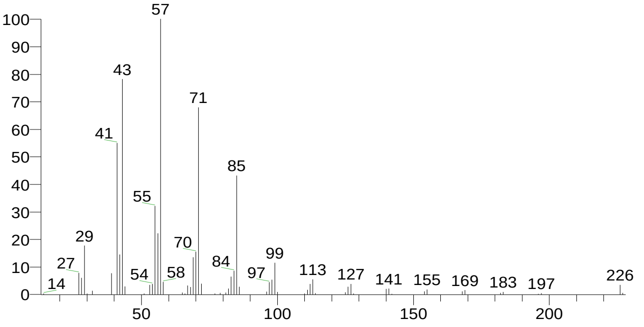


Tetradecane
Formula C14H30, MW 198, CAS# 629-59-4, Entry# 24296
n-Tetradecane

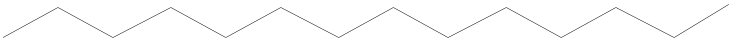

SI 846, RSI 906, mainlib, Entry# 24296, CAS# 629-59-4, Tetradecane

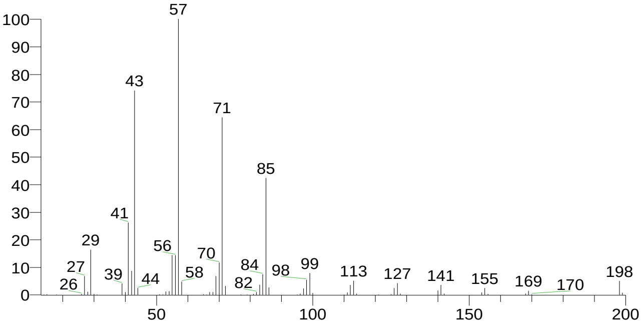


There is no signature data to report. Anamika Singh

### Library Search Report

| Compound Structure | Hit Spectrum |
|---|---|

Tridecane
Formula C13H28, MW 184, CAS# 629-50-5, Entry# 5965
n-Tridecane

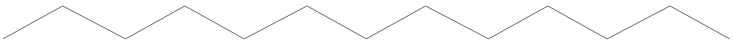

SI 845, RSI 924, replib, Entry# 5965, CAS# 629-50-5, Tridecane

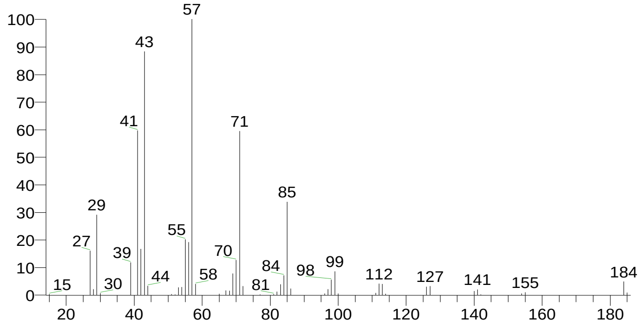


| RT | Compound Name | Area % | Molecular Weight | Molecular Formula | Probability |
|---|---|---|---|---|---|
| 14.84 | Tetradecane | 4.49 | 198 | C14H30 | 34.67 |
| 14.84 | Tetradecane | 4.49 | 198 | C14H30 | 34.67 |
| 14.84 | Tetradecane | 4.49 | 198 | C14H30 | 34.67 |
| 14.84 | Hexadecane | 4.49 | 226 | C16H34 | 9.82 |
| 14.84 | Tridecane | 4.49 | 184 | C13H28 | 7.52 |

There is no signature data to report. Anamika Singh

### Library Search Report

#### Lib. Search Graphics Table

| Compound Structure | Hit Spectrum |
| --- | --- |

2-Myristynoyl pantetheine
Formula C25H44N2O5S, MW 484, CAS# NA, Entry# 4545

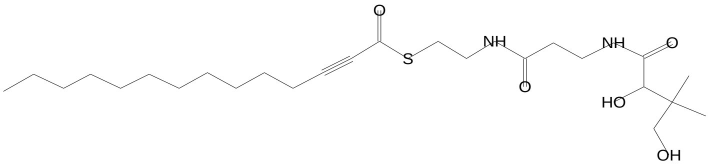


SI 682, RSI 686, mainlib, Entry# 4545, CAS# NA, 2-Myristynoyl pantetheine

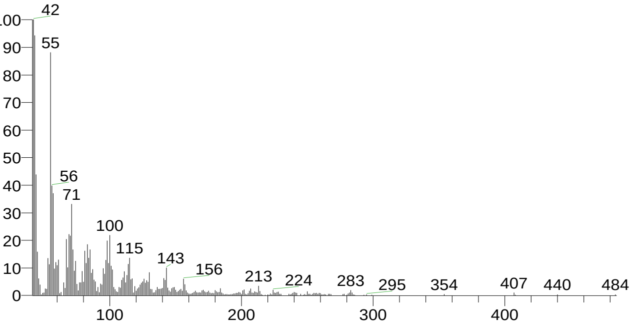


Tetradecane, 2,6,10-trimethyl-
Formula C17H36, MW 240, CAS# 14905-56-7, Entry# 24375
2,6,10-Trimethyltetradecane

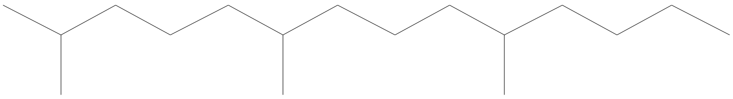

SI 677, RSI 786, mainlib, Entry# 24375, CAS# 14905-56-7, Tetradecane, 2,6,10-trimethyl-

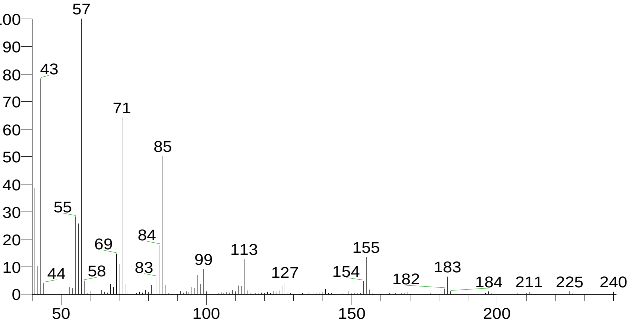


á-D-Glucopyranose, 4-O-á-D-galactopyranosyl-
Formula C12H22O11, MW 342, CAS# 5965-66-2, Entry# 40630
Lactose, á-

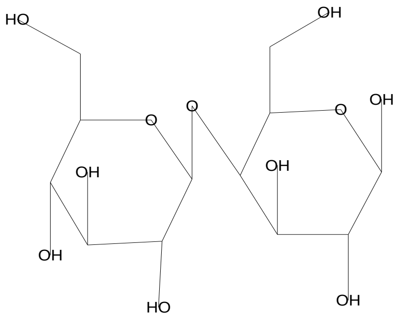


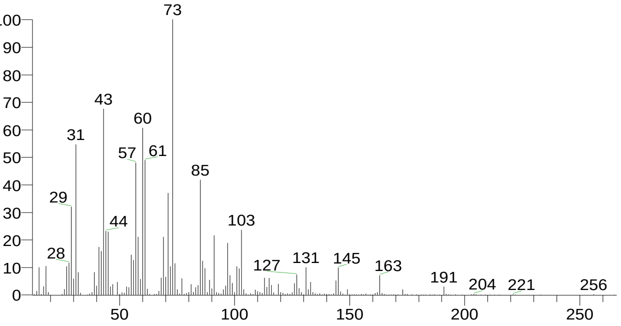


1-Hexadecanol, 2-methyl-
Formula C17H36O, MW 256, CAS# 2490-48-4, Entry# 24113
2-Methylhexadecan-1-ol

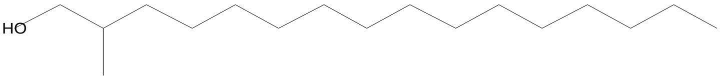


SI 663, RSI 673, mainlib, Entry# 24113, CAS# 2490-48-4, 1-Hexadecanol, 2-methyl-

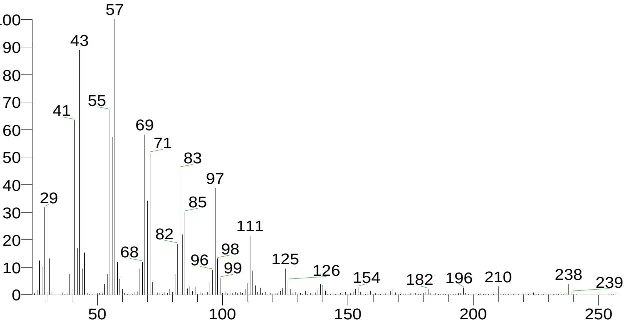


There is no signature data to report. Anamika Singh

### Library Search Report

Compound Structure

1-Dodecanol, 3,7,11-trimethyl-
Formula C15H32O, MW 228, CAS# 6750-34-1, Entry# 6352
Hexa-hydro-farnesol

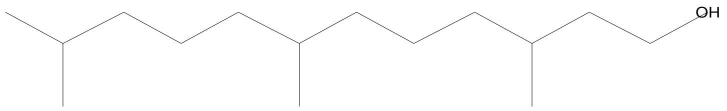


Hit Spectrum

SI 659, RSI 708, replib, Entry# 6352, CAS# 6750-34-1, 1-Dodecanol, 3,7,11-trimethyl-

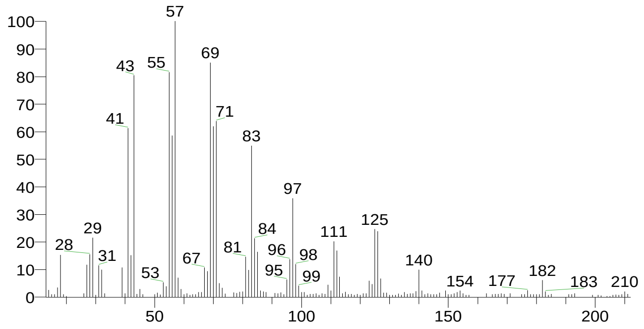


| RT | Compound Name | Area % | Molecular Weight | Molecular Formula | Probability |
|---|---|---|---|---|---|
| 15.97 | 2-Myristynoyl pantetheine | 1.03 | 484 | C25H44N2O5S | 11.23 |
| 15.97 | Tetradecane, 2,6,10-trimethyl- | 1.03 | 240 | C17H36 | 9.05 |
| 15.97 | á-D-Glucopyranose, 4-O-á-D-galactopyranosyl- | 1.03 | 342 | C12H22O11 | 6.21 |
| 15.97 | 1-Hexadecanol, 2-methyl- | 1.03 | 256 | C17H36O | 5.48 |
| 15.97 | 1-Dodecanol, 3,7,11-trimethyl- | 1.03 | 228 | C15H32O | 4.63 |

There is no signature data to report. Anamika Singh

### Library Search Report

#### Lib. Search Graphics Table

| Compound Structure | Hit Spectrum |
| --- | --- |

2-Myristynoyl pantetheine
Formula C25H44N2O5S, MW 484, CAS# NA, Entry# 4545

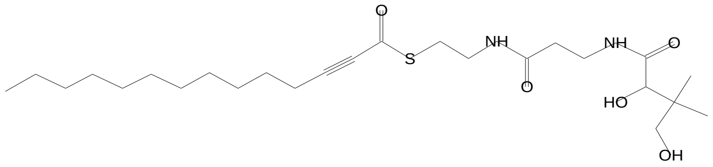


SI 697, RSI 707, mainlib, Entry# 4545, CAS# NA, 2-Myristynoyl pantetheine

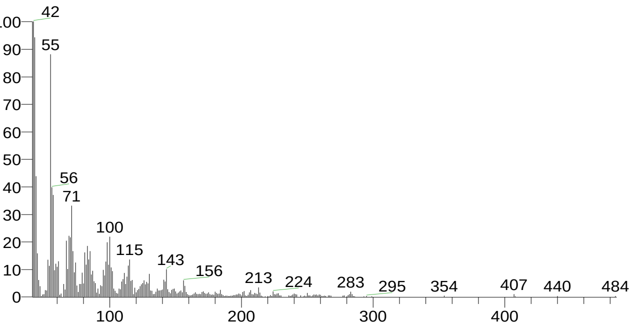


7-Methyl-Z-tetradecen-1-ol acetate
Formula C17H32O2, MW 268, CAS# NA, Entry# 7041
(8Z)-7-Methyl-8-tetradecenyl acetate #

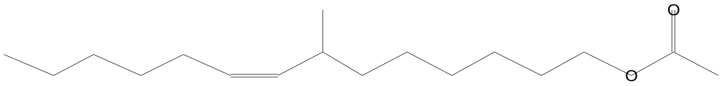


SI 694, RSI 712, mainlib, Entry# 7041, CAS# NA, 7-Methyl-Z-tetradecen-1-ol acetate

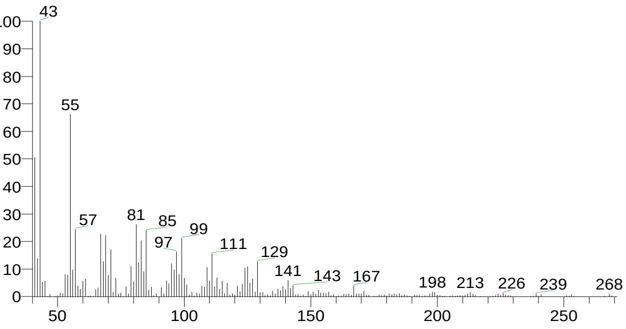


Desulphosinigrin
Formula C10H17NO6S, MW 279, CAS# 5115-81-1, Entry# 30961
1-S-[(1E)-N-Hydroxy-3-butenimidoyl]-1-thiohexopyranose #

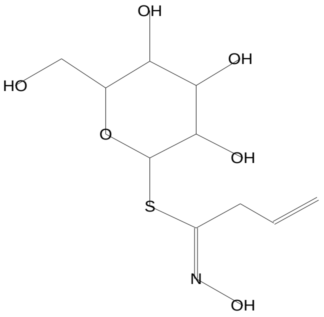


SI 692, RSI 733, mainlib, Entry# 30961, CAS# 5115-81-1, Desulphosinigrin

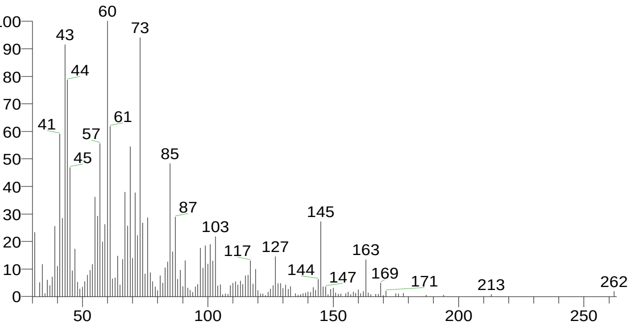


Melezitose
Formula C18H32O16, MW 504, CAS# 597-12-6, Entry# 41139
à-D-Glucopyranoside, O-à-D-glucopyranosyl-(1.fwdarw.3)-á-D-fructofuranosyl

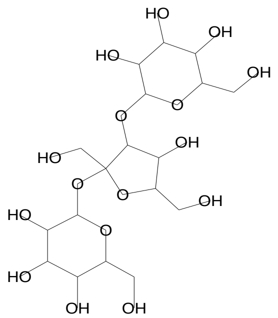


SI 684, RSI 714, mainlib, Entry# 41139, CAS# 597-12-6, Melezitose

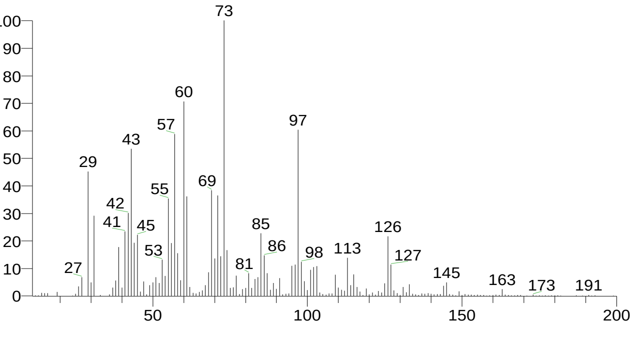

### Library Search Report

Compound Structure

tert-Hexadecanethiol
Formula C16H34S, MW 258, CAS# 25360-09-2, Entry# 25117
1,1-Dimethyltetradecyl hydrosulfide #

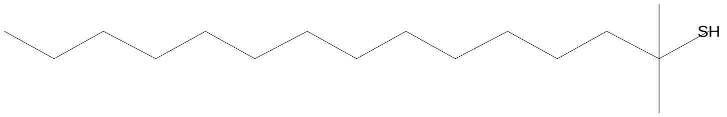


Hit Spectrum

SI 678, RSI 698, mainlib, Entry# 25117, CAS# 25360-09-2, tert-Hexadecanethiol

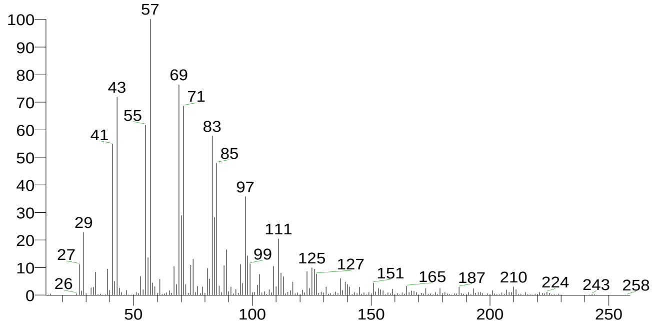


| RT | Compound Name | Area % | Molecular Weight | Molecular Formula | Probability |
|---|---|---|---|---|---|
| 16.48 | 2-Myristynoyl pantetheine | 1.09 | 484 | C25H44N2O5S | 9.38 |
| 16.48 | 7-Methyl-Z-tetradecen-1-ol acetate | 1.09 | 268 | C17H32O2 | 8.28 |
| 16.48 | Desulphosinigrin | 1.09 | 279 | C10H17NO6S | 7.64 |
| 16.48 | Melezitose | 1.09 | 504 | C18H32O16 | 5.70 |
| 16.48 | tert-Hexadecanethiol | 1.09 | 258 | C16H34S | 4.48 |

There is no signature data to report. Anamika Singh

### Library Search Report

#### Lib. Search Graphics Table

| Compound Structure | Hit Spectrum |
|---|---|

Tetradecane, 2,6,10-trimethyl-
Formula C17H36, MW 240, CAS# 14905-56-7, Entry# 24375
2,6,10-Trimethyltetradecane

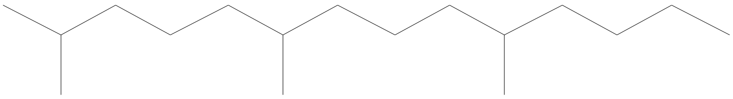

SI 735, RSI 794, mainlib, Entry# 24375, CAS# 14905-56-7, Tetradecane, 2,6,10-trimethyl-

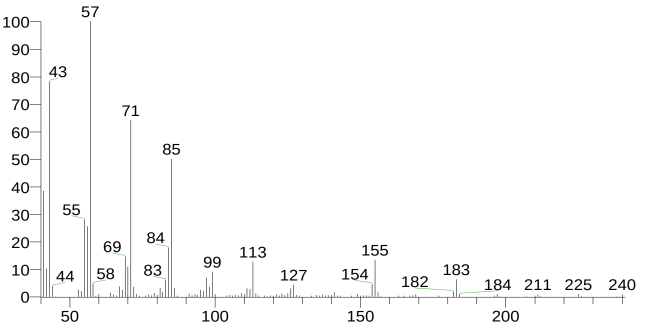


Disulfide, di-tert-dodecyl
Formula C24H50S2, MW 402, CAS# 27458-90-8, Entry# 25353
Di-tert-dodecyl disulfide

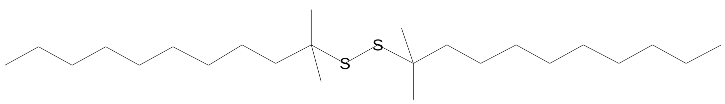


SI 716, RSI 744, mainlib, Entry# 25353, CAS# 27458-90-8, Disulfide, di-tert-dodecyl

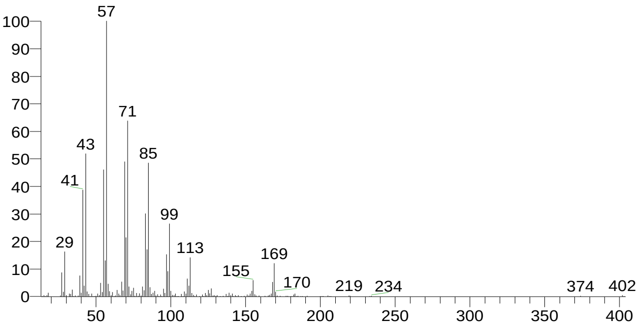


Ethanol, 2-(hexadecyloxy)-
Formula C18H38O2, MW 286, CAS# 2136-71-2, Entry# 6132
2-Hexadecoxyethanol

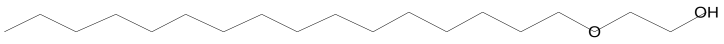


SI 704, RSI 751, replib, Entry# 6132, CAS# 2136-71-2, Ethanol, 2-(hexadecyloxy)-

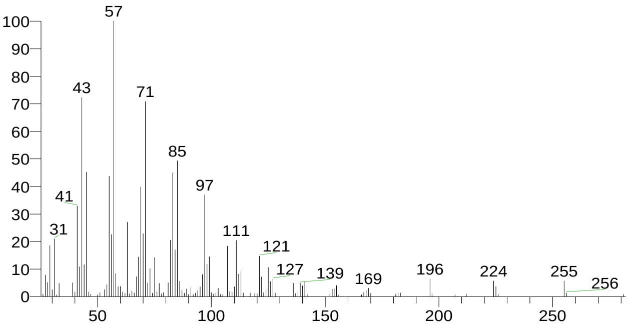


Heptadecane, 2,6,10,15-tetramethyl-
Formula C21H44, MW 296, CAS# 54833-48-6, Entry# 25344
2,6,10,15-Tetramethylheptadecane #

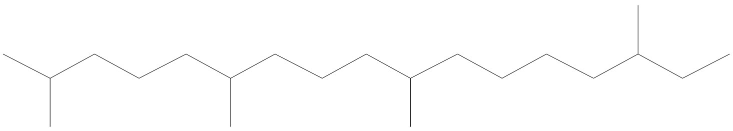

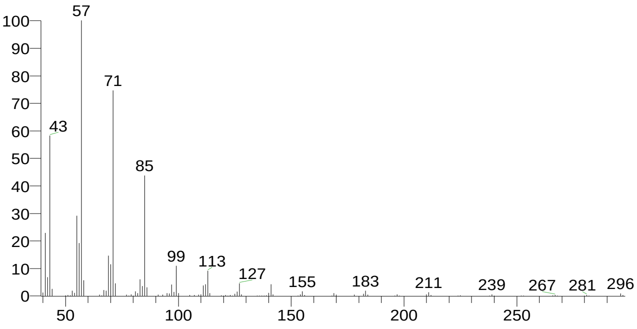


There is no signature data to report. Anamika Singh

### Library Search Report

Compound Structure

Methoxyacetic acid, 2-tetradecyl ester
Formula C17H34O3, MW 286, CAS# NA, Entry# 24635
1-Methyltridecyl methoxyacetate #

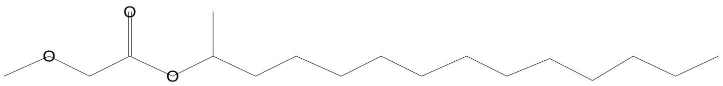

Hit Spectrum

SI 695, RSI 749, mainlib, Entry# 24635, CAS# NA, Methoxyacetic acid, 2-tetradecyl ester

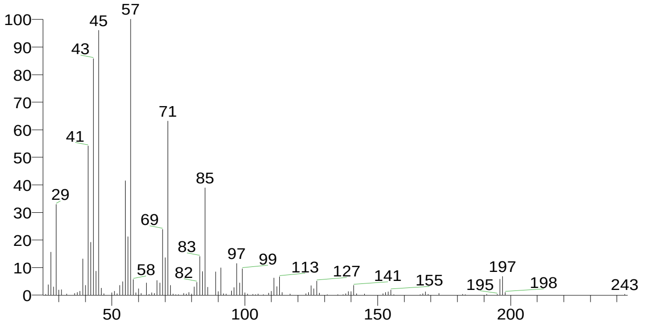


| RT | Compound Name | Area % | Molecular Weight | Molecular Formula | Probability |
|---|---|---|---|---|---|
| 16.59 | Tetradecane, 2,6,10-trimethyl- | 1.47 | 240 | C17H36 | 11.76 |
| 16.59 | Disulfide, di-tert-dodecyl | 1.47 | 402 | C24H50S2 | 5.71 |
| 16.59 | Ethanol, 2-(hexadecyloxy)- | 1.47 | 286 | C18H38O2 | 3.80 |
| 16.59 | Heptadecane, 2,6,10,15-tetramethyl- | 1.47 | 296 | C21H44 | 3.21 |
| 16.59 | Methoxyacetic acid, 2-tetradecyl ester | 1.47 | 286 | C17H34O3 | 2.59 |

There is no signature data to report. Anamika Singh

### Library Search Report

#### Lib. Search Graphics Table

| Compound Structure | Hit Spectrum |
|---|---|

á-D-Glucopyranose, 4-O-á-D-galactopyranosyl-
Formula C12H22O11, MW 342, CAS# 5965-66-2, Entry# 9475
Lactose, á-

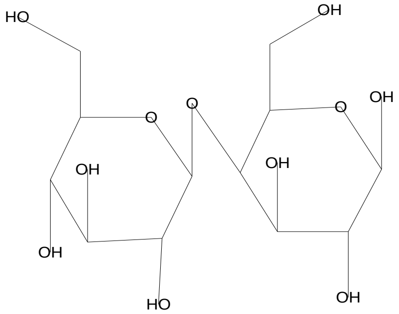


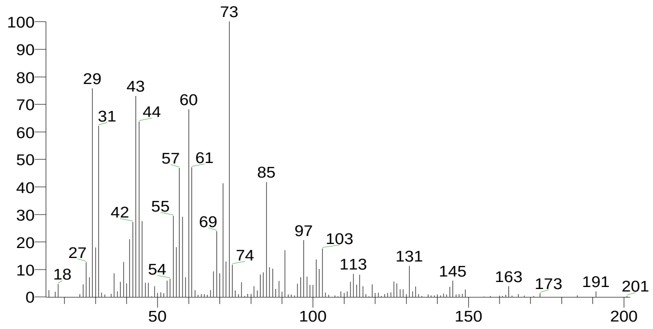


d-Mannose
Formula C6H12O6, MW 180, CAS# 3458-28-4, Entry# 41111
Mannose, d-

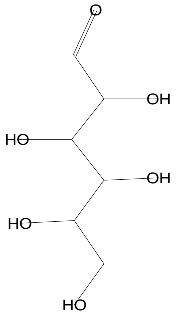


SI 714, RSI 736, mainlib, Entry# 41111, CAS# 3458-28-4, d-Mannose

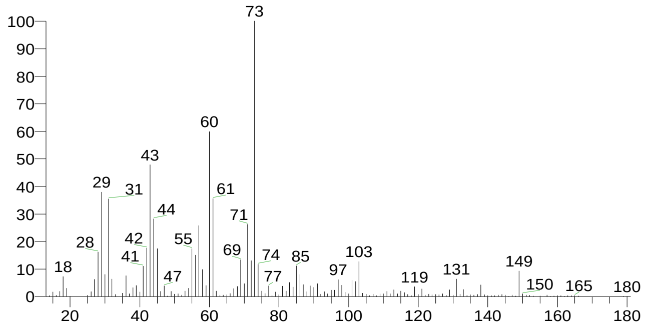


Desulphosinigrin
Formula C10H17NO6S, MW 279, CAS# 5115-81-1, Entry# 30961
1-S-[(1E)-N-Hydroxy-3-butenimidoyl]-1-thiohexopyranose #

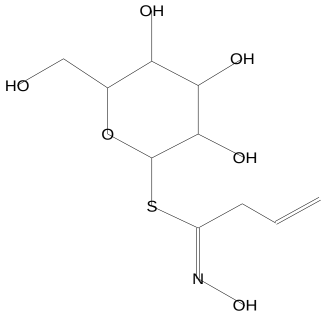


SI 709, RSI 727, mainlib, Entry# 30961, CAS# 5115-81-1, Desulphosinigrin

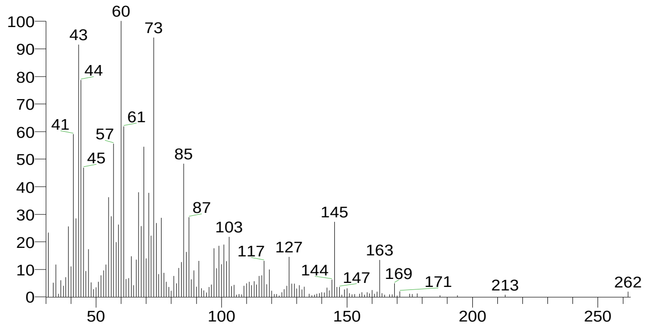


á-D-Glucopyranose, 4-O-á-D-galactopyranosyl-
Formula C12H22O11, MW 342, CAS# 5965-66-2, Entry# 40630
Lactose, á-

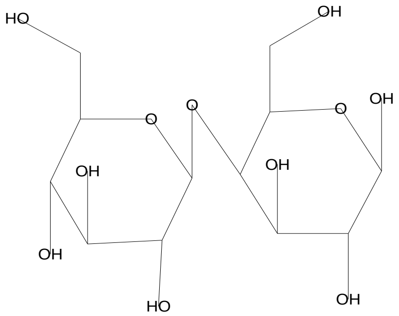


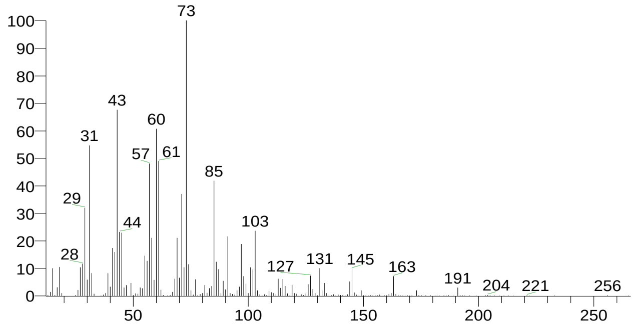


| | |
|---|---|
| There is no signature data to report. | Anamika Singh |

### Library Search Report

Compound Structure

d-Glycero-d-ido-heptose
Formula C7H14O7, MW 210, CAS# NA, Entry# 9206
Heptose #

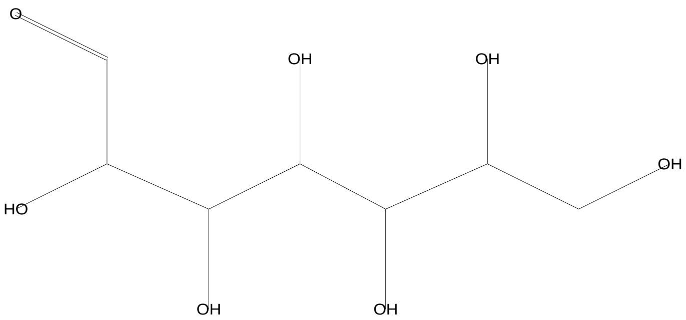


Hit Spectrum

SI 685, RSI 712, mainlib, Entry# 9206, CAS# NA, d-Glycero-d-ido-heptose

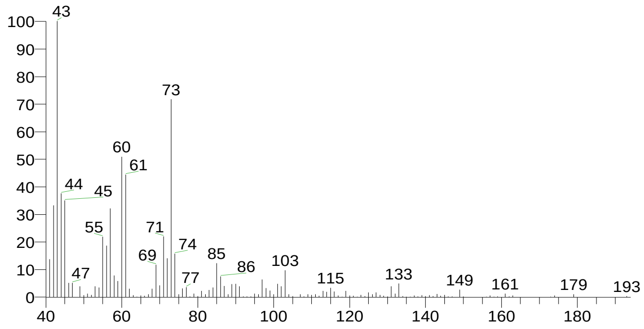


| RT | Compound Name | Area % | Molecular Weight | Molecular Formula | Probability |
|---|---|---|---|---|---|
| 18.34 | á-D-Glucopyranose, 4-O-á-D-galactopyranosyl- | 2.04 | 342 | C12H22O11 | 14.38 |
| 18.34 | á-D-Glucopyranose, 4-O-á-D-galactopyranosyl- | 2.04 | 342 | C12H22O11 | 14.38 |
| 18.34 | d-Mannose | 2.04 | 180 | C6H12O6 | 13.82 |
| 18.34 | Desulphosinigrin | 2.04 | 279 | C10H17NO6S | 11.14 |
| 18.34 | d-Glycero-d-ido-heptose | 2.04 | 210 | C7H14O7 | 3.73 |

There is no signature data to report. Anamika Singh

### Library Search Report

#### Lib. Search Graphics Table

| Compound Structure | Hit Spectrum |
|---|---|
| Ribitol<br>Formula C5H12O5, MW 152, CAS# 488-81-3, Entry# 44657<br>Adonit | SI 769, RSI 834, mainlib, Entry# 44657, CAS# 488-81-3, Ribitol |

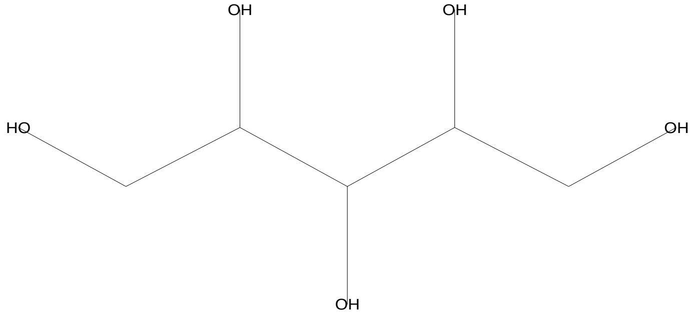


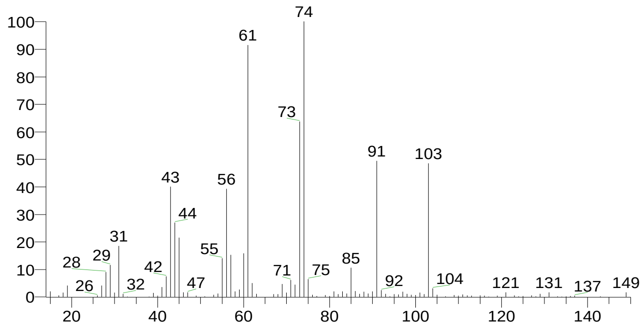


| Compound Structure | Hit Spectrum |
|---|---|
| DL-Arabinitol<br>Formula C5H12O5, MW 152, CAS# 6018-27-5, Entry# 31264<br>Pentitol # | SI 742, RSI 860, mainlib, Entry# 31264, CAS# 6018-27-5, DL-Arabinitol |

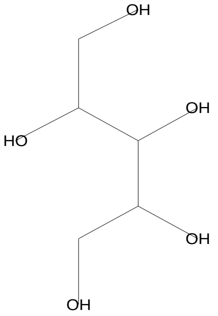


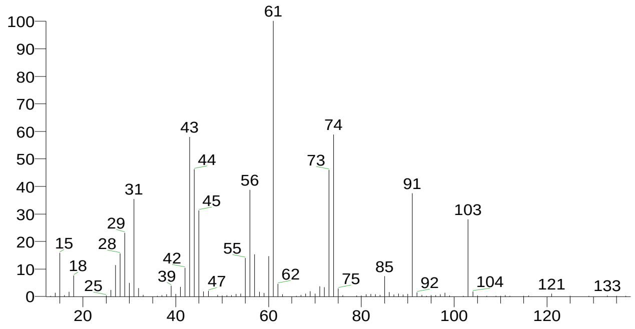


| Compound Structure | Hit Spectrum |
|---|---|
| d-Mannose<br>Formula C6H12O6, MW 180, CAS# 3458-28-4, Entry# 41111<br>Mannose, d- | SI 722, RSI 736, mainlib, Entry# 41111, CAS# 3458-28-4, d-Mannose |

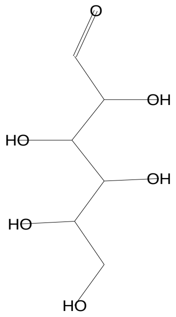


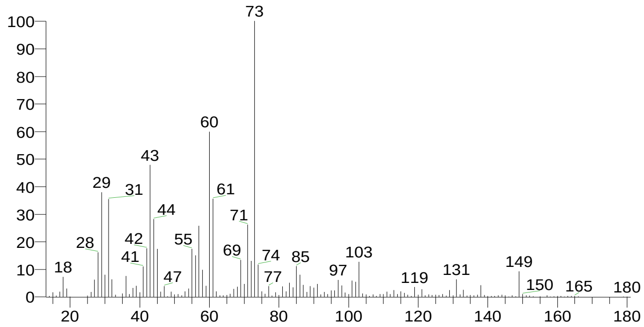


| Compound Structure | Hit Spectrum |
|---|---|
| D-Arabinitol<br>Formula C5H12O5, MW 152, CAS# 488-82-4, Entry# 31170<br>Arabinitol, D- | SI 717, RSI 864, mainlib, Entry# 31170, CAS# 488-82-4, D-Arabinitol |

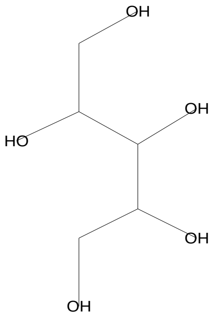


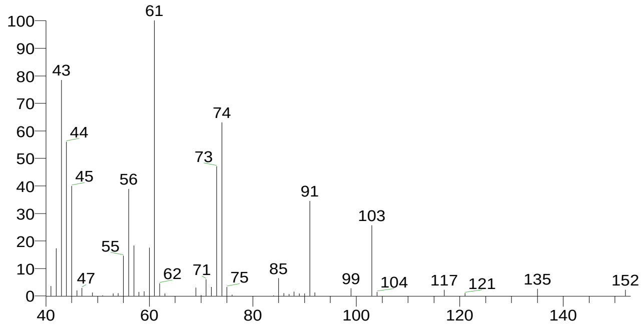

### Library Search Report

Compound Structure

L-Arabinitol
Formula C5H12O5, MW 152, CAS# 7643-75-6, Entry# 7608
Arabinitol, L-

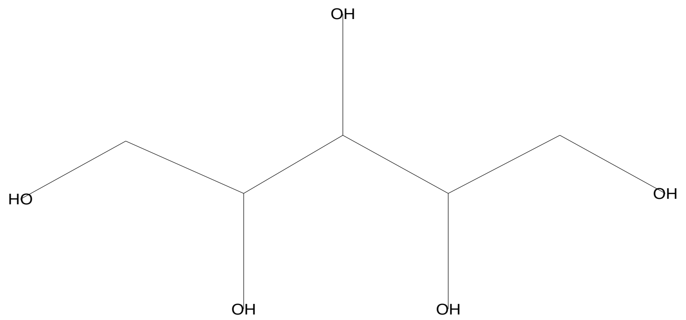


Hit Spectrum

SI 714, RSI 812, replib, Entry# 7608, CAS# 7643-75-6, L-Arabinitol

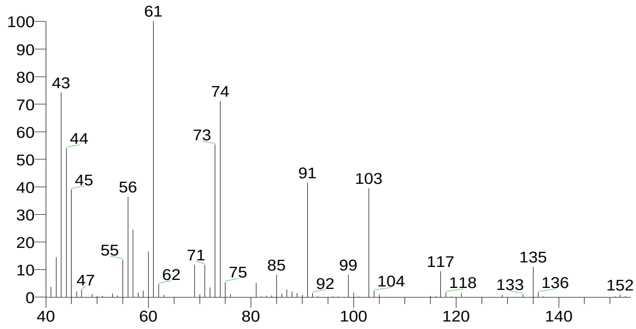


| RT | Compound Name | Area % | Molecular Weight | Molecular Formula | Probability |
|---|---|---|---|---|---|
| 18.52 | Ribitol | 2.39 | 152 | C5H12O5 | 45.37 |
| 18.52 | DL-Arabinitol | 2.39 | 152 | C5H12O5 | 13.34 |
| 18.52 | d-Mannose | 2.39 | 180 | C6H12O6 | 6.08 |
| 18.52 | D-Arabinitol | 2.39 | 152 | C5H12O5 | 4.90 |
| 18.52 | L-Arabinitol | 2.39 | 152 | C5H12O5 | 4.33 |

### Library Search Report

#### Lib. Search Graphics Table

| Compound Structure | Hit Spectrum |
|---|---|

Ribitol
Formula C5H12O5, MW 152, CAS# 488-81-3, Entry# 44657
Adonit

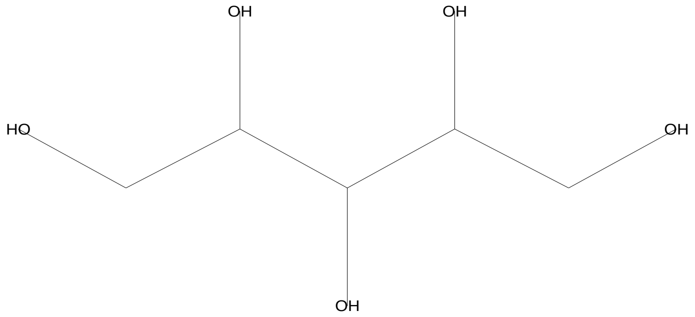


SI 778, RSI 838, mainlib, Entry# 44657, CAS# 488-81-3, Ribitol

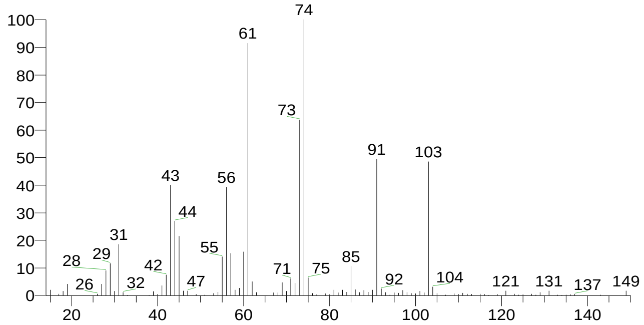


DL-Arabinitol
Formula C5H12O5, MW 152, CAS# 6018-27-5, Entry# 31264
Pentitol #

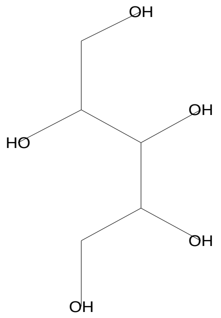


SI 751, RSI 862, mainlib, Entry# 31264, CAS# 6018-27-5, DL-Arabinitol

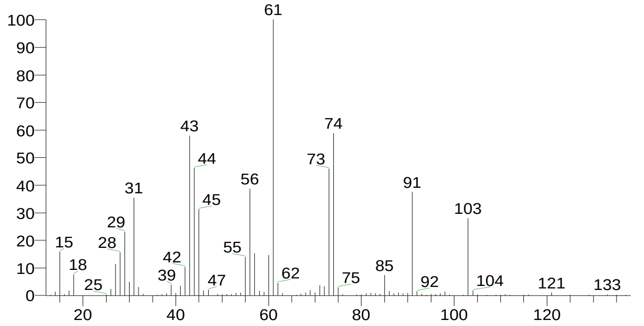


D-Arabinitol
Formula C5H12O5, MW 152, CAS# 488-82-4, Entry# 31170
Arabinitol, D-

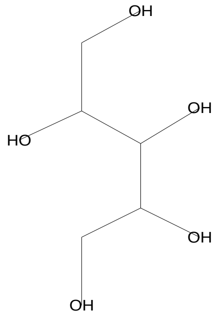


SI 723, RSI 860, mainlib, Entry# 31170, CAS# 488-82-4, D-Arabinitol

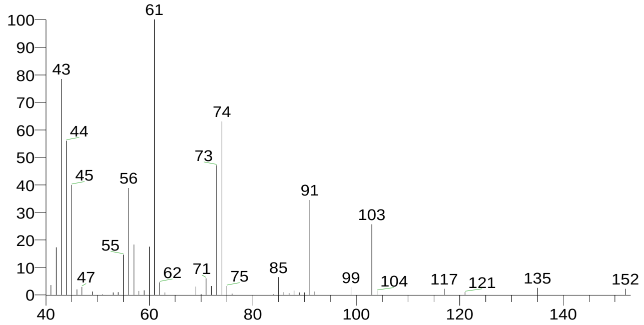


d-Mannose
Formula C6H12O6, MW 180, CAS# 3458-28-4, Entry# 41111
Mannose, d-

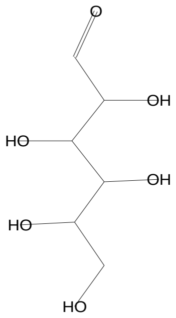


SI 722, RSI 734, mainlib, Entry# 41111, CAS# 3458-28-4, d-Mannose

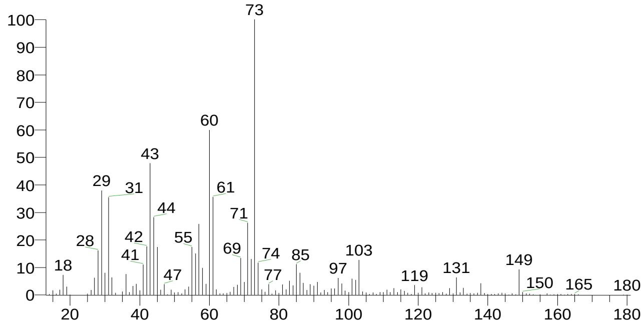


There is no signature data to report. Anamika Singh

### Library Search Report

Compound Structure

L-Arabinitol
Formula C5H12O5, MW 152, CAS# 7643-75-6, Entry# 7608
Arabinitol, L-

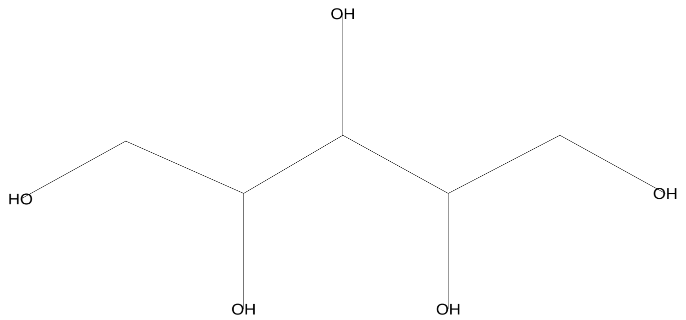


Hit Spectrum

SI 719, RSI 807, replib, Entry# 7608, CAS# 7643-75-6, L-Arabinitol

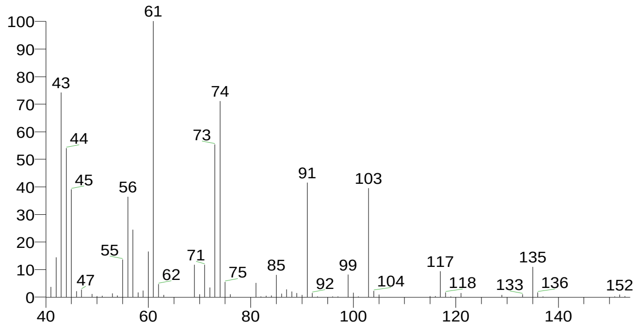


| RT | Compound Name | Area % | Molecular Weight | Molecular Formula | Probability |
|---|---|---|---|---|---|
| 18.61 | Ribitol | 2.40 | 152 | C5H12O5 | 50.47 |
| 18.61 | DL-Arabinitol | 2.40 | 152 | C5H12O5 | 14.84 |
| 18.61 | D-Arabinitol | 2.40 | 152 | C5H12O5 | 4.28 |
| 18.61 | d-Mannose | 2.40 | 180 | C6H12O6 | 4.12 |
| 18.61 | L-Arabinitol | 2.40 | 152 | C5H12O5 | 3.64 |

### Library Search Report

#### Lib. Search Graphics Table

| Compound Structure | Hit Spectrum |
| --- | --- |

Ribitol
Formula C5H12O5, MW 152, CAS# 488-81-3, Entry# 44657
Adonit

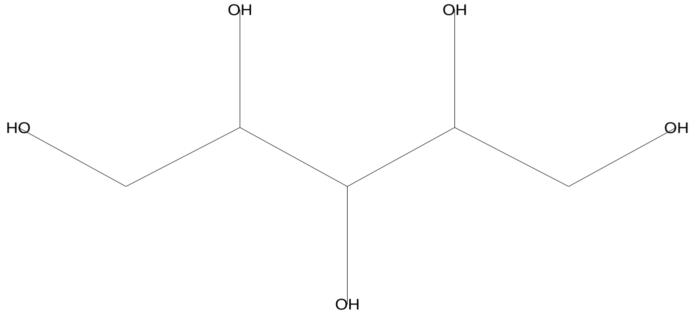


SI 795, RSI 853, mainlib, Entry# 44657, CAS# 488-81-3, Ribitol

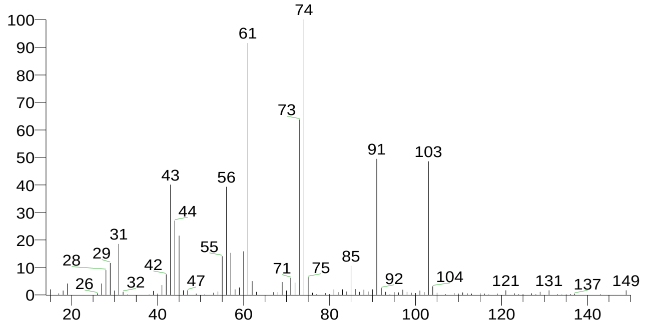


DL-Arabinitol
Formula C5H12O5, MW 152, CAS# 6018-27-5, Entry# 31264
Pentitol #

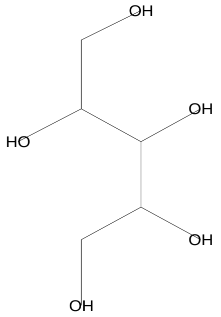


SI 771, RSI 875, mainlib, Entry# 31264, CAS# 6018-27-5, DL-Arabinitol

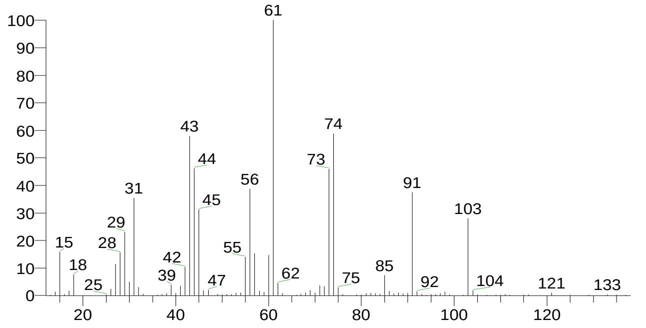


D-Arabinitol
Formula C5H12O5, MW 152, CAS# 488-82-4, Entry# 31170
Arabinitol, D-

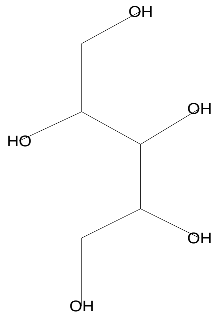


SI 731, RSI 859, mainlib, Entry# 31170, CAS# 488-82-4, D-Arabinitol

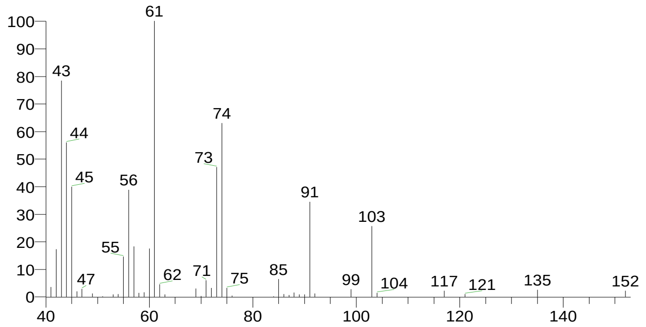


Xylitol
Formula C5H12O5, MW 152, CAS# 87-99-0, Entry# 7603
Xylite

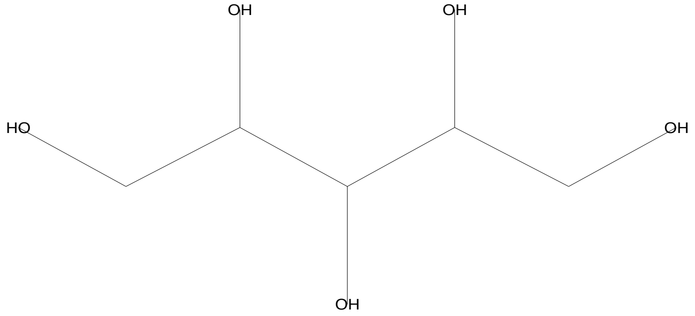


SI 728, RSI 829, replib, Entry# 7603, CAS# 87-99-0, Xylitol

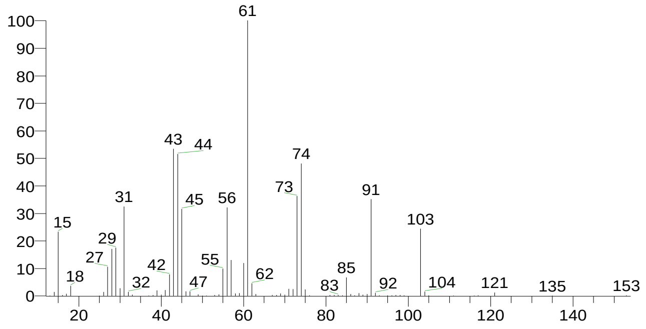


There is no signature data to report. Anamika Singh

### Library Search Report

Compound Structure

L-Arabinitol
Formula C5H12O5, MW 152, CAS# 7643-75-6, Entry# 7608
Arabinitol, L-

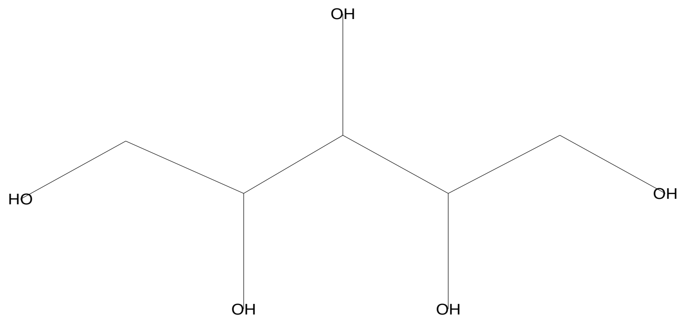


Hit Spectrum

SI 726, RSI 808, replib, Entry# 7608, CAS# 7643-75-6, L-Arabinitol

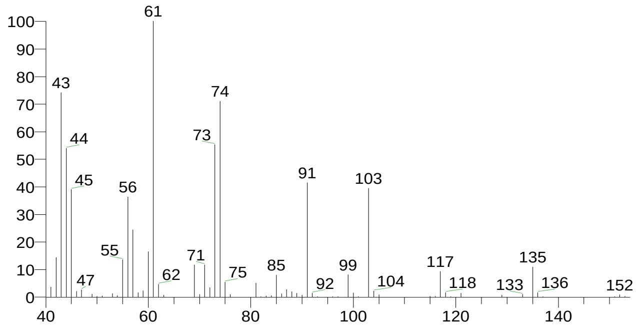


| RT | Compound Name | Area % | Molecular Weight | Molecular Formula | Probability |
|---|---|---|---|---|---|
| 18.71 | Ribitol | 1.97 | 152 | C5H12O5 | 50.78 |
| 18.71 | DL-Arabinitol | 1.97 | 152 | C5H12O5 | 17.01 |
| 18.71 | D-Arabinitol | 1.97 | 152 | C5H12O5 | 3.94 |
| 18.71 | Xylitol | 1.97 | 152 | C5H12O5 | 3.48 |
| 18.71 | L-Arabinitol | 1.97 | 152 | C5H12O5 | 3.21 |

There is no signature data to report. Anamika Singh

### Library Search Report

#### Lib. Search Graphics Table

| Compound Structure | Hit Spectrum |
|---|---|

Ribitol
Formula C5H12O5, MW 152, CAS# 488-81-3, Entry# 44657
Adonit

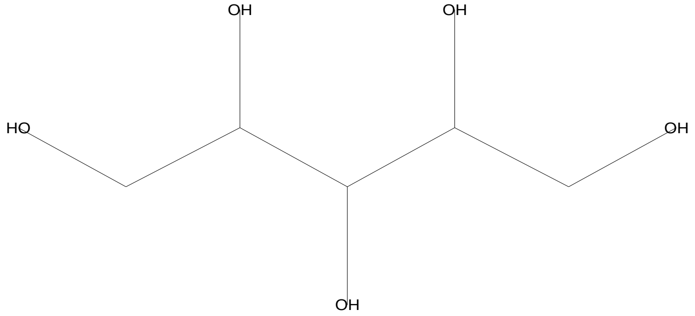


SI 811, RSI 865, mainlib, Entry# 44657, CAS# 488-81-3, Ribitol

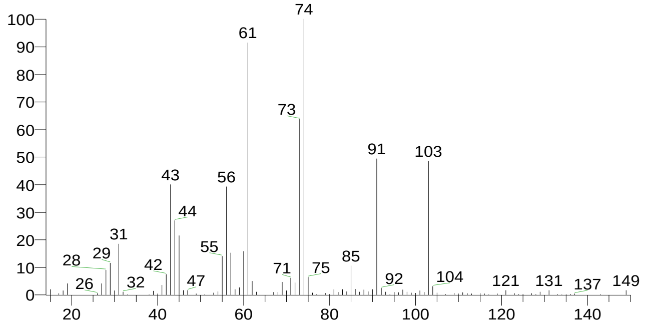


DL-Arabinitol
Formula C5H12O5, MW 152, CAS# 6018-27-5, Entry# 31264
Pentitol #

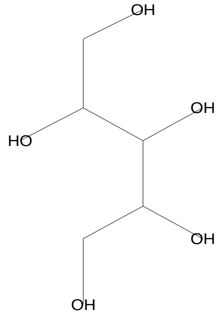


SI 774, RSI 870, mainlib, Entry# 31264, CAS# 6018-27-5, DL-Arabinitol

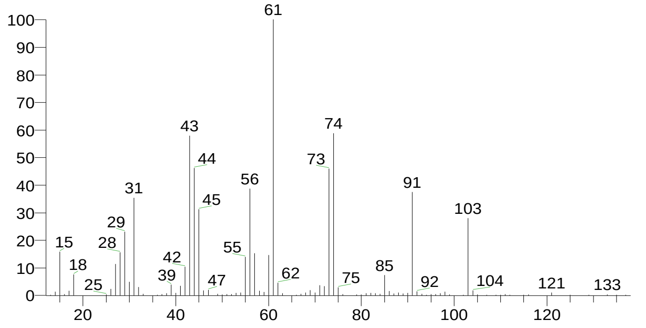


D-Arabinitol
Formula C5H12O5, MW 152, CAS# 488-82-4, Entry# 31170
Arabinitol, D-

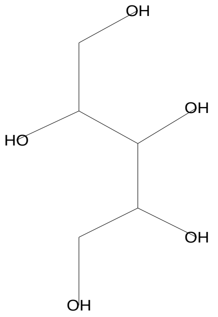


SI 750, RSI 870, mainlib, Entry# 31170, CAS# 488-82-4, D-Arabinitol

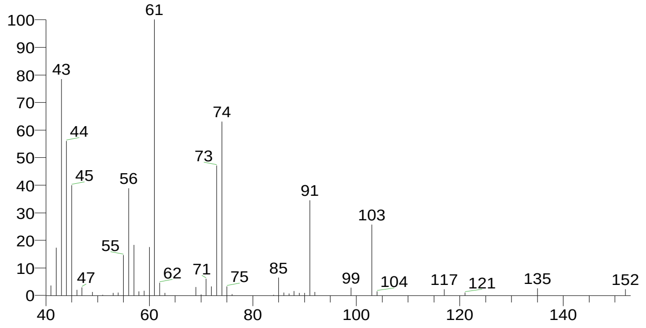


L-Arabinitol
Formula C5H12O5, MW 152, CAS# 7643-75-6, Entry# 7608
Arabinitol, L-

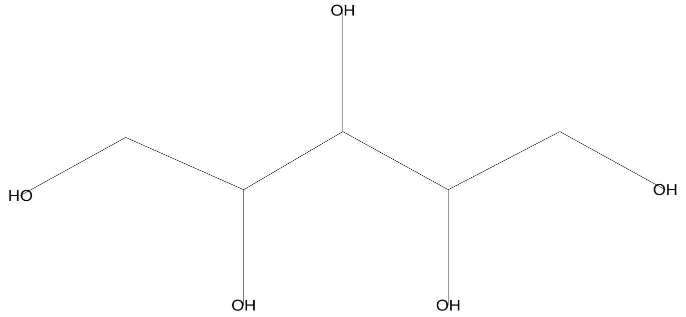


SI 735, RSI 811, replib, Entry# 7608, CAS# 7643-75-6, L-Arabinitol

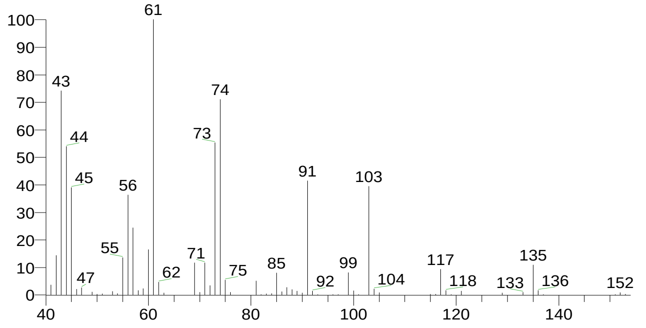


There is no signature data to report. Anamika Singh

### Library Search Report

| Compound Structure | Hit Spectrum |
|---|---|
| Pentane-1,2,3,4,5-pentaol<br>Formula C5H12O5, MW 152, CAS# 6917-36-8, Entry# 31265<br>d-Arabinitol # | SI 734, RSI 863, mainlib, Entry# 31265, CAS# 6917-36-8, Pentane-1,2,3,4,5-pentaol |

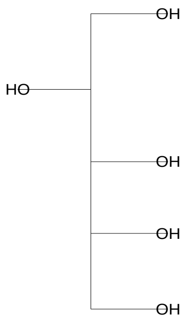


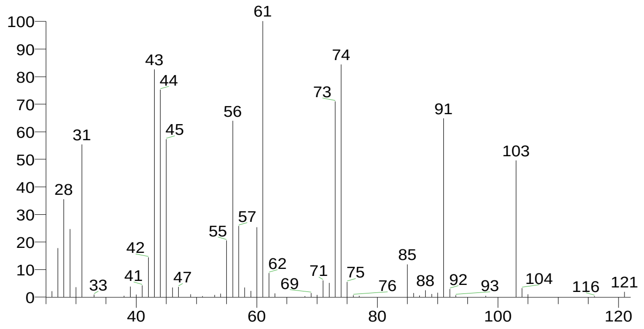


| RT | Compound Name | Area % | Molecular Weight | Molecular Formula | Probability |
|---|---|---|---|---|---|
| 18.79 | Ribitol | 3.48 | 152 | C5H12O5 | 59.86 |
| 18.79 | DL-Arabinitol | 3.48 | 152 | C5H12O5 | 14.57 |
| 18.79 | D-Arabinitol | 3.48 | 152 | C5H12O5 | 4.88 |
| 18.79 | L-Arabinitol | 3.48 | 152 | C5H12O5 | 2.95 |
| 18.79 | Pentane-1,2,3,4,5-pentaol | 3.48 | 152 | C5H12O5 | 2.84 |

### Library Search Report

#### Lib. Search Graphics Table

| Compound Structure | Hit Spectrum |
|---|---|

d-Mannose
Formula C6H12O6, MW 180, CAS# 3458-28-4, Entry# 41111
Mannose, d-

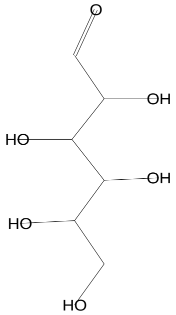

SI 751, RSI 776, mainlib, Entry# 41111, CAS# 3458-28-4, d-Mannose

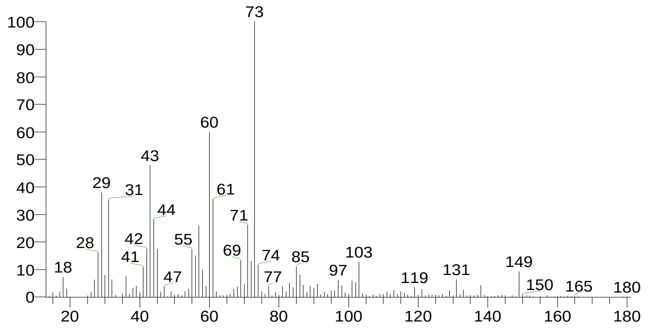


á-D-Glucopyranose, 4-O-á-D-galactopyranosyl-
Formula C12H22O11, MW 342, CAS# 5965-66-2, Entry# 9475
Lactose, á-

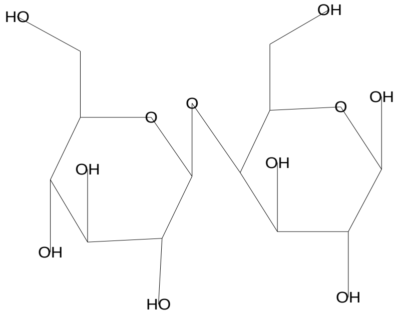

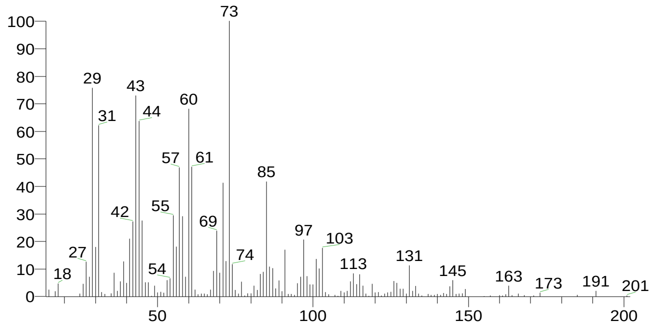


Desulphosinigrin
Formula C10H17NO6S, MW 279, CAS# 5115-81-1, Entry# 30961
1-S-[(1E)-N-Hydroxy-3-butenimidoyl]-1-thiohexopyranose #

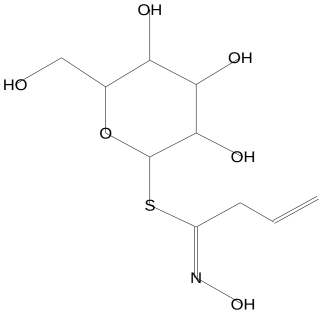

SI 730, RSI 751, mainlib, Entry# 30961, CAS# 5115-81-1, Desulphosinigrin

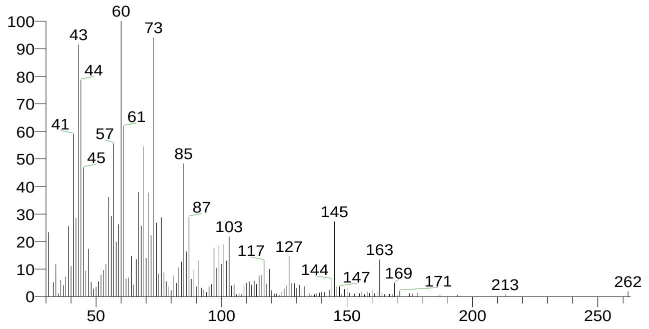


á-D-Glucopyranose, 4-O-á-D-galactopyranosyl-
Formula C12H22O11, MW 342, CAS# 5965-66-2, Entry# 40630
Lactose, á-

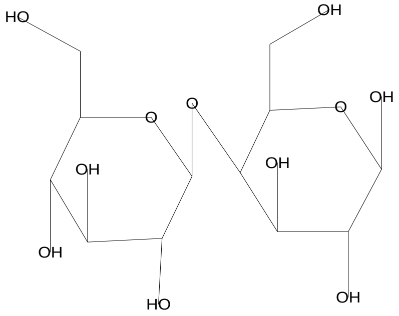

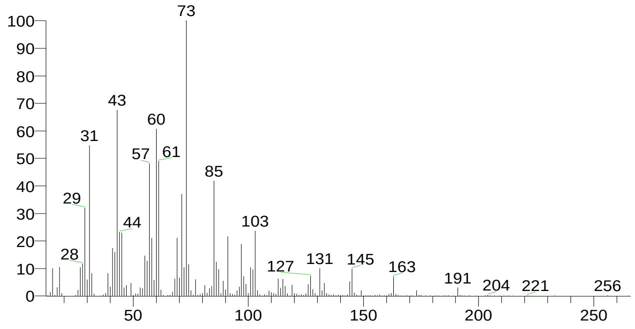


There is no signature data to report. Anamika Singh

### Library Search Report

| Compound Structure | Hit Spectrum |
|---|---|
| L-Glucose<br>Formula C6H12O6, MW 180, CAS# 921-60-8, Entry# 40628<br>Hexopyranose # | SI 723, RSI 752, mainlib, Entry# 40628, CAS# 921-60-8, L-Glucose |

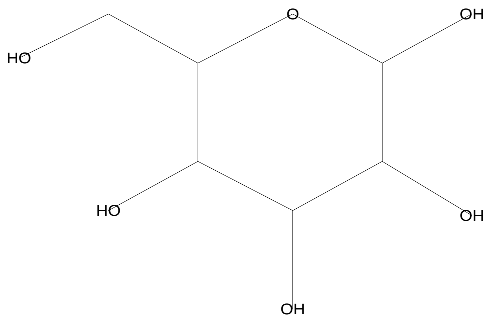


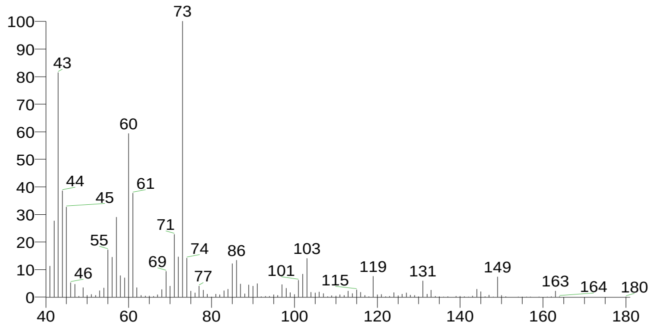


| RT | Compound Name | Area % | Molecular Weight | Molecular Formula | Probability |
|---|---|---|---|---|---|
| 19.17 | d-Mannose | 4.22 | 180 | C6H12O6 | 14.38 |
| 19.17 | á-D-Glucopyranose, 4-O-á-D-galactopyranosyl- | 4.22 | 342 | C12H22O11 | 9.58 |
| 19.17 | á-D-Glucopyranose, 4-O-á-D-galactopyranosyl- | 4.22 | 342 | C12H22O11 | 9.58 |
| 19.17 | Desulphosinigrin | 4.22 | 279 | C10H17NO6S | 6.95 |
| 19.17 | L-Glucose | 4.22 | 180 | C6H12O6 | 5.32 |

There is no signature data to report. Anamika Singh

### Library Search Report

#### Lib. Search Graphics Table

| Compound Structure | Hit Spectrum |
|---|---|

Hexadecanoic acid, methyl ester
Formula C17H34O2, MW 270, CAS# 112-39-0, Entry# 44729
Palmitic acid, methyl ester

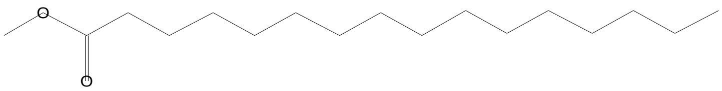


SI 842, RSI 864, mainlib, Entry# 44729, CAS# 112-39-0, Hexadecanoic acid, methyl ester

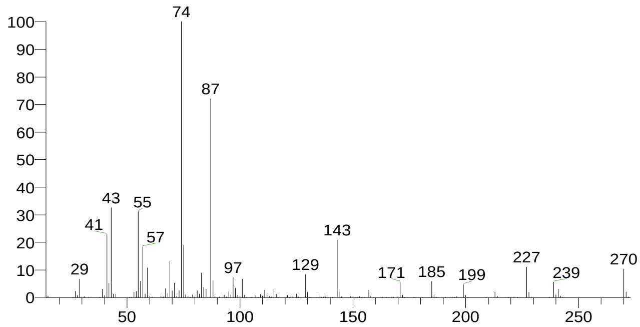


Hexadecanoic acid, methyl ester
Formula C17H34O2, MW 270, CAS# 112-39-0, Entry# 10413
Palmitic acid, methyl ester

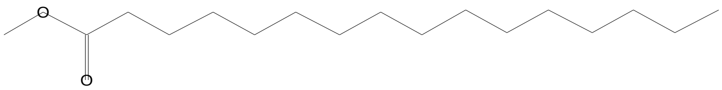


SI 833, RSI 868, replib, Entry# 10413, CAS# 112-39-0, Hexadecanoic acid, methyl ester

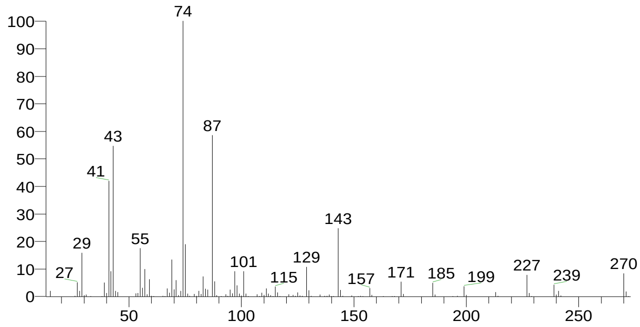


Hexadecanoic acid, methyl ester
Formula C17H34O2, MW 270, CAS# 112-39-0, Entry# 10412
Palmitic acid, methyl ester

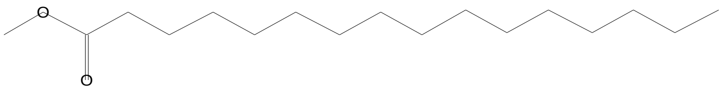


SI 828, RSI 902, replib, Entry# 10412, CAS# 112-39-0, Hexadecanoic acid, methyl ester

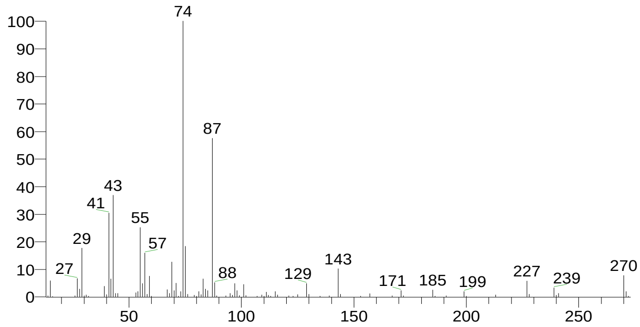


Hexadecanoic acid, methyl ester
Formula C17H34O2, MW 270, CAS# 112-39-0, Entry# 10416
Palmitic acid, methyl ester

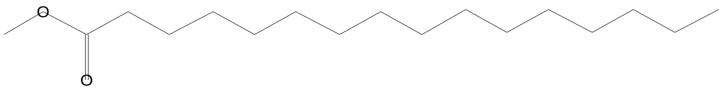


SI 822, RSI 846, replib, Entry# 10416, CAS# 112-39-0, Hexadecanoic acid, methyl ester

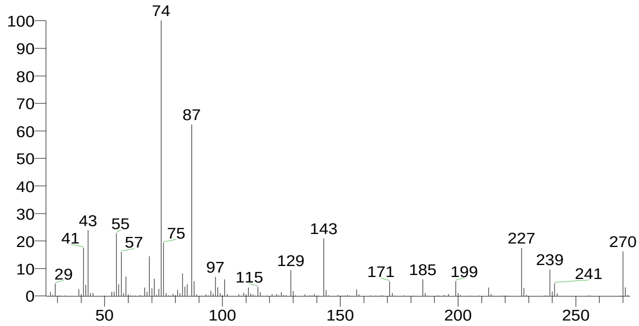


There is no signature data to report. Anamika Singh

### Library Search Report

| Compound Structure | Hit Spectrum |
|---|---|
| Hexadecanoic acid, methyl ester<br>Formula C17H34O2, MW 270, CAS# 112-39-0, Entry# 10414<br>Palmitic acid, methyl ester | SI 820, RSI 855, replib, Entry# 10414, CAS# 112-39-0, Hexadecanoic acid, methyl ester |

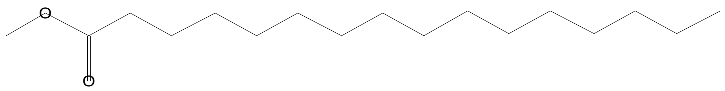


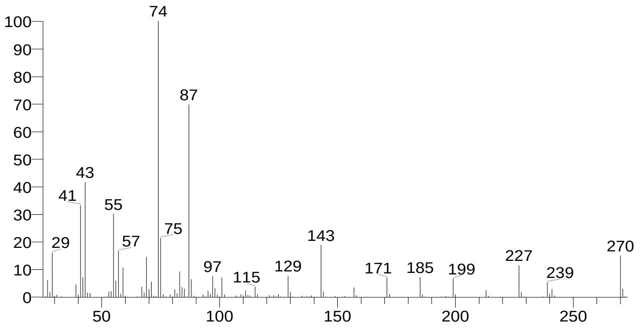


| RT | Compound Name | Area % | Molecular Weight | Molecular Formula | Probability |
|---|---|---|---|---|---|
| 23.34 | Hexadecanoic acid, methyl ester | 4.15 | 270 | C17H34O2 | 64.25 |
| 23.34 | Hexadecanoic acid, methyl ester | 4.15 | 270 | C17H34O2 | 64.25 |
| 23.34 | Hexadecanoic acid, methyl ester | 4.15 | 270 | C17H34O2 | 64.25 |
| 23.34 | Hexadecanoic acid, methyl ester | 4.15 | 270 | C17H34O2 | 64.25 |
| 23.34 | Hexadecanoic acid, methyl ester | 4.15 | 270 | C17H34O2 | 64.25 |

There is no signature data to report. Anamika Singh

### Library Search Report

#### Lib. Search Graphics Table

| Compound Structure | Hit Spectrum |
| --- | --- |

Estra-1,3,5(10)-trien-17á-ol
Formula C18H24O, MW 256, CAS# 2529-64-8, Entry# 7736
Estra-1,3,5(10)-trien-17-ol, (17á)-

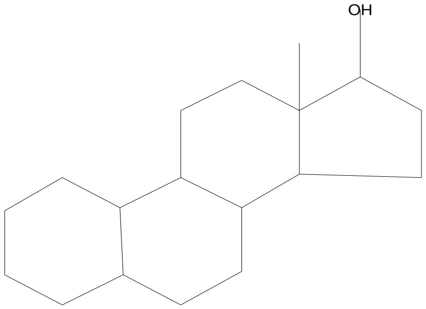


SI 733, RSI 795, mainlib, Entry# 7736, CAS# 2529-64-8, Estra-1,3,5(10)-trien-17á-ol

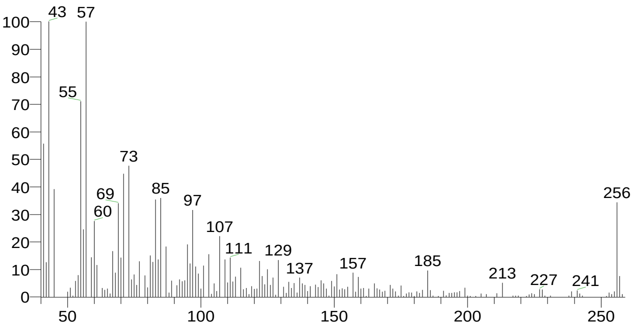


Hexadecanoic acid, 1-(hydroxymethyl)-1,2-ethanediyl ester
Formula C35H68O5, MW 568, CAS# 761-35-3, Entry# 7720
Palmitin, 1,2-di-

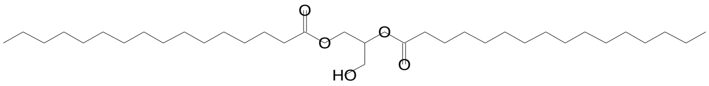


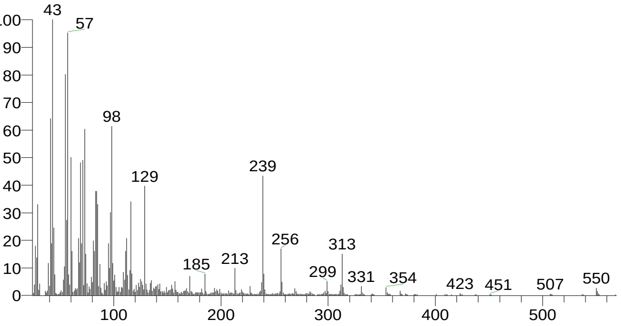


à-D-Glucofuranose, 6-O-(trimethylsilyl)-, cyclic 1,2:3,5-bis(butylboronate)
Formula C17H34B2O6Si, MW 384, CAS# 72347-48-9, Entry# 97049

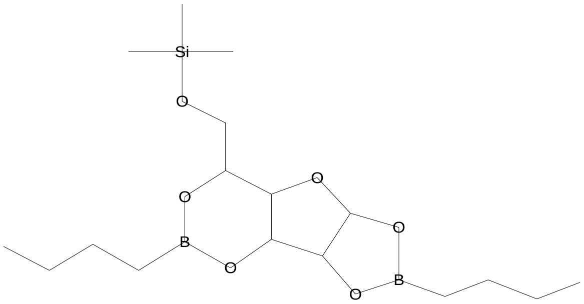


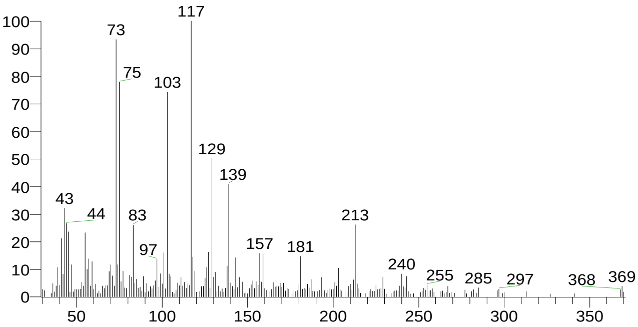


l-(+)-Ascorbic acid 2,6-dihexadecanoate
Formula C38H68O8, MW 652, CAS# 28474-90-0, Entry# 25622
$:28TUYRNAGGIJZRNM-UHFFFAOYSA-N

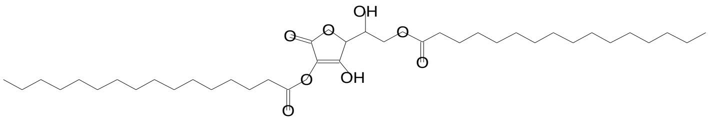


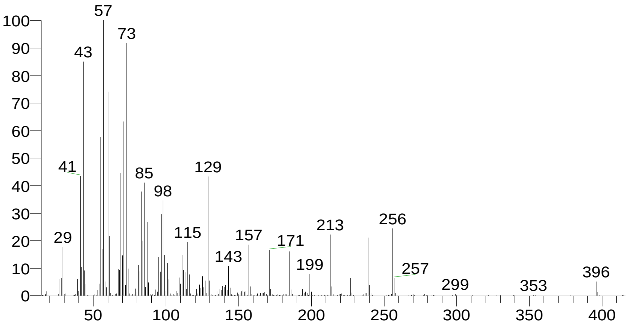

### Library Search Report

Compound Structure

Eicosanoic acid
Formula C20H40O2, MW 312, CAS# 506-30-9, Entry# 7899
Arachic acid

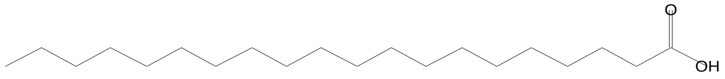


Hit Spectrum

SI 675, RSI 724, mainlib, Entry# 7899, CAS# 506-30-9, Eicosanoic acid

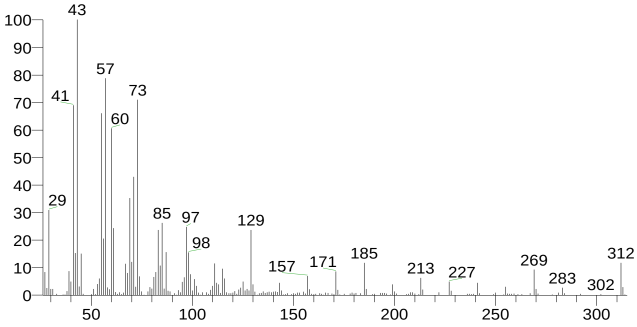


| RT | Compound Name | Area % | Molecular Weight | Molecular Formula | Probability |
|---|---|---|---|---|---|
| 23.87 | Estra-1,3,5(10)-trien-17á-ol | 1.36 | 256 | C18H24O | 38.86 |
| 23.87 | Hexadecanoic acid, 1-(hydroxymethyl)-1,2-ethanediyl ester | 1.36 | 568 | C35H68O5 | 17.70 |
| 23.87 | à-D-Glucofuranose, 6-O-(trimethylsilyl)-, cyclic 1,2:3,5-bis(butylboronate) | 1.36 | 384 | C17H34B2O6Si | 4.92 |
| 23.87 | l-(+)-Ascorbic acid 2,6-dihexadecanoate | 1.36 | 652 | C38H68O8 | 4.15 |
| 23.87 | Eicosanoic acid | 1.36 | 312 | C20H40O2 | 3.51 |

There is no signature data to report. Anamika Singh

### Library Search Report

#### Lib. Search Graphics Table

| Compound Structure | Hit Spectrum |
|---|---|

Heptadecanoic acid, 16-methyl-, methyl ester
Formula C19H38O2, MW 298, CAS# 5129-61-3, Entry# 44788
Methyl isostearate

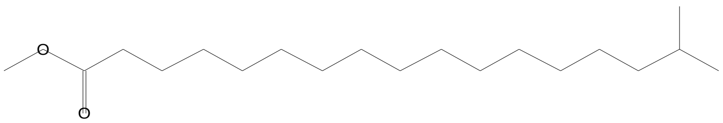


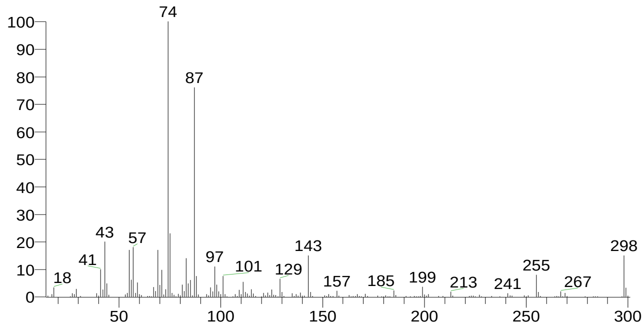


Methyl stearate
Formula C19H38O2, MW 298, CAS# 112-61-8, Entry# 44751
Octadecanoic acid, methyl ester

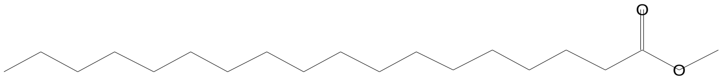


SI 788, RSI 860, mainlib, Entry# 44751, CAS# 112-61-8, Methyl stearate

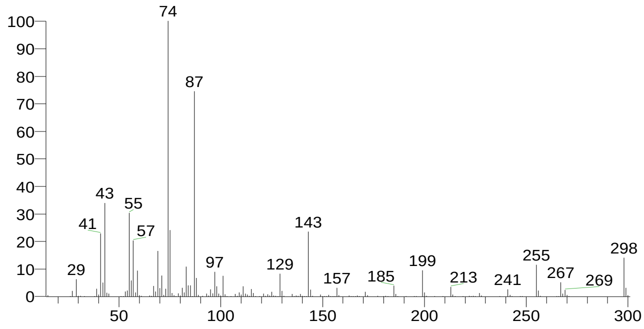


Methyl stearate
Formula C19H38O2, MW 298, CAS# 112-61-8, Entry# 10453
Octadecanoic acid, methyl ester

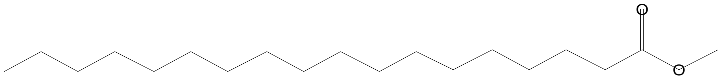


SI 784, RSI 871, replib, Entry# 10453, CAS# 112-61-8, Methyl stearate

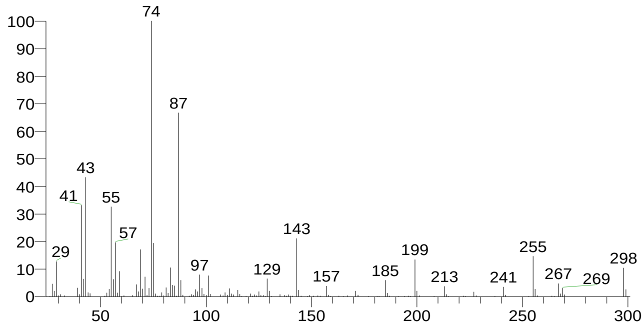


Heptadecanoic acid, 16-methyl-, methyl ester
Formula C19H38O2, MW 298, CAS# 5129-61-3, Entry# 10486
Methyl isostearate

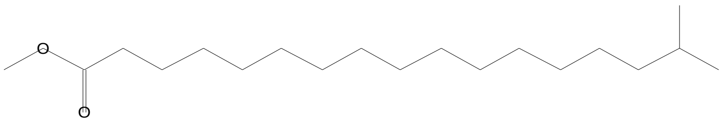


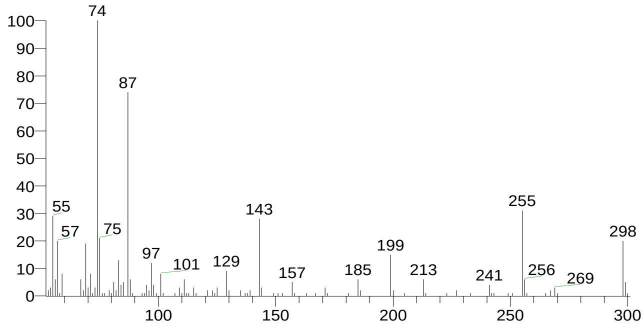


There is no signature data to report. Anamika Singh

### Library Search Report

Compound Structure

Heptadecanoic acid, 9-methyl-, methyl ester
Formula C19H38O2, MW 298, CAS# 54934-57-5, Entry# 44502
Methyl 9-methylheptadecanoate #

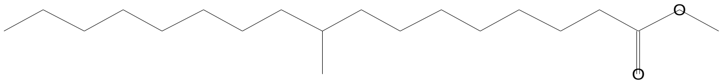


Hit Spectrum

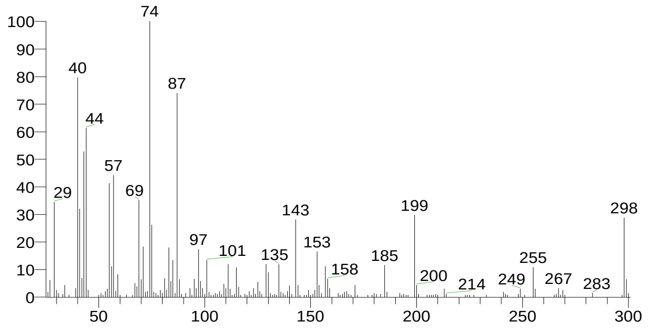


| RT | Compound Name | Area % | Molecular Weight | Molecular Formula | Probability |
|---|---|---|---|---|---|
| 26.06 | Heptadecanoic acid, 16-methyl-, methyl ester | 1.77 | 298 | C19H38O2 | 34.13 |
| 26.06 | Heptadecanoic acid, 16-methyl-, methyl ester | 1.77 | 298 | C19H38O2 | 34.13 |
| 26.06 | Methyl stearate | 1.77 | 298 | C19H38O2 | 20.68 |
| 26.06 | Methyl stearate | 1.77 | 298 | C19H38O2 | 20.68 |
| 26.06 | Heptadecanoic acid, 9-methyl-, methyl ester | 1.77 | 298 | C19H38O2 | 15.01 |
