## supplemental for "Electro-Fermentation of Grape Must via *Candida tropicalis* SY005: Accelerating Kinetics, Modulating Bio-chemical Pathway and Improving Bio-active Content": SU-SG-SS-WC1_0001.pdf

Agilent Technologies

| | | | |
|---|---|---|---|
| Sample ID: | SU-SG-SS-WC1 | Method Name: | shoolini |
| Sample Scans: | 32 | User: | admin |
| Background Scans: | 32 | Date/Time: | 5/17/2024 3:28:40PM |
| Resolution: | 16 cm-1 | Range: | 4,000.00 - 650.00 |
| System Status: | Good | Apodization: | Happ-Genzel |
| File Location: | C:\Program Files\Agilent\MicroLab PC\Results\SU-SG-SS-WC1_0001.a2r | | |

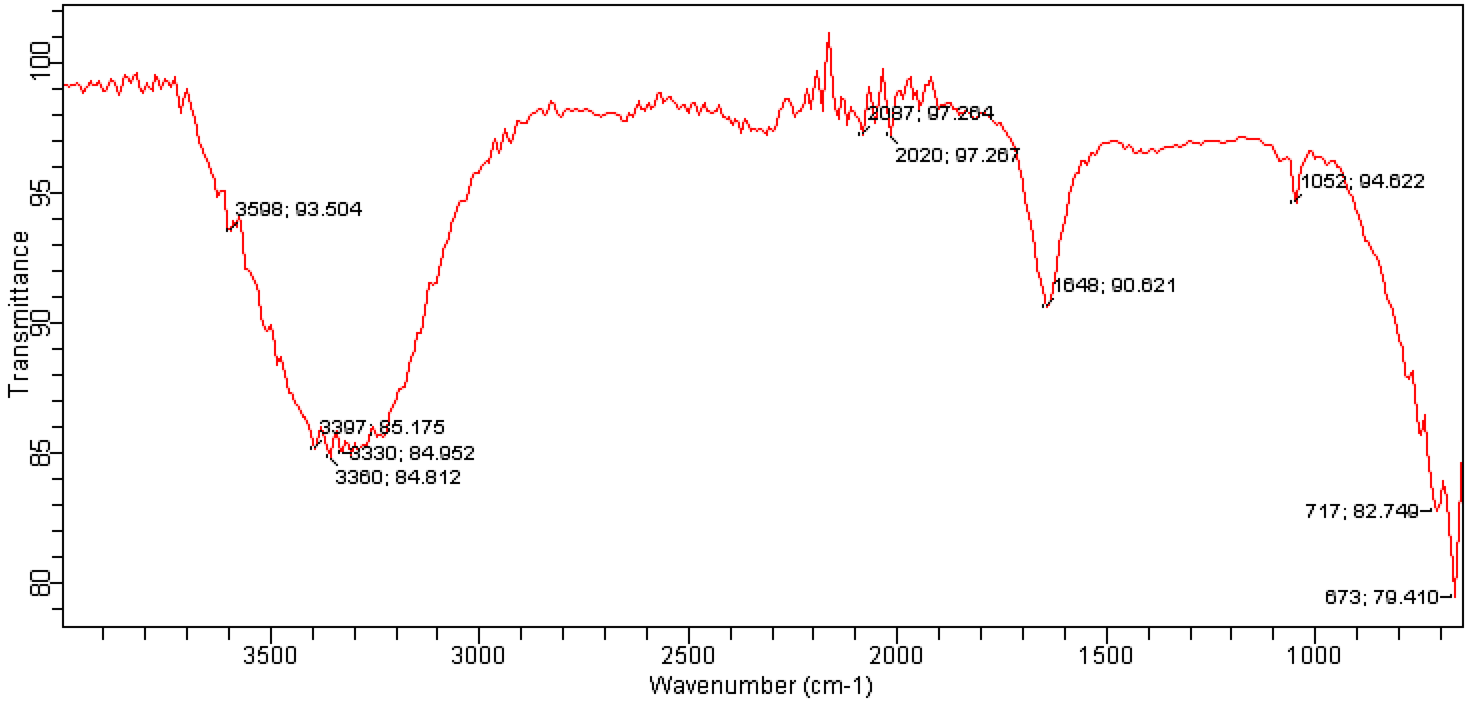
