## supplemental for "Electro-Fermentation of Grape Must via *Candida tropicalis* SY005: Accelerating Kinetics, Modulating Bio-chemical Pathway and Improving Bio-active Content": T4-EDS.pdf

### Smp_Image

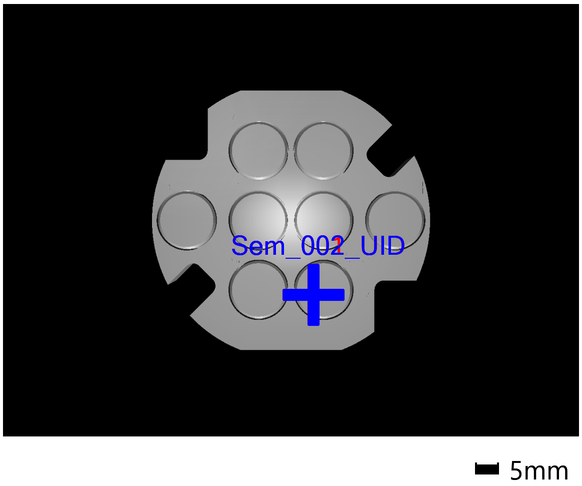


### Sem_001_UID

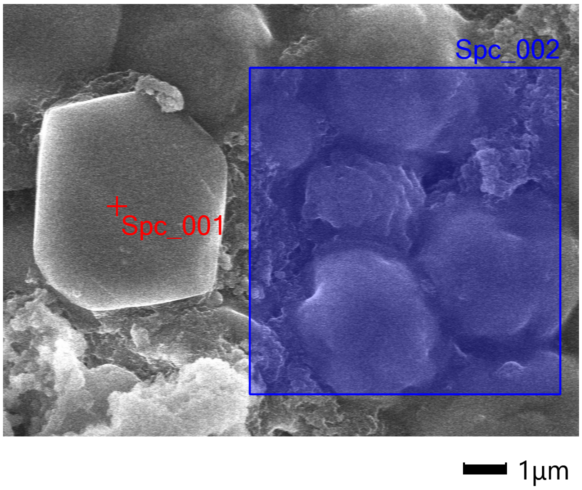


Landing Voltage 10.00 kV
Vacuum Mode HV
Magnification x10.0 k
WD 7.6 mm
Signal UID

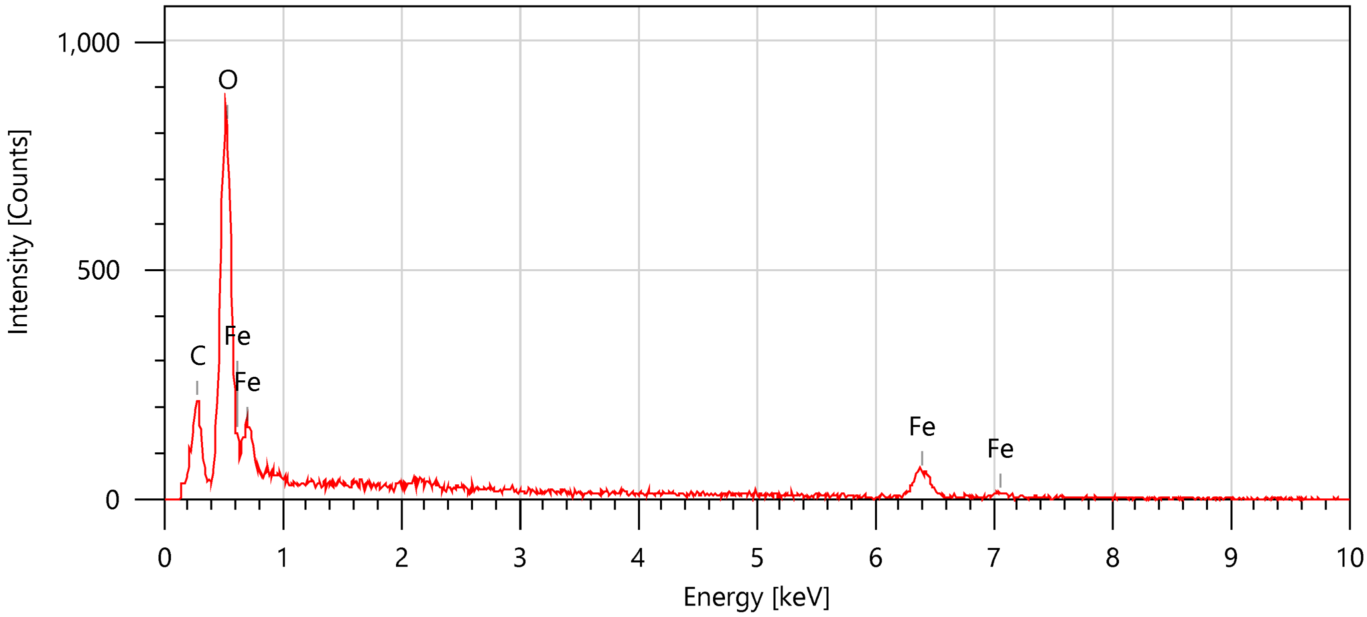


| Items | Value |
|---|---|
| measurement conditions | |
| Acceleration voltage | 10.00 kV |
| Probe current | - |
| Magnification | x 10000 |
| Process time | T1 |
| Measurement detector | First |
| Live time | 30.00 seconds |
| Real time | 31.02 seconds |
| Dead time | 3.00 % |
| Count rate | 858.00 CPS |

| Display name | Standard data | Quantification method | Result Type |
|---|---|---|---|
| Spc_001 | Standardless | ZAF | Metal |

| Element | Line | Mass% | Atom% |
|---|---|---|---|
| C | K | 12.46±0.18 | 23.92±0.34 |
| O | K | 38.79±0.44 | 55.94±0.64 |
| Fe | K | 48.75±1.88 | 20.14±0.78 |
| Total | | 100.00 | 100.00 |
| Spc_001 | | | Fitting ratio 0.1267 |

### Sem_002_UID

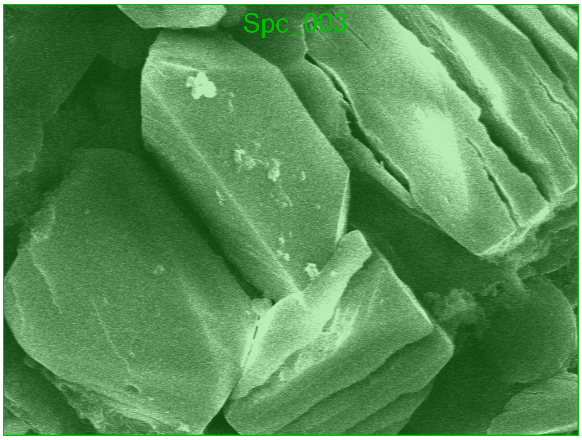


Landing Voltage 10.00 kV
Vacuum Mode HV
Magnification x6.50 k
WD 7.6 mm
Signal UID

1µm

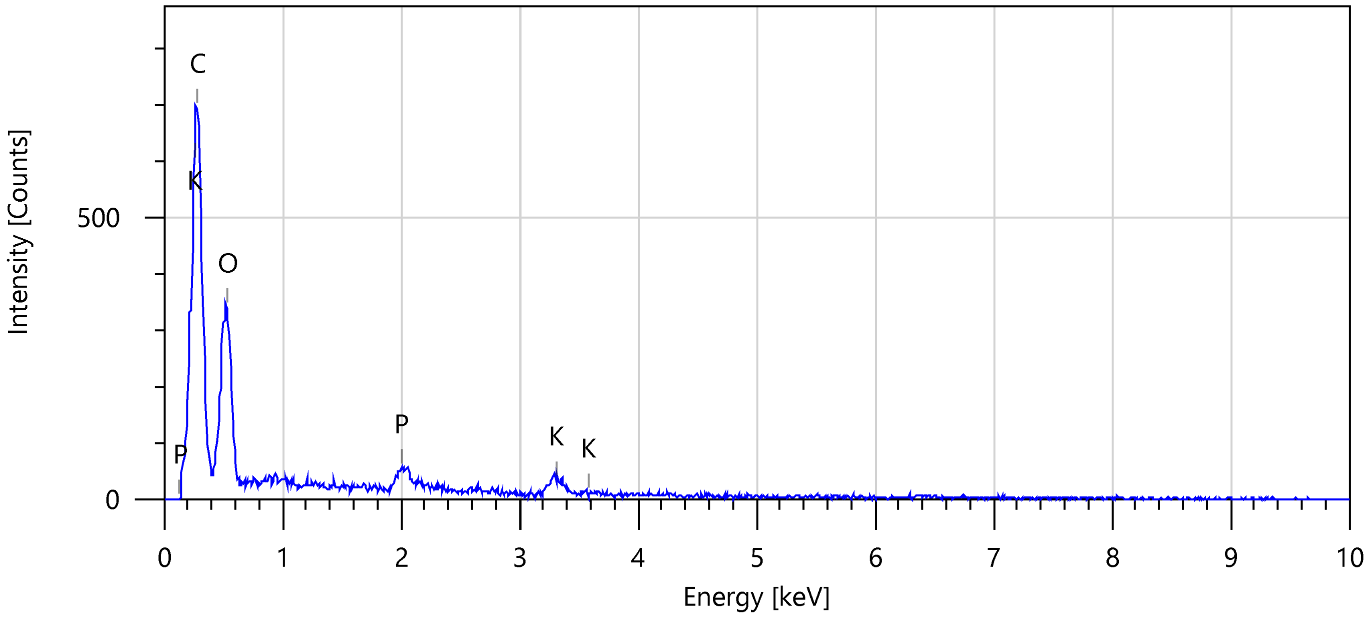


| Items | Value |
|---|---|
| measurement conditions | |
| Acceleration voltage | 10.00 kV |
| Probe current | - |
| Magnification | x 10000 |
| Process time | T1 |
| Measurement detector | First |
| Live time | 30.00 seconds |
| Real time | 32.33 seconds |
| Dead time | 6.00 % |
| Count rate | 559.00 CPS |

| Display name | Standard data | Quantification method | Result Type |
|---|---|---|---|
| Spc_002 | Standardless | ZAF | Metal |

| Element | Line | Mass% | Atom% |
|---|---|---|---|
| C | K | 50.04±0.37 | 58.99±0.44 |
| O | K | 43.39±0.78 | 38.40±0.69 |
| P | K | 2.42±0.21 | 1.11±0.10 |
| K | K | 4.15±0.34 | 1.50±0.12 |
| Total | | 100.00 | 100.00 |
| Spc_002 | | | Fitting ratio 0.1986 |

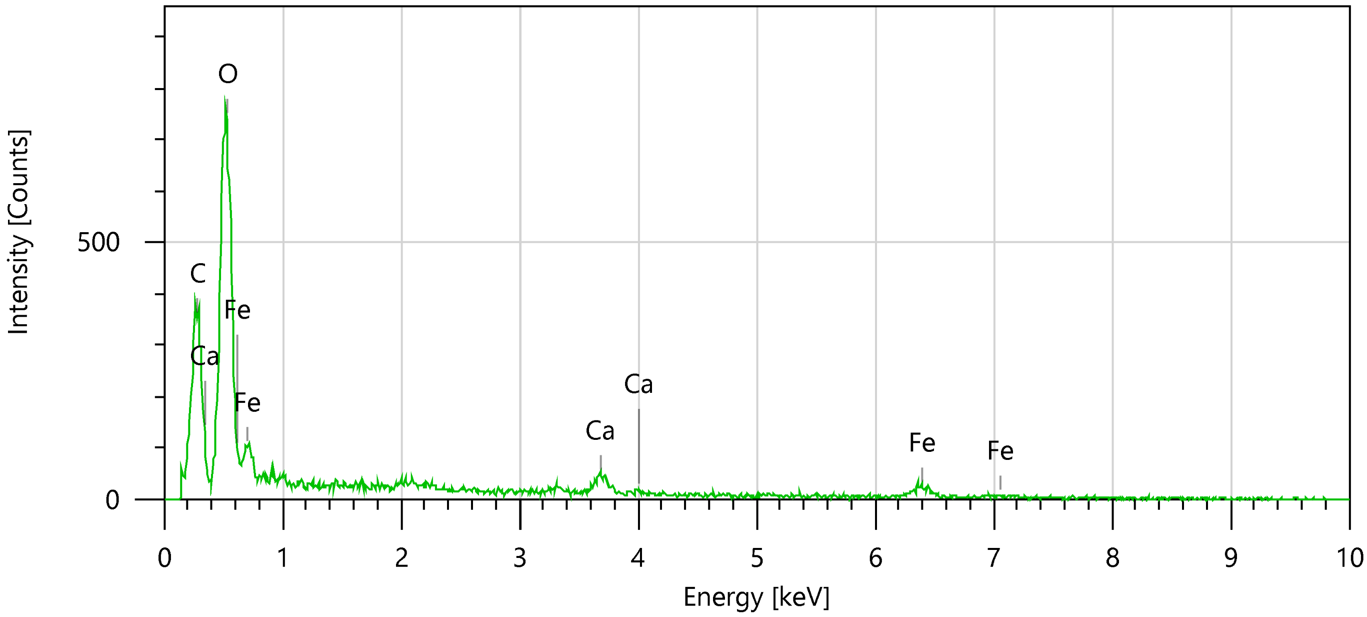


| Items | Value |
|---|---|
| measurement conditions | |
| Acceleration voltage | 10.00 kV |
| Probe current | - |
| Magnification | x 6500 |
| Process time | T1 |
| Measurement detector | First |
| Live time | 30.00 seconds |
| Real time | 31.57 seconds |
| Dead time | 5.00 % |
| Count rate | 718.00 CPS |

| Display name | Standard data | Quantification method | Result Type |
|---|---|---|---|
| Spc_003 | Standardless | ZAF | Metal |

| Element | Line | Mass% | Atom% |
|---|---|---|---|
| C | K | 23.80±0.24 | 34.33±0.35 |
| O | K | 53.64±0.64 | 58.10±0.70 |
| Ca | K | 4.70±0.36 | 2.03±0.15 |
| Fe | K | 17.86±1.35 | 5.54±0.42 |
| Total | | 100.00 | 100.00 |
| Spc_003 | | | Fitting ratio 0.1460 |
